# PWO proteins are associated with PRC2 since their emergence in vascular plants

**DOI:** 10.1101/2025.03.15.643013

**Authors:** Ahamed Khan, Saqlain Haider, Abdoallah Sharaf, Alžbeta Kusová, Jan Skalák, Claire Jourdain, Martin Rennie, Petra Procházková Schrumpfová, Jan Hejátko, Daniel Schubert, Sara Farrona, Iva Mozgová

## Abstract

PWWP-Domain Interactor of Polycombs 1 (PWO1), also known as PWWP1, interacts with the catalytic subunits of the Polycomb Repressive Complex 2 (PRC2) and, together with PWO2/3 proteins, plays a critical role in the development of *Arabidopsis thaliana* (At). PWOs are unique to plants and impact chromatin structure by enabling crosstalk between active and repressive epigenetic marks through mechanisms that are not yet fully understood. We aimed to understand the evolution of PWO proteins and whether their interaction with PRC2 has been conserved through evolution. Our study reveals that PWO proteins are present in vascular plants, but absent in bryophytes and green algae. The ancestral clade of PWO proteins includes the *Selaginella moellendorffii* (Sm) PWO orthologs SmPWOa and SmPWOb. Transient expression assays showed that both AtPWO1 and SmPWOa form nuclear speckles where they tether At, Sm, but also *Physcomitrium patens* (Pp) PRC2 catalytic subunits, despite the absence of PWO proteins in Pp. The PWO-PRC2 interactions were confirmed by protein-protein analyses. A newly identified evolutionarily conserved short C-terminal alpha-helix (c-motif) in PWO proteins contributes to an interaction interface for PWO-PRC2 binding. SmPWOs partially rescue the *pwo1;pwo2* mutant phenotype in Arabidopsis, highlighting the functional conservation of PWOs in vascular plants.

## Introduction

The PWWP (Pro-Trp-Trp-Pro) domain, a member of the Royal family of domains [Tudor, Chromo (chromatin-binding), MBT (malignant brain tumor), and PWWP domains], functions as a recognition domain that binds both DNA and methylated lysine residues within histones (Qin and Min 2014; Rona et al. 2016). PWWP domain-containing proteins are usually part of complexes associated with chromatin structure, facilitating crosstalk between different epigenetic marks, and thereby regulating gene expression (Qin and Min 2014; Tan et al. 2018). Proteins carrying the PWWP domain are found in both plants and animals, where the PWWP domain occurs either as a single conserved domain or in a conserved combinatorial arrangement with other domains (Alvarez-Venegas and Avramova 2012). Among PWWP domain-containing proteins, the PWWP-DOMAIN INTERACTOR OF POLYCOMBS (PWO/PWWP) proteins are characterized by a distinct single N-terminal PWWP domain without any additional structured domains (Hohenstatt et al. 2018; Tan et al. 2018).

In the flowering dicot model *Arabidopsis thaliana* (Arabidopsis, At), three PWO orthologs— AtPWO1, AtPWO2, AtPWO3 - are present and play important roles in plant development. Homozygous *pwo* triple mutant plants (*pwo1;pwo2;pwo3*) are lethal at an early seedling stage but milder phenotype defects are observed in *pwo* single and double mutant plants (Hohenstatt et al. 2018). PWO1 interacts with the evolutionarily conserved histone methyltransferase subunits of the Polycomb Repressive Complex 2 (PRC2), namely CURLY LEAF (CLF) and SWINGER (SWN) (Hohenstatt et al. 2018). These subunits contain a Su(var)3-9, Enhancer-of-zeste (E(z)), and Trithorax (SET) domain that catalyzes the deposition of H3K27me3 at specific loci resulting in transcriptional gene silencing (Baile et al. 2022). A genetic interaction between *PWO1* and *CLF* exists —*pwo1;clf* enhances the *clf* phenotype, exacerbating upward leaf curling and affecting the transcription of PRC2 target genes (Hohenstatt et al. 2018). Recently, PWO1 was shown to bind the boundary regions of H3K27me3-enriched compartment domains (CDs), which are topologically associated chromatin domains positioned at the nuclear periphery. In this context, PWO1 maintains the repressive structure of H3K27me3-CDs (Yang et al. 2024). PWO1 also associates with the lamin-like protein CROWDED NUCLEI 1 (CRWN1), controlling nuclear morphology and regulating partially overlapping sets of target genes in Arabidopsis (Mikulski et al. 2019). PWOs may therefore act as a bridge connecting H3K27me3-CDs and CRWN1 at the nuclear periphery and contributing to a repressive chromatin state (Yang et al. 2024). In addition, PWOs have been identified as core components of the PEAT complex, which includes PWOs, AT-rich interaction domain-containing proteins (ARIDs), and TELOMERE REPEAT BINDING proteins (TRBs). Although it was initially proposed that PEAT complex maintains heterochromatin silencing through its interaction to histone deacetylases (Tan et al. 2018), more recently it has been shown to interact with histone acetyltransferases of the MYST family (HAMs) and UBIQUITIN PROTEASE 5 (UBP5) to induce gene expression by histone 4 (H4) acetylation and H2A deubiquitination (Zheng et al. 2023; Godwin et al. 2024). Despite PWO1 ability to contribute to different chromatin protein networks, the molecular mechanisms underlying the contrasting effects of the putative PWO- associated protein complexes on chromatin structure remain elusive.

Emerging evidence shows that PRC2 is evolutionarily conserved in unicellular and multicellular eukaryotes (Schubert 2019; Déléris et al. 2021; Sharaf et al. 2022; de Potter et al. 2023; Hisanaga et al. 2023). PRC2-mediated gene silencing is crucial for cell differentiation, developmental transitions, and the establishment and maintenance of cell and organ identity in both plants and animals (Hennig and Derkacheva 2009; de Lucas et al. 2016; Schuettengruber et al. 2017). Mutation of PRC2 subunits, such as CLF and SWN, leads to severe developmental defects in Arabidopsis, characterized by the formation of masses of aerial tissues that resemble cotyledonary/callus-like structures (Chanvivattana et al. 2004; Mozgová et al. 2017; Shu et al. 2020). Loss of PRC2 also leads to severe developmental defects in basal land plants. In the moss *Physcomitrium patens,* loss of *CLF* results in apogamy, where a sporophyte-like body emerges from a gametophytic vegetative cell (Okano et al. 2009). In addition to developmental genes, PRC2 also targets genes involved in environmental sensing and stress response, extending its functions beyond developmental control (Folsom et al. 2014; Vyse et al. 2022; Faivre et al. 2024; Zarif et al. 2024). Considering the significant innovations associated with the evolution of land plants (Donoghue et al. 2021), it is intriguing to ask how the functions of PRC2 have evolved to meet the demands for developmental and environmental acclimation in land plants. This raises compelling associated questions about the molecular adaptations and regulatory mechanisms that enabled PRC2 to integrate other accessory proteins, that potentially play a crucial role in transcriptional regulation in terrestrial ecosystems. PWOs, as components of both repressive (e.g., PRC2 complex) and activating (e.g., PEAT complex) regulatory complexes (Hohenstatt et al. 2018; Tan et al. 2018; Zheng et al. 2023; Godwin et al. 2024), may play key roles in transcriptional switches associated with environmental and developmental responses, adjusted during land plant evolution.

In this study, we address the evolution and function of PWO proteins, with a focus on the conservation of PWO-PRC2 interaction. We show that PWOs emerge in lycophytes but are absent in bryophytes and green algae. To understand molecular interactions and functions of PWOs in plant evolution, we study *Selaginella moellendorffii* (Sm) SmPWOa and SmPWOb as representatives of early-emerging PWOs. We demonstrate that, similar to AtPWO1, SmPWOa physically interacts with CLF, tethering CLF orthologs from different species to the sub-nuclear compartment where it localizes. In addition to the N-terminal PWWP domain, we identify a novel conserved C-terminal motif (C-motif) that is required for PWO interaction with PRC2 catalytic subunits throughout evolution. Finally, we demonstrate that SmPWOs partially complement the *pwo1;pwo2* double mutant phenotype in Arabidopsis. Altogether, our data indicate that PWOs have been closely associated with PRC2 since their emergence in lycophytes, suggesting a deep evolutionary origin of their function in coordination with the Polycomb Group (PcG) repressive pathway.

## Results

### PWO proteins are found in vascular plants and form four major clades

To identify PWO orthologs in the green lineage and address their evolutionary relationships, we used a previously established bioinformatic pipeline (Sharaf et al. 2022). We searched for orthologs of AtPWO1 in 183 species representing Chlorophyta and Streptophyta, including algae and land plants. PWO orthologs were identified in 66 species (36%) (Supplementary Table 1), all representing land plants. Interestingly, PWO orthologs were absent in non-vascular plant groups but were well conserved in vascular plants (Tracheophyta), first appearing in extant lycophytes (Lycopodiopsida) (Fig. 1A; Supplementary Table 1).

**Figure 1.**
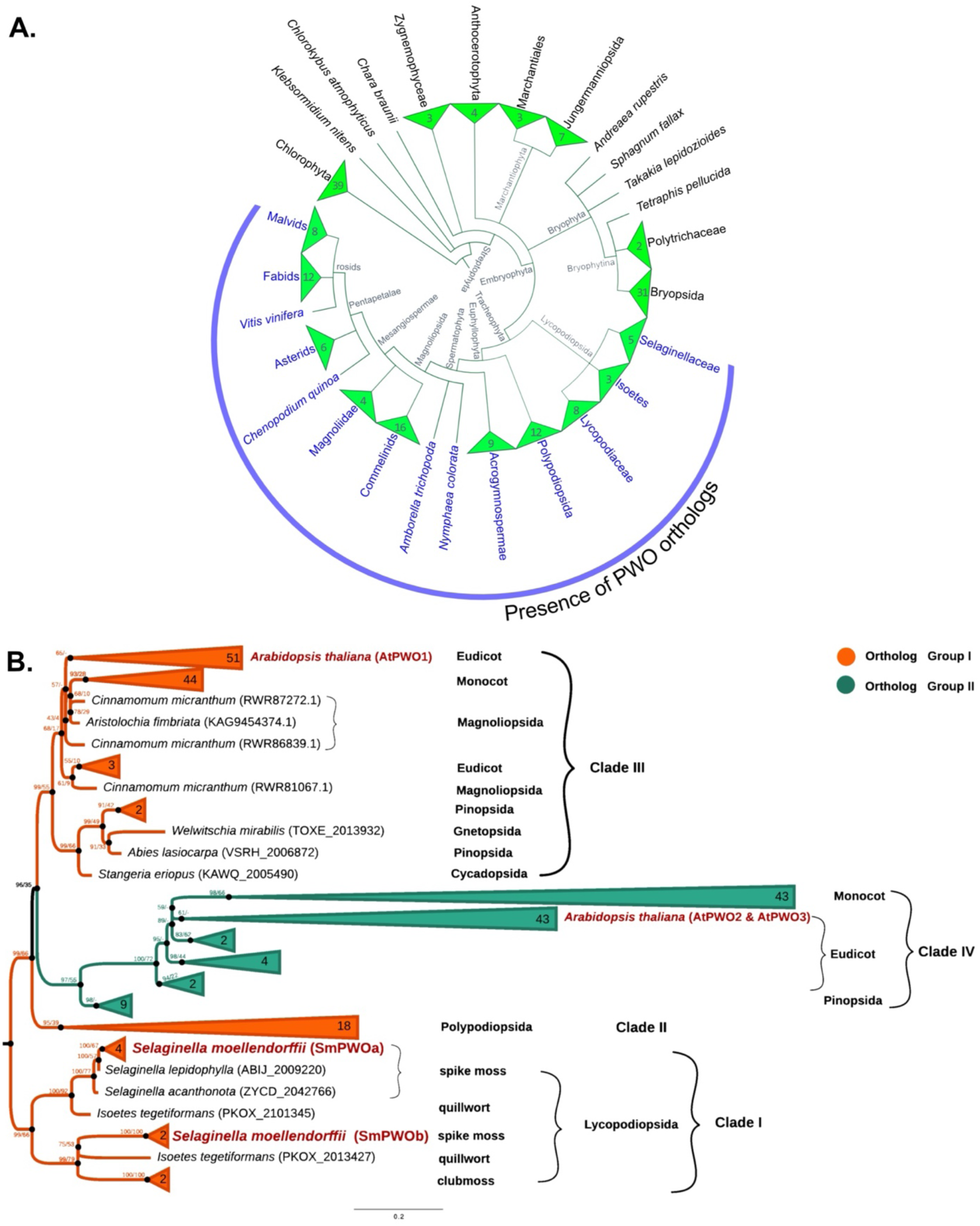
The evolution and distribution of PWO proteins in Viridiplantae. **A.** Cladogram showing the diversity of PWO protein orthologs across the green lineage. Blue line indicates presence of PWO orthologs**. B.** Maximum-likelihood (ML) phylogenetic tree of PWO protein orthologs showing separation into four clades (Clade I, Clade II, Clade III, Clade IV). The tree was constructed using the whole protein sequence, and the ML branch support values are given in % (IQ-TREE/RAxML-NG). The phylogenetic tree was rooted with sequences of Lycopodiopsida PWO orthologs.

The identified PWO sequences clustered into four major clades (Clades I-IV) (Fig. 1B; Supplementary Table 2). Moreover, orthology assignment using the EggNOG database (Huerta-Cepas et al. 2019a) identified two orthology groups: ortholog group I (ENOG5028MA3) and ortholog group II (ENOG5028N12) (Fig. 1B). The orthologs of the first group clustered into three clades (Fig. 1B). Clade I (“ancestral” PWO clade) comprised lycophyte orthologs, where orthologs of clubmosses, spike mosses and quillworts diverged into two subclades. One of the subclades included *S. moellendorffii* SmPWOa paralogs. The other subclade included the SmPWOb paralogs. PWO clade II included Polypodiopsida (ferns and horsetails) orthologs, grouped into a monophyletic clade at the root of the remaining clades (Fig. 1B). PWO clade III clustered at the crown of the tree and contained orthologs of seed plants—gymnosperms (Cycadopsida, Pinopsida, Gnetopsida) and angiosperms (Magnoliopsida, monocots, and eudicots). This clade included the queried AtPWO1. Finally, clade IV, which belongs to the second orthology group, clustered at the root of clade III and contained orthologs of seed plants (Pinopsida, monocots, and eudicots), including AtPWO2 and AtPWO3.

### PWOs structural prediction shows an evolutionarily conserved PWWP domain and a short C-motif

PWO proteins share (Fig. 2A) a conserved N-terminal PWWP domain including a SWWP motif, a variation of this domain (Supplementary Fig. 1) (Qin and Min 2014; Rona et al. 2016). In addition, we identified one or more predicted nuclear localization signals (NLS) in the central region (Supplementary Table 3) and a conserved sequence motif at the C-terminus, referred to as the C-motif from here on. The 3D structures of full-length PWO sequences from two species, representing three clades of ortholog groups I–II, were predicted using AlphaFold2 (AF2) (Jumper et al. 2021; Varadi et al. 2024). PWOs of the dicot *A. thaliana* represented Clade III (AtPWO1) and Clade IV (AtPWO2, AtPWO3) (Fig. 2B-D; Supplementary Table 4). PWOs of the spike moss *S. moellendorffii* (SmPWOa, SmPWOb) represented the ancestral Clade I (Fig. 2F-G). The most structured regions of the PWOs (predicted Local Distance Difference Test (pLDDT) > 70) were within the N-terminal PWWP domain (Supplementary Fig. 2A-B; Fig. 2B-D, 2F-G) and the short C-terminal C-motif, which folded into a short alpha-helical structure (Fig. 2E, 2H). Outside of these regions, AtPWOs and SmPWOs exhibited long stretches of low confidence structure (per-residue model confidence score (pLDDT) < 50), consistent with intrinsically disordered regions (IDRs) (Supplementary Fig. 2A; Fig. 2B-D, 2F-G) (Erdős and Dosztányi 2024).

**Figure 2.**
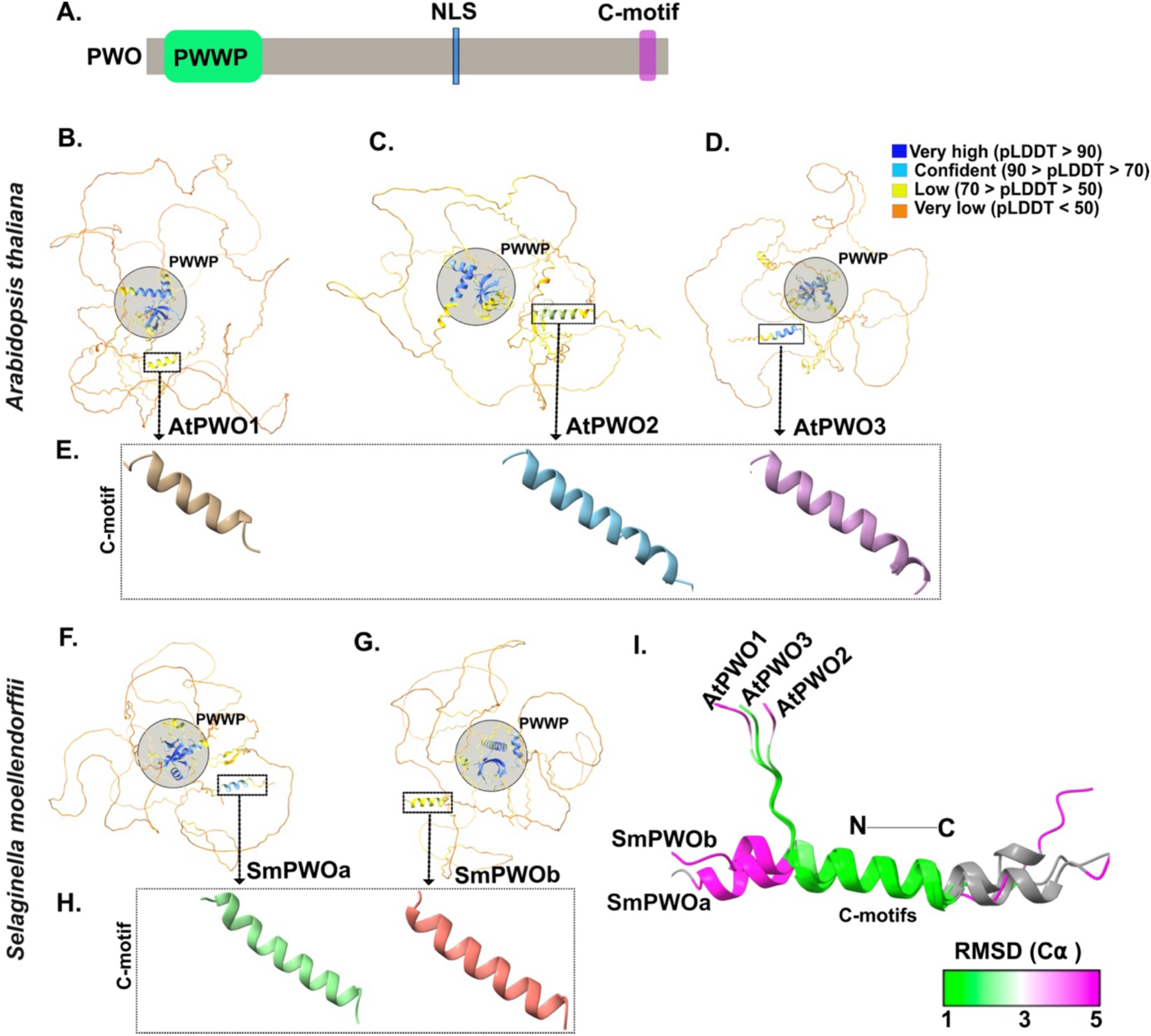
AlphaFold2-based protein structure and domain/motif prediction of PWO proteins. **A.** Schematic representation of full-length PWO proteins showing the conserved N-terminal PWWP domain and C-terminal C-motif. PWOs have one or several predicted nuclear localization signal(s) (NLS), here represented by a single one. Structure prediction of **B-D.** Arabidopsis PWO orthologs (AtPWO1, AtPWO2, AtPWO3) including their **E.** C-motif (boxed). **F-G.** *S. moellendorffii* PWO orthologs (SmPWOa, SmPWOb), showing the PWWP domain, and **H.** C-motif (boxed). The predicted full-length PWO structures are depicted with per-residue confidence scores (pLDDT) represented by different colors, ranging from 0 to 100, where regions with pLDDT below 50 may be considered unstructured. Different pseudocolours are used to depict the C-motif (E, H). **I.** Superposition of predicted C-motifs from *A. thaliana* and *S. moellendorffii* PWO orthologs on the AtPWO1 C-motif (used as reference). The Root Mean Square Deviation of Alpha Carbon atoms (RMSD (C α)) color range is displayed, with green representing values of 1 Å or less indicating high structural similarity, white for intermediate values around 3 Å, and magenta for an RMSD of 5 Å indicating the least structural similarity.

The predicted PWWP domains of PWO orthologs from *A. thaliana* (Supplementary Fig. 3A-E) and *S. moellendorffii* (Supplementary Fig. 3A, B, F-G) corresponded to the canonical structure of PWWP domains (Wu et al. 2011; Qin and Min 2014) with five antiparallel β-strands (β1– β5) arranged in a β-barrel, followed by three α-helices. The position of α0 was variable, while α1 and α2 consistently formed after the β-barrel structure (Supplementary Fig. 3C-G). Superimposition of the PWWP domains of *A. thaliana* and *S. moellendorffii* PWO orthologs showed structural conservation between AtPWO1 and AtPWO2-3, as well as SmPWOa-b (Supplementary Fig. 3H; Supplementary Table 5).

The predicted C-motif from *A. thaliana* and *S. moellendorffii* PWOs also exhibited significant structural similarity when superimposed onto the AtPWO1 C-motif (Fig. 2I; Supplementary Table 6). Moreover, prediction of 18 more PWO sequences representing different species from all four PWO clades confirmed consistent presence of the C-motif (Supplementary Fig. 4A). Superposing all the selected C-motifs onto the AtPWO1 C-motif highlighted structural conservation (Supplementary Fig. 4B; Supplementary Table 6). Multiple sequence alignment showed high level of amino acid (aa) conservation within the C-motif, enriched with hydrophobic aa residues (Φ) (CΦPΦK/RΦΦΦXRΦXEXΦ) (Supplementary Fig. 5).

### AtPWO1 and SmPWOs form nuclear speckles and colocalize in the same subnuclear space within plant nuclei

The predicted structures of PWO proteins contained long stretches of IDRs (Fig. 2B-D, F-G; Supplementary Fig. 2), which are often associated with a high propensity to form condensates (Solis-Miranda et al. 2023). Additionally, one or more NLSs (Supplementary Table 3) located within the middle domain of the PWO proteins suggested their localization within the nucleus. Fluorescently tagged AtPWO1 and SmPWOs transiently expressed in *N. benthamiana* showed that GFP-SmPWOa (Fig. 3A) and GFP-SmPWOb (Fig. 3B), as well as AtPWO1-mCherry (Fig. 3C; (Hohenstatt et al. 2018)) form speckles in the nucleoplasm. Notably, GFP-SmPWOb, unlike GFP-SmPWOa or AtPWO1-GFP, localized to the nucleolus (Fig. 3A-C). The localization of stably expressed SmPWOa-GFP and SmPWOb-GFP in *A. thaliana* (Fig. 3D, 3E) resembled that in *N. benthamiana* (Fig. 3A, 3B). The co-expression of GFP-SmPWOa with AtPWO1- mCherry (Fig. 3F) or mCherry-SmPWOb (Fig. 3G) resulted in the colocalization of the nucleoplasmic speckles. Similarly, the co-expression of GFP-SmPWOb and AtPWO1-mCherry showed colocalization of speckles within the nucleoplasm (Fig. 3H). Additionally, SmPWOb also localized in the nucleolus (Fig. 3G-H). Importantly, mCherry expressed alone did not form speckles and was excluded from the nucleolus, indicating that PWOs are needed to promote the speckle formation and/or nucleolar localization (Supplementary Fig. 6A-C). Overall, SmPWOs and AtPWO1 form nucleoplasmic speckles that share the same subnuclear space, while SmPWOb additionally localizes to the nucleolus.

**Figure 3.**
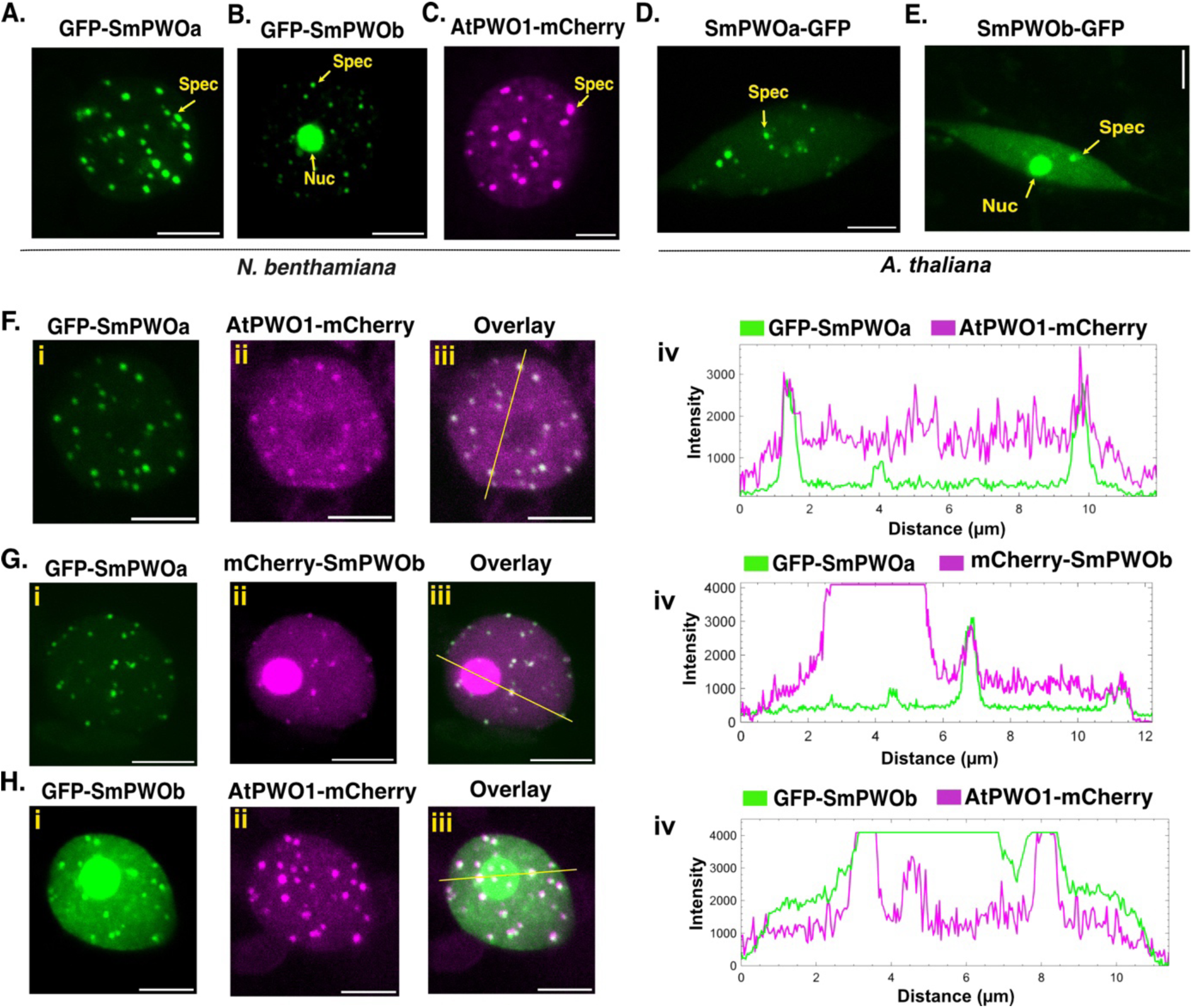
*In planta* subcellular localization of PWOs. **A-C**. Representative confocal microscopy images of *N. benthamiana* leaf nuclei, infiltrated with: **A.** *i35S_pro_::GFP-SmPWOa*, **B.** *i35S_pro_::GFP-SmPWOb*, and **C.** *i35S_pro_::AtPWO1- mCherry*. **D-E**. Transgenic *A. thaliana* lines expressing **D.** *2X35_pro_::SmPWOa-GFP* and **E.** *2X35_pro_::SmPWOb-GFP* in the *pwo1-1;pwo2-1* double mutant background in root tissues. The most prominent subnuclear structures are marked with arrows. ‘Spec’- speckles, and ‘Nuc’ - nucleolus. **F – H.** Representative confocal microscopy images of nuclei of *N. benthamiana* leaf co-infiltrated with: **Fi-iii.** *i35S_pro_::GFP-SmPWOa* and *i35S_pro_::AtPWO1-mCherry*, **Gi-iii.** *i35S_pro_::GFP-SmPWOa* and *i35S_pro_::mCherry-SmPWOb*, **Hi-iii.** *i35S_pro_::GFP-SmPWOb* and *i35S_pro_::AtPWO1-mCherry*. **F-Hiv**. Profiles of mCherry and GFP fluorescence intensities along the yellow line indicated in F-Giii. Scale bar = 5 µm.

### PWOs tether CLF from the nucleoplasm to sub-compartments in plant nuclei

Previously, AtPWO1 was shown to interact with CLF, the catalytic subunit of PRC2, and to tether AtCLF to nuclear speckles (Hohenstatt et al. 2018). This led us to investigate whether the tethering of CLF to PWOs-derived nuclear speckles is conserved across PWOs and CLF orthologs in other species. When expressed alone in *N. benthamiana*, mCherry-AtCLFΔSET, which has previously been shown to display identical localization and interacting behavior as the full-length protein (Hohenstatt et al. 2018; Mikulski et al. 2019; Godwin et al. 2024), resulted in diffuse nuclear localization but distinct nuclear speckles were never observed (Fig. 4A). In contrast, co-expressing mCherry-AtCLFΔSET, with AtPWO1-GFP changed the nuclear distribution of mCherry-AtCLFΔSET in 36% of the nuclei, in which mCherry-AtCLFΔSET co-localized with AtPWO1-GFP speckles (Fig. 4B; Supplementary Fig. 7A). The co-expression of GFP-SmPWOa and mCherry-AtCLFΔSET resulted in the formation of mCherry-AtCLFΔSET speckles that overlapped with SmPWOa speckles in 68% nuclei. (Fig. 4C; Supplementary Fig. 7B). Similarly, co-expression of GFP-SmPWOb and mCherry-AtCLFΔSET resulted in the formation of mCherry-AtCLFΔSET speckles, although only in 18% of the nuclei (Fig. 4D; Supplementary Fig. 7C-E) and, interestingly, promoted the localization of mCherry-AtCLFΔSET to the nucleolus in 90% of the nuclei (Fig. 4D; Supplementary Fig. 7C-D). Thus, the presence of PWOs potentiates the relocation of AtCLF within the nucleus.

**Figure 4.**
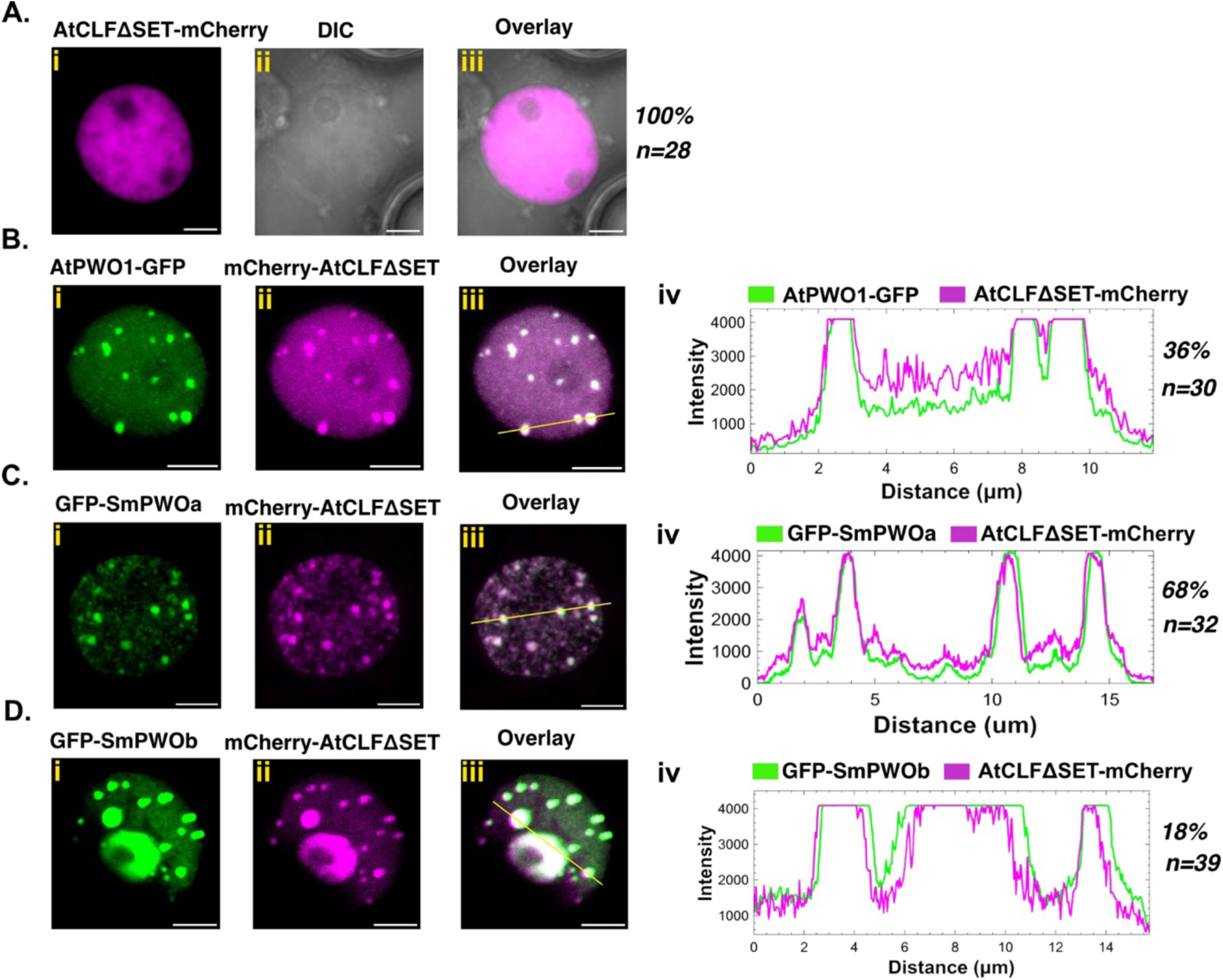
Colocalization of *A. thaliana* and *S. moellendorffii* PWOs with AtCLF. Representative confocal microscopy images showing nuclei of *N. benthamiana* leaf cells infiltrated with **A.i-iii.** *i35S_pro_::mCherry-AtCLFΔSET*, or co-infiltrated with **Bi-iii.** *i35S_pro_::AtPWO1-GFP* and *i35S_pro_::mCherry-AtCLFΔSET*, **Ci-iii.** *i35S_pro_::GFP-SmPWOa* and *i35S_pro_::mCherry-AtCLFΔSET* and **Di-iii.** *i35S_pro_::GFP-SmPWOb* and *i35S_pro_::mCherry-AtCLFΔSET*. **B-Div** Profiles of mCherry and GFP fluorescence intensities along the yellow line in B-Diii. Percentage of nuclei with the observed pattern versus total number of analyzed nuclei (n) is indicated on the right. ‘ΔSET’ denotes the deletion of SET domain. Scale bar = 5 µm.

Next, we asked whether AtPWO1 and SmPWOs could recruit SmCLF to PWO speckles. The localization of SmCLF-mCherry alone showed uniform distribution within the nucleoplasm in approximately 32% of nuclei (Fig. 5A), while the remaining nuclei showed strong accumulation of the SmCLF-mCherry signal (Supplementary Fig. 8A). Notably, this accumulation did not resemble the typical speckle pattern (Fig. 5A; Supplementary Fig. 8A). When AtPWO1-GFP and SmCLF-mCherry were co-expressed, distinct patches of SmCLF-mCherry overlapping with AtPWO1-GFP were formed in 65% of nuclei (Fig. 5B). The remaining 35% nuclei showed evenly distributed SmCLF-mCherry, which was not concentrated into AtPWO1-GFP speckles (Supplementary Fig. 8B). Similarly, co-expression of GFP-SmPWOa and SmCLF-mCherry resulted in 62% of nuclei displaying SmCLF-mCherry speckles (Fig. 5C), while 23% of nuclei showed small patches of SmCLF-mCherry with no substantial co-localization with GFP- SmPWOa (Supplementary Fig. 8C). The remaining 15% of nuclei showed a marginal overlap of GFP-SmPWOa speckles with SmCLF-mCherry (Supplementary Fig. 8D). The co-expression of GFP-SmPWOb and SmCLF-mCherry led to a subtle change in the localization of SmCLF- mCherry from the nucleoplasm to the nucleolus, with no distinct formation of SmCLF-mCherry speckles (Fig. 5D).

**Figure 5.**
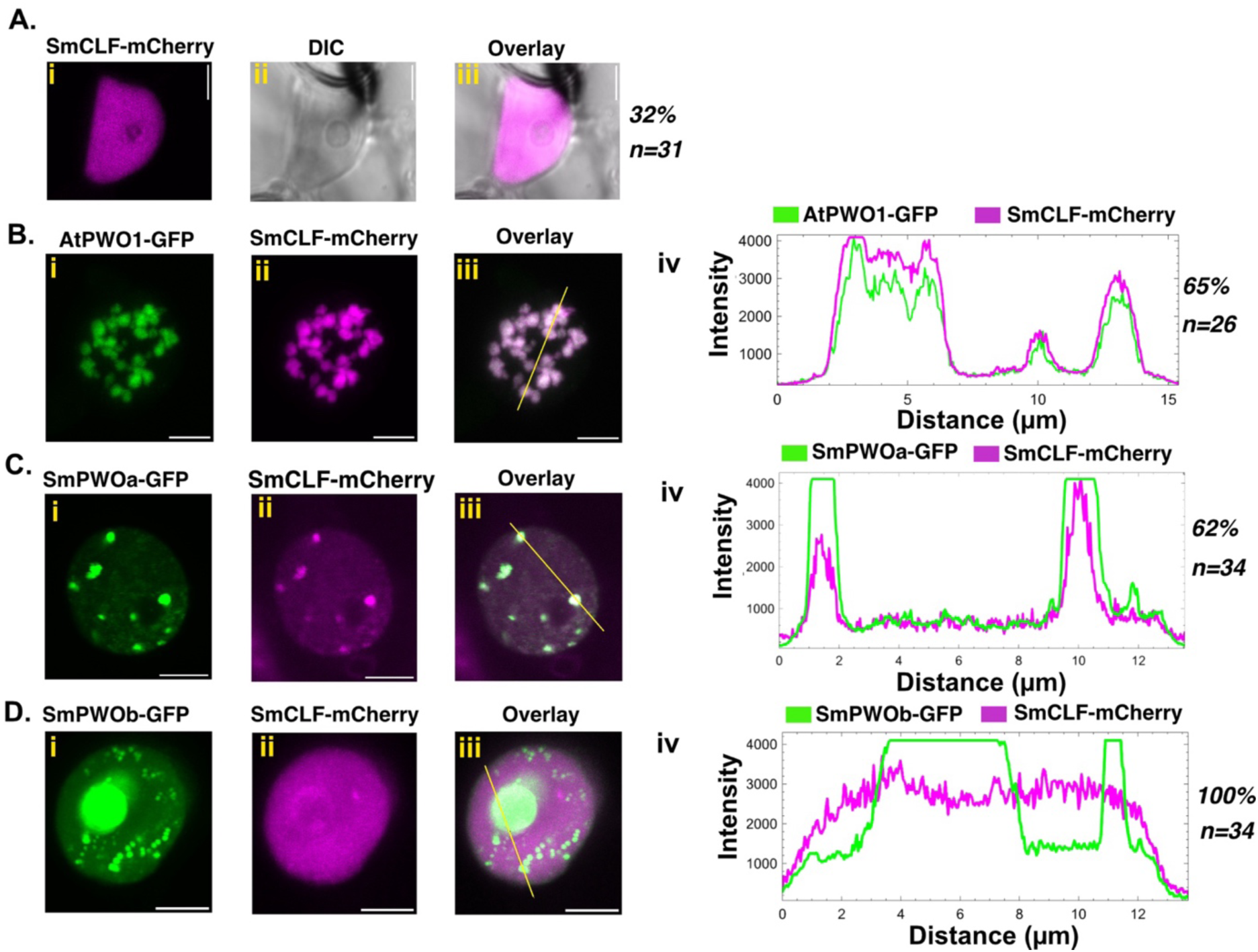
Colocalization of *S. moellendorffii* (Sm)CLF with AtPWO1, SmPWOa and SmPWOb in *N. benthamiana*. Representative confocal microscopy images showing nuclei of *N. benthamiana* leaf cells infiltrated with **A.i-iii.** *i35S_pro_::SmCLF-mCherry*, and co-infiltrated with **B.i-iii.** *i35S_pro_::AtPWO1-GFP* and *i35S_pro_::SmCLF-mCherry*, **Ci-iii.** *i35S_pro_::GFP-SmPWOa* and *i35S_pro_::SmCLF-mCherry* and **D.i-iii.** *i35S_pro_::GFP-SmPWOb* and *i35S_pro_::SmCLF-mCherry*. **B-D.iv** Profiles of mCherry and GFP fluorescence intensities along the yellow line in B-Diii. Percentage of nuclei with the observed pattern versus total number of analyzed nuclei (n) is indicated on the right. Scale bar = 5 µm.

Finally, we asked whether PWOs can tether PpCLF from *P. patens*, which lacks PWO proteins (Fig. 1A). When expressed alone, PpCLF-mCherry diffusely localized in the nucleoplasm in non-uniform patterns but did not form distinct speckles (Fig. 6A). Co-expression of AtPWO1- GFP (Fig. 6B) or GFP-SmPWOa (Fig. 6C) and PpCLF-mCherry resulted in the formation of PpCLF-mCherry speckles in the nucleoplasm of 44% and 78% of nuclei, respectively. These speckles overlapped with AtPWO1-GFP or GFP-SmPWOa speckles (Fig. 6B-C; Supplementary Fig. 9A-C). In contrast, co-expression of GFP-SmPWOb and PpCLF-mCherry resulted neither in the formation of PpCLF-mCherry speckles in the nucleoplasm nor in PpCLF- mCherry nucleolar signal (Fig. 6D).

**Figure 6.**
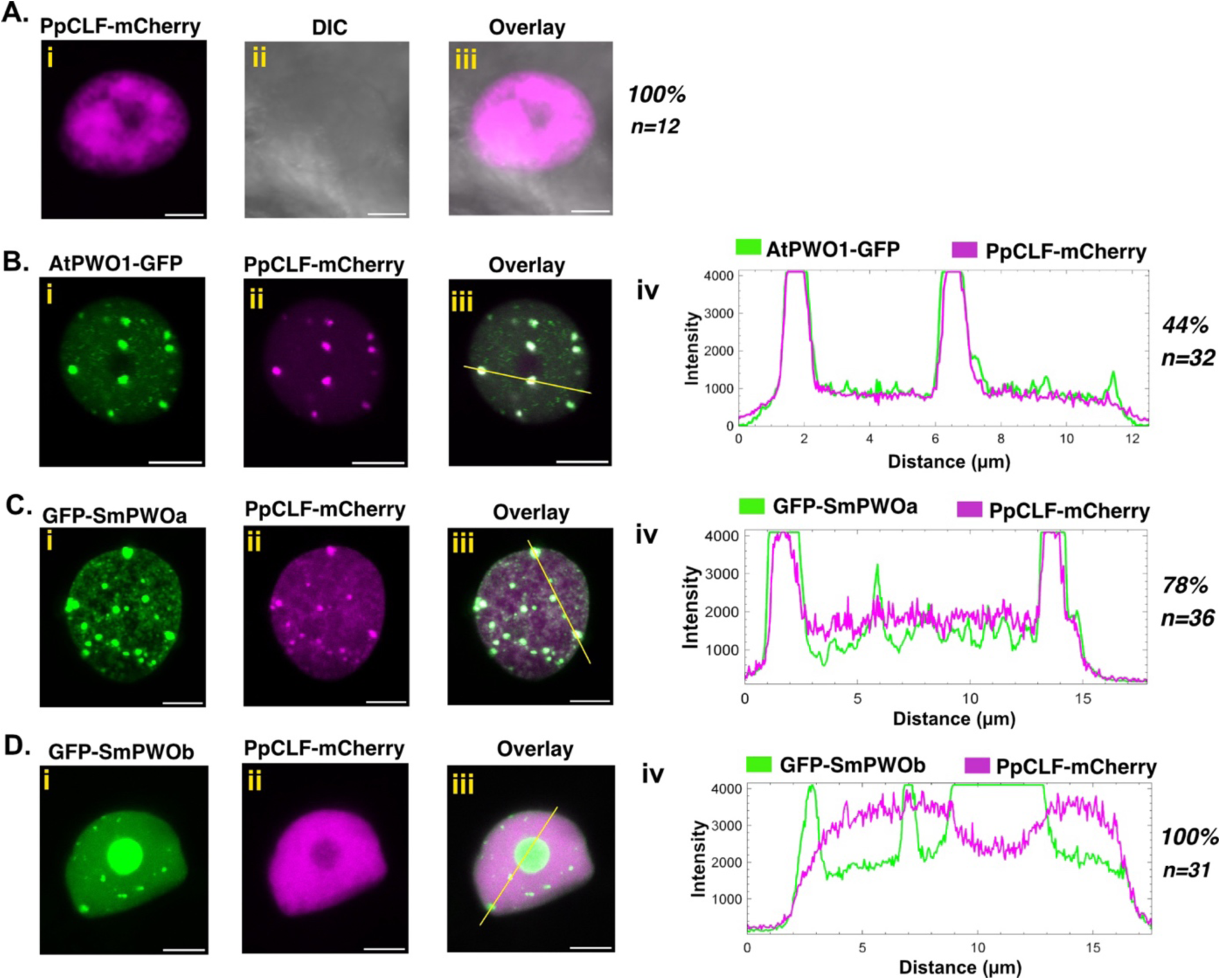
Colocalization of *P. patens* (Pp)CLF with AtPWO1, SmPWOa and SmPWOb in *N. benthamiana*. Representative confocal microscopy images showing nuclei of *N. benthamiana* leaf cells infiltrated with **Ai-iii.** *i35S_pro_::PpCLF-mCherry*, or co-infiltrated with **Bi-iii.** *i35S_pro_::AtPWO1- GFP* and *i35S_pro_::PpCLF-mCherry*, **C.i-iii.** *i35Spro::GFP-SmPWOa* and *i35S_pro_::PpCLF- mCherry* and **Di-iii.** *i35S_pro_::GFP-SmPWOb* and *i35S_pro_::PpCLF-mCherry*. **B-Div:** Profiles of mCherry and GFP fluorescence intensities along the yellow line in B-Diii. Percentage of nuclei with the observed pattern versus total number of analyzed nuclei (n) is indicated on the right. Scale bar = 5 µm.

Overall, AtPWO1 and SmPWOa potentiate the relocation of AtCLF, SmCLF, and PpCLF to PWO-derived speckles. Most noticeably, SmPWOb changes the localization of AtCLF and SmCLF from the nucleoplasm to the nucleolus when co-expressed. Altogether, this suggests a conservation of function for AtPWO1 and SmPWOa in mediating CLF re-localization to specific PWO-enriched subnuclear speckles.

### SmPWOa physically interacts with PRC2 catalytic subunits from *A. thaliana*, *S. moellendorffii* and *P. patens*

AtPWO1 and SmPWOa tether the PRC2 catalytic subunit CLF to their subnuclear compartments in *N. benthamiana* (Fig. 4-6; Supplementary Fig. 7-9). Therefore, we asked whether this tethering involves direct physical interaction between PWOs and the CLF orthologs. Yeast two-hybrid (Y2H) assays identified a physical interaction between AtPWO1 and AtSWN (Fig. 7A, D) (Hohenstatt et al. 2018). The AtPWO1-AtCLF interaction, though weaker, was also confirmed (Fig. 7A, D). A weaker interaction is observed between AtPWO1 and PpCLF compared to AtPWO1-AtSWN, while no significant interaction occurs between AtPWO1 and SmCLF (Fig. 7A, D). Moreover, SmPWOa interacts with the PRC2 catalytic subunit of *S. moellendorffii* (SmCLF), *P. patens* (PpCLF), and *A. thaliana* (AtCLF/SWN) (Fig. 7B, D). A similar interaction for SmPWOb is not observed under high-stringency (-LWHA) Y2H conditions (Fig. 7C, D), but under less stringent conditions (-LWH), a weaker interaction with PpCLF is detected (Supplementary Fig. 10).

**Figure 7.**
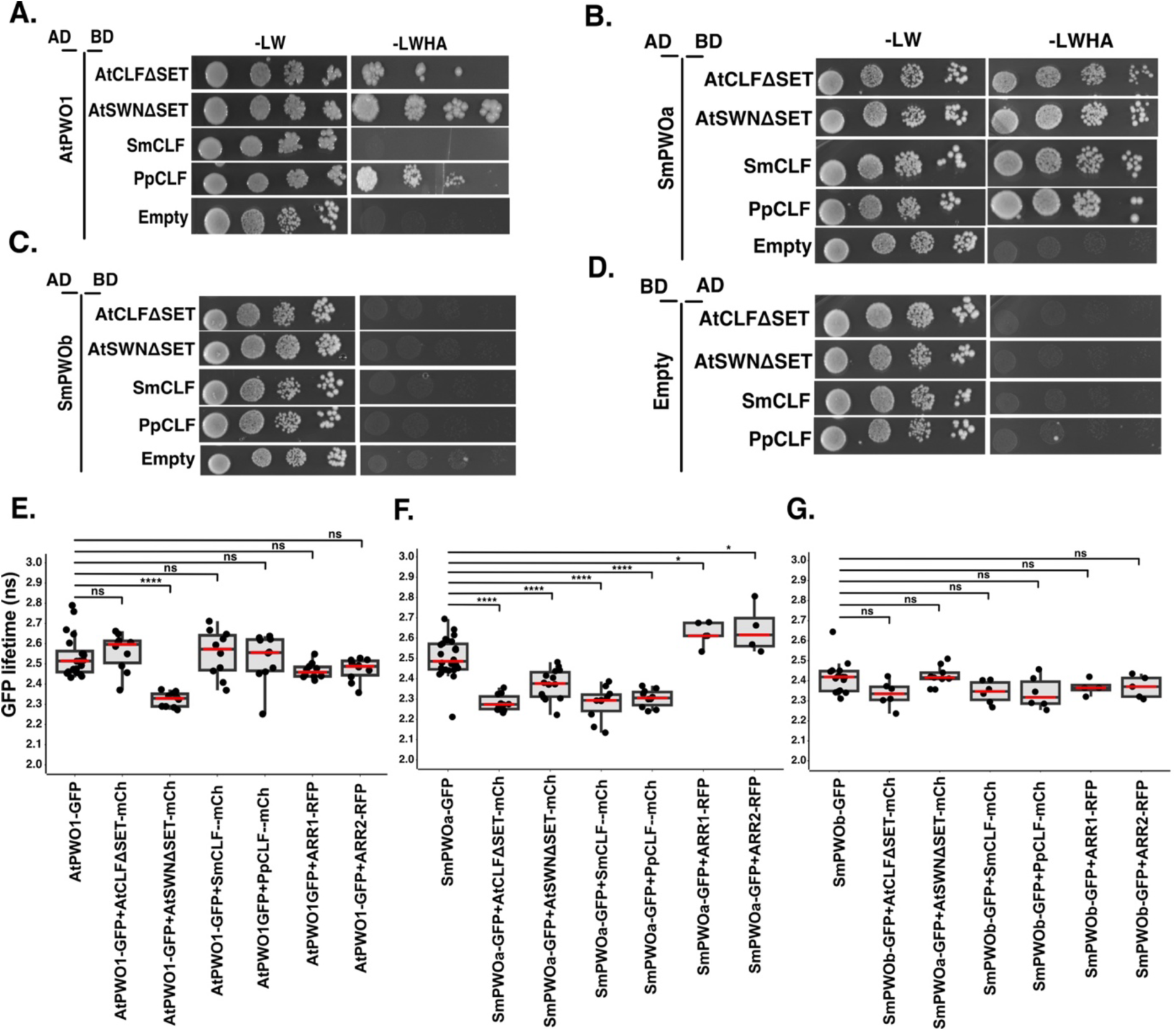
Protein-protein interactions of PWOs and PRC2 subunits. **A-C**. Yeast two-hybrid (Y2H) analyses of **A.** AtPWO1, **B.** SmPWOa, and **C.** SmPWOb with AtCLFΔSET, AtSWNΔSET, PpCLF, and SmCLF. In the control panel **D.**, interactions for all combinations were assessed using an empty ‘BD’ plasmid. The interactions were evaluated by growing transformed yeast cells on non-selective medium (−LW; lacking leucine and tryptophan) to facilitate plasmid co-transformation and on selective medium (−LWAH; lacking leucine, tryptophan, adenine, and histidine) to activate the reporter gene. In this setup, ‘BD’ refers to the GAL4-DNA binding domain, while ‘AD’ indicates the GAL4-DNA activation domain fusion. Constructs containing CLF and SWN with SET domain deletions are denoted as ΔSET. **E-G.** FLIM-FRET analyses of GFP lifetime in *N. tabacum* nuclei co-expressing *i35S_pro_::mCherry-AtCLFΔSET*, *i35S_pro_::SmCLF-mCherry*, and *i35S_pro_::PpCLF-mCherry* with **E.** *i35S_pro_::AtPWO1-GFP,* **F.** *i35S_pro_::GFP-SmPWOa* in speckles and **G.** *i35S_pro_::GFP- SmPWOb* in nucleolus. Speckles were selected for measurements. For the negative control, each setup included *35S_pro_::RFP-ARR1, 35S_pro_::RFP-ARR2* transcription factors co-expressed with PWOs (SmPWOa, SmPWOb, AtPWO1). Error bars represent SE. Statistical significance was determined using one-way ANOVA, with multiple comparisons using Tukey’s test. **** P ≤ 0.0001, *P ≤ 0.05, “ns” indicates not significant with P > 0.05.

Furthermore, Förster resonance energy transfer (FRET) experiments measured by Fluorescence Lifetime Imaging Microscopy (FLIM) were performed to confirm these interactions (Fig. 7E-G). The FLIM-FRET analyses were carried out either within nuclear speckles (AtPWO1, SmPWOa) or in the nucleolus (SmPWOb) of *N. tabacum* co-expressing PWOs with the CLF orthologs from different species. Co-expressing AtPWO1-GFP with CLF orthologs (AtCLF-, AtSWN-, SmCLF-, and PpCLF-mCherry) resulted in a decrease in the AtPWO1-GFP lifetime within nuclear speckles only when co-expressed with AtSWN-mCherry (Fig. 7E; Supplementary Fig. 11A). This supports the notion that AtPWO1 efficiently interacts with AtSWN but not with CLF from lower land plant models or not even with AtCLF. In contrast, co-expressing GFP-SmPWOa with AtCLF-, AtSWN-, SmCLF-, and PpCLF-mCherry resulted in a significant decrease in the fluorescence lifetime of GFP-SmPWOa compared to sole expression of GFP-SmPWOa, indicating direct physical interaction with all tested CLF orthologs (Fig. 7F; Supplementary Fig. 11B). However, co-expression of GFP-SmPWOb with AtCLF-, AtSWN-, SmCLF-, and PpCLF-mCherry resulted in no significant changes in the GFP-SmPWOb lifetime, indicating lack of interaction under our experimental conditions (Fig. 7G Supplementary Fig. 11C). As a negative control, we selected nuclear localized transcription factors ARABIDOPSIS RESPONSE REGULATOR1 (ARR1) and ARR2, which do not interact with PWOs, confirming the specificity of the interactions (Fig. 7E-G; Supplementary Fig. 11A-C). Overall, the Y2H and FLIM-FRET supported the notion that SmPWOa is able to promiscuously interact with CLF orthologs from different species across plant evolution.

### The conserved PWO C-motif provides an interface for interactions with CLF orthologs

Based on the colocalization (Fig. 3-6) and interaction (Fig. 7) of PWO and CLF orthologs, we proceeded to investigate the interaction interface between them. Using AtPWO1, AtSWN, and AtCLF, we modeled their interactions with AlphaFold2-Multimer (AF2-M) (Evans et al. 2021; Homma et al. 2023, 2024; Ibrahim et al. 2023). The interaction prediction for full-length AtPWO1-AtCLF, AtPWO1-AtSWN or SmPWOa-SmCLF did not detect a high-confidence interaction interface, possibly due to the presence of IDRs in AtPWO1 (Supplemental Fig. 12A- C; Supplementary Fig. 2, Supplementary Table 7). However, when we systematically analysed the interaction in a series of truncated PWO and CLF/SWN fragments using AF2-M, we found that the central region of CLF/SWN, interacted with a truncated fragment of AtPWO1/SmPWOa lacking the PWWP domain (Supplementary Fig. 13A, F, K). Specifically, the truncated fragments of AtCLF (166–364 aa) or AtSWN (166–364 aa) and AtPWO1 (377– 769 aa) (Supplementary Fig. 13B-E, G-J) displayed local interactions. We also predicted a similar interaction surface for truncated SmCLF (166–364) and SmPWOa (377–619) (Supplementary Fig. 13L-O). The C-motif (Fig. 2) of AtPWO1 and SmPWOa was identified as the primary interface within these fragments for interaction with AtCLF (Supplementary Fig. 13D-E), AtSWN (Supplementary Fig. 13I-J) and SmCLF (Supplementary Fig. 13N-O).

Given the conservation of the C-motif across PWO phylogeny (Supplementary Fig. 4), we modeled the interaction interfaces focusing on the PWO C-motif and its interactions with CLF and SWN in representative species from different phylogenetic groups, including *A. thaliana* (dicots) (Supplementary Fig. 14A), *Zea mays* (monocots) (Supplementary Fig. 14B), *Ceratopteris richardii* (Polypodiopsida) (Supplementary Fig. 14C), and *S. moellendorffii* (Lycopodopsida) (Supplementary Fig. 14D). The PWO C-motif was predicted to be involved in the interaction with CLF orthologs across diverse land plant species (Supplemental Fig. 14; Supplementary Table 7).

Truncated versions of AtPWO1-ΔC-motif (deletion of residues 739–750 aa) (Fig. 8A) and SmPWOa-ΔC-motif (deletion of residues 594–612 aa) (Fig. 8B) were used to assess the contribution of the C-motif to interactions with the CLF orthologs. The Y2H results showed a reduced interaction between AtPWO1-ΔC-motif and AtSWNΔSET compared to the control (AtPWO1 with AtSWNΔSET) (Fig. 8C, E). SmPWOa-ΔC-motif with AtCLFΔSET showed a complete loss of interaction (Fig. 8D, E) and its interaction with AtSWNΔSET was reduced compared to respective control setup using full-length SmPWOa (Fig. 8D, E). Deletion of the C-motif in SmPWOa did not considerably change the interaction between SmPWOa and SmCLF or PpCLF (Fig. 8D, E).

**Figure 8.**
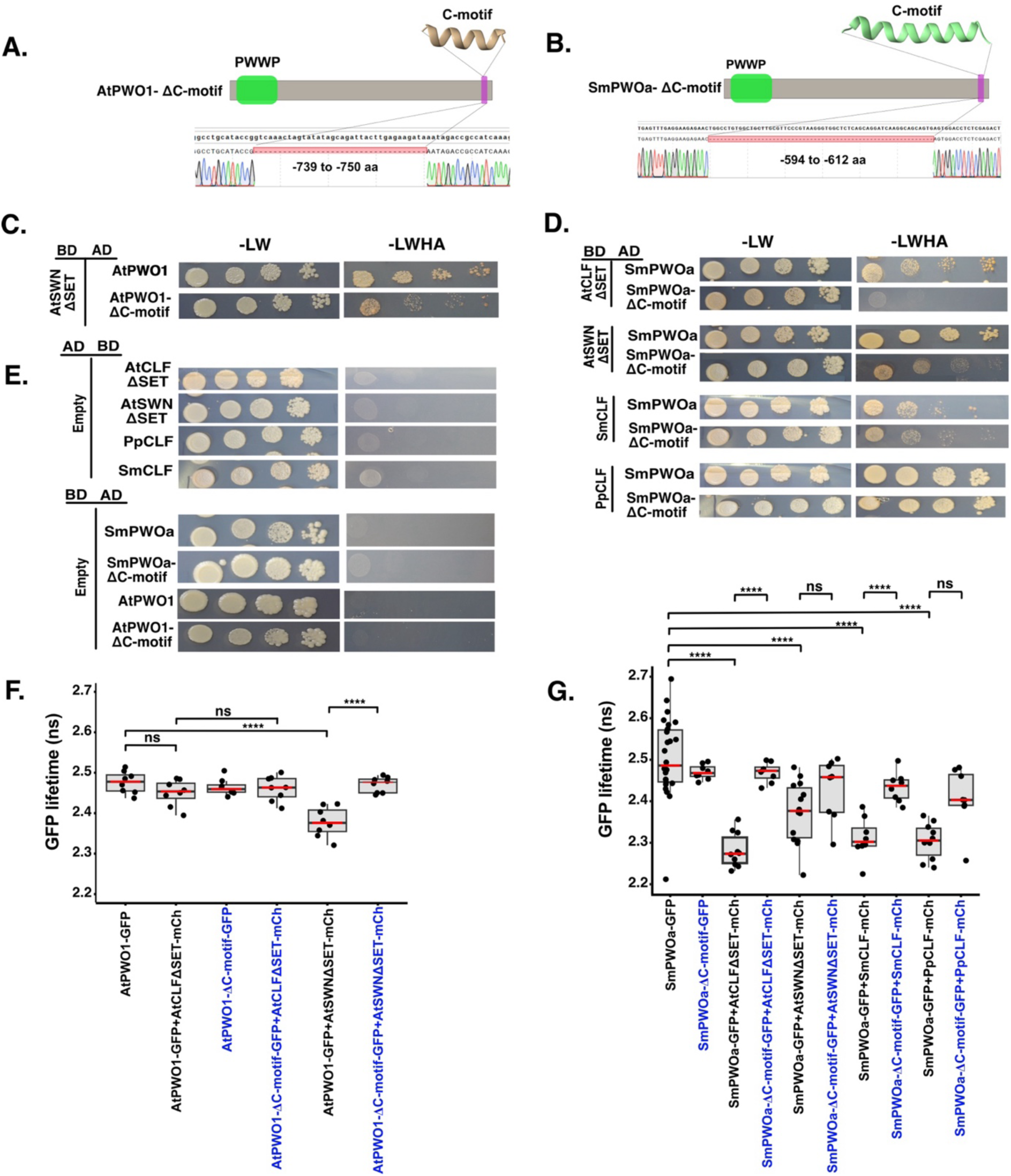
PWOs C-motif engagement in interaction with PRC2 catalytic subunits. Schematic representation of the C-motif deletion constructs of **A.** AtPWO1 and **B.** AtPWO1- ΔC-motif, used for Y2H and FLIM-FRET analysis. Y2H analysis showing interactions of **C.** AtPWO1 and AtPWO1-ΔC-motif (Δ739–750 aa) with AtSWNΔSET and **D.** SmPWOa and SmPWOa-ΔC-motif (Δ594–612 aa) with AtSWNΔSET, AtCLFΔSET, SmCLF, and PpCLF, using -LW (non-selective medium) and -LWAH (selective medium). **E.** In the control panel for all combinations, interactions were assessed with an empty ‘BD’ or ‘AD’ plasmid. The ‘BD’ represents the GAL4 DNA-binding domain, ‘AD’ represents the GAL4 DNA activation domain fusion and Δ represent for deletion of specific aa in plasmid. **F–G.** FLIM-FRET analysis of GFP lifetime in *N. tabacum* nuclei co-expressing **F.** *i35S_pro_::AtPWO1-GFP* and *i35S_pro_::AtPWO1-ΔC-motif-GFP* and **G.** *i35S_pro_::GFP-SmPWOa* and *i35S_pro_::SmPWOa-ΔC- motif-GFP with i35S_pro_::mCherry-AtCLFΔSET*, *i35S_pro_::mCherry-AtSWNΔSET*, *i35S_pro_::SmCLF-mCherry*, and *i35S_pro_::PpCLF-mCherry* within speckles. Error bars represent SE. Statistical significance was determined using one-way ANOVA with multiple comparisons using Tukey’s test. (****P ≤ 0.0001, *P ≤ 0.05); “ns” indicates not significant (P > 0.05). Blue highlighted combination represents C-motif deleted constructs.

Co-expression of AtPWO1-ΔC-motif-GFP with AtSWN-mCherry showed an increase in AtPWO1-ΔC-motif-GFP fluorescence lifetime compared to AtPWO1-GFP (WT) using FLIM-FRET (Fig. 8F), supporting a lower binding affinity of the AtPWO1-ΔC-motif for AtSWN. Similarly, co-expression of SmPWOa-ΔC-motif-GFP with CLF (AtCLF, AtSWN, SmCLF, and PpCLF)-mCherry resulted in a higher GFP fluorescence lifetime compared to full-length SmPWOa-GFP co-expressed with CLF or SWN (Fig. 8G), supporting a reduced affinity of SmPWOa-ΔC-motif for CLF and SWN across species (Fig. 8G). Overall, these results indicate that the C-motif participates in mediating PWO interactions with PRC2 components throughout plant evolution.

### *SmPWOa* partly complements the *pwo1-1;pwo2-2* double mutant phenotype in Arabidopsis

We expressed *SmPWOa-GFP* or *SmPWOb-GFP* in the Arabidopsis *pwo1-1;pwo2-2* double mutant background to determine their potential for restoring a WT-like phenotype. Nuclear localization of full-length SmPWOs-GFP was confirmed in the roots of the complemented plants (Fig. 3D-E). Interestingly, *SmPWOa-GFP* could fully or partially complement nearly all the developmental defects of the *pwo1-1;pwo2-2* mutant (i.e. root length, rosette area, plant height, nuclear size, and flowering time), while *SmPWOb-GFP* only partially complemented some of the traits (Figure 9 A-J, Supplementary Figure S15). For example, both expression lines showed an increase in root length compared to *pwo1-1;pwo2-2* double mutant but did not fully complement this trait (Figure 9A, 9E). *pwo1-1;pwo2-2* and *SmPWOb-GFP* displayed reduced rosette size (total leaf area) compared to WT, while rosette size was restored in *SmPWOa-GFP* (Figure 9B, 9F). A delay in flowering is observed in *pwo1-1;pwo2-2* double mutants, which was partially restored in *SmPWOa-GFP* line (Figure 9C, 9G). While *pwo1-1;pwo2-2* double mutants showed reduced height, *SmPWOa-GFP* plants were significantly taller than the mutant (Figure 9D, 9H). The loss of apical dominance in *pwo1-1;pwo2-2* double mutant was partially complemented by both constructs (Supplementary Fig. 15A). Similarly, *SmPWOa-GFP* partially restored reproductive defects in *pwo1-1;pwo2-2*, including the reduced silique number per plant and seed number per silique, whereas *SmPWOb-GFP* only partially complemented the silique number phenotype (Supplementary Figure S15B-C). Previously, *pwo1-1* mutants were shown to exhibit a reduced nuclear area and an increased circularity index (Mikulski et al. 2019). A similar phenotype was confirmed for *pwo1-1; pwo2-2* (Figure 9I–K). Nuclear area of *pwo1-1; pwo2-2* is complemented by both constructs, while circularity index is fully restored to WT-level only by *SmPWOa-GFP* (Figure 9I–K).

**Figure 9.**
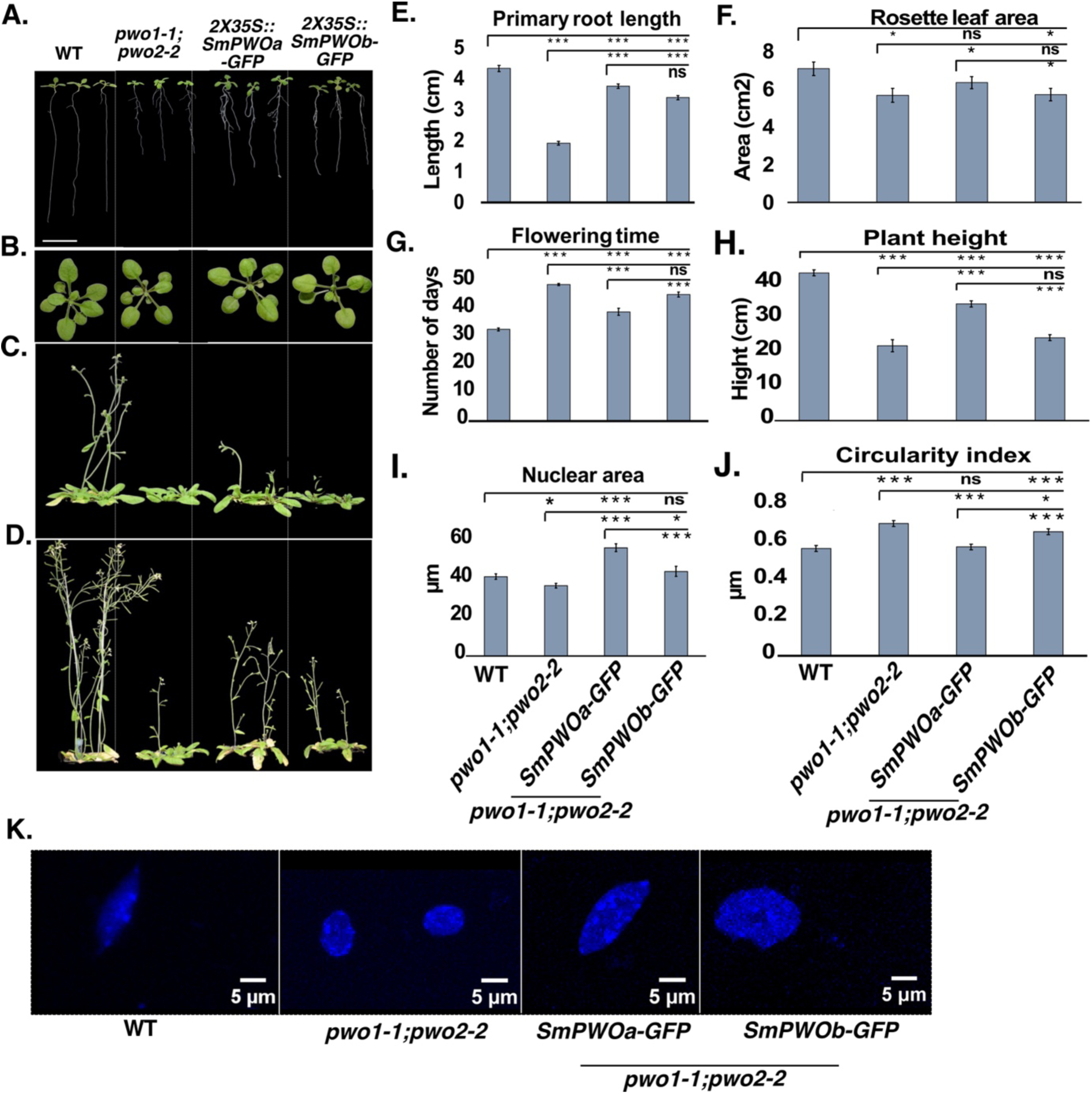
Phenotypic analyses of Arabidopsis *pwo1-1;pwo2-2* mutant lines overexpressing *SmPWOa* or *SmPWOb*. Representative images of **A.** Root length phenotype in 10-day-old seedlings; **B.** Rosette leaf area in 21-day-old plants; **C.** Flowering time phenotype in 39-day-old plants; and **D.** Plant height in 47-day-old plants for the genotypes Col-0, *pwo1-1;pwo2-2*, *2×35S_pro_::SmPWOa- GFP*/*pwo1-1;pwo2-2*, and *2×35S_pro_::SmPWOb-GFP*/*pwo1-1;pwo2-2* lines. **E.** Root length (n = 14); **F.** Rosette leaf area (n = 14); **G.** Flowering time (n = 10); **H.** Plant height (n = 10); **I.** Nuclear area (n = 100); and **J.** Nuclear circularity (n = 100) for the same genotypes. The nuclear area **I.** and circularity **J.** measurements showing the largest area of a particular nucleus (n = 100). **K.** Representative DAPI-stained nuclei from analyzed genotypes are shown. Error bars correspond to ±SD. Asterisks represent p-values: ***p ≤ 0.001, **p ≤ 0.01, *p ≤ 0.05. The two-tailed Student’s t-test was applied to calculate the significance level. Scale bars = 5 µm.

Published transcriptomic data have demonstrated the role of PWOs in the regulation of gene expression (Mikulski et al. 2019; Zheng et al. 2023; Yang et al. 2024). We selected several genes that are known to be misregulated in the *pwo1-1;pwo2-2* mutant background (Zheng et al. 2023) and tested their expression in the *SmPWOs-GFP* lines. *ACTIN1* (*ACT1*) and *SHOOT MERISTEMLESS* (*STM*) expression levels were fully restored by *SmPWOa-GFP*, while *SAMBA* was upregulated. On the other hand, *GAMETOPHYTE DEFECTIVE 1* (*GAF1*) and *SAMBA* expression was only complemented by *SmPWOb-GFP* (Supplementary Fig. 16).

These results together show that SmPWOa or SmPWOb can partially complement the lack of AtPWOs activity in Arabidopsis. SmPWOa consistently shows a higher potential for phenotypic complementation, suggesting a higher level of functional conservation.

## Discussion

### PWOs emerged from lycophytes and are well conserved in vascular plants

The transition from aquatic to terrestrial habitats is connected to the emergence of key new traits that enabled the plants to endure sudden environmental changes (Spencer et al. 2021; Woudenberg et al. 2022). These traits were linked with genetic variation that facilitated the emergence of new genes. Such genetic innovations primarily involved transcriptional regulators, which are typically associated with biotic and abiotic stress responses (Donoghue et al. 2021). PWOs emerged in Lycophytes, the early-diverged group of vascular plants (Fig.1A). Therefore, it is tempting to speculate that the appearance of PWOs at this critical point in plant evolution may have been necessary for land plants to acquire new developmental or adaptive traits required for survival in terrestrial environments. The presence of PWOs in distinct clades (Fig. 1B) highlights both evolutionary conservation and divergence, which may lead to sub- and neofunctionalization, facilitating the evolution of novel traits (Ohno 1970; Force et al. 1999; Zalewski et al. 2013). Two major separate clades of PWO proteins are found in seed plants, represented by *A. thaliana* AtPWO1 and AtPWO2/AtPWO3. Similarly, two independently emerged PWO clades are found in Lycopodiopsida, represented by *S. moellendorfii* SmPWOa and SmPWOb (Fig. 1B). Interestingly, SmPWOa and SmPWOb display distinct nuclear localization and protein-protein binding characteristics (Fig. 3-7), supporting at least partial divergence. The conservation of PWO proteins in vascular plants highlights their functional importance and suggests their essentiality. Indeed, full absence of PWOs in Arabidopsis is seedling lethal, suggesting a prime role for PWOs in plant development and a partial redundancy of AtPWO1-3 (Hohenstatt et al. 2018). Whether redundancy between PWOs may exist in lycophytes needs to be determined in the future. Nevertheless, considering the localization of SmPWOa (Fig. 3A, D) and SmPWOb (Fig. 3B, E) within the nucleus, it is possible that SmPWOs may function together or exhibit overlapping functions. Therefore, understanding the functions of SmPWOs in *S. moellendorfii* and how they contribute to the acquisition of new traits for land plant evolution are key questions to address in the future.

### PWOs associate with PRC2 since their emergency in lycophytes and have a conserved functions in plant development

A distinct feature of the PWO proteins is the presence of highly intrinsically disordered regions (IDRs) (**Supplementary Figure S2).** IDRs function as protein assemblers, providing large binding interfaces that scaffold multiple partners to facilitate higher-order protein complex formation. Proteins containing IDRs often facilitate both stable and transient protein-protein interactions, play a crucial role in transcription and chromatin regulation, mediate stress responses, and contribute to development (Sun et al. 2013; Salladini et al. 2020; Cermakova and Hodges 2023; Hsiao 2024; Gupta et al. 2025; Miao and Chong 2025). Therefore, we propose that the presence of IDRs in PWO proteins across multiple evolutionary groups may confer conserved PWO functions, such as its ability to interact with other chromatin regulators (Tan et al. 2018; Zheng et al. 2023; Godwin et al. 2024). IDRs are also known to facilitate phase separation by selectively partitioning proteins in the subnuclear space through liquid-liquid phase separation (LLPS), forming different types of nuclear speckles (Cermakova and Hodges 2023). These nuclear bodies (NBs) are often associated with specific genomic loci and exhibit a dynamic nature in response to different environmental stimuli, including acclimation to stress conditions and immune responses in plants (Wang and Gu 2022; Solis-Miranda et al. 2023). As PWOs (AtPWO1, SmPWOa, and SmPWOb) form varying amounts of nuclear speckles/NBs in the nucleoplasm (Fig. 3-6), it will be interesting to determine their possible dynamics in stress conditions and to address a possible role of PWOs in stress responses, as previously suggested (Hohenstatt et al. 2018; Mikulski et al. 2019). It will be of particular interest to determine whether PWOs may confer resistance to environmental challenges imposed especially on land plants.

In Arabidopsis, PWOs interact with key chromatin regulatory complexes involved in gene activation (e.g. histone acetyl transferases and deubiquitinases) or repression (e.g. PRC2) (Hohenstatt et al. 2018; Tan et al. 2018; Zheng et al. 2023; Godwin et al. 2024). The PWO-CLF interaction is conserved in the orthologs SmPWOa and SmCLF (Fig. 7A, 7E). Interestingly, SmPWOa interacts with CLF of *P. patens* (PpCLF) (Fig. 7A, 7E), where PWO proteins are absent. This indicates that PWO emerged to interact with a conserved CLF interaction surface rather than vice versa. Unlike PWOs, most known PWO interactors, such as PRC2 (Huang et al. 2017; Sharaf et al. 2022; de Potter et al. 2023) and UBP5 (Zheng et al. 2023; Godwin et al. 2024), are evolutionarily conserved across the green lineage. Similarly, the PWO1 interacting TRB proteins are also conserved from bryophytes to flowering plants (Kusová et al. 2023; Amiard et al. 2024). This implies that the emergence of PWOs in vascular plants may have triggered a shift in chromatin complex regulation. Interestingly, AtPWO1 exhibits a robust interaction with AtSWN (Fig. 7C, 7F, (Hohenstatt et al. 2018)), while its interaction with basic land plant CLFs (SmCLF, PpCLF) was weaker or insignificant (Fig. 7C, 7F). This specificity might arise from the specialization of higher plant AtPWO1 towards SWN, which emerged only in angiosperms (Huang et al. 2017). One of the key features of PWOs observed upon co-expression of PWO and CLF proteins in *N. benthamiana* was tethering of CLF to PWO-derived speckles (Fig. 4-6). This indicates that PWOs play a crucial role in spatially organizing CLF within the nucleus, possibly affecting its function in chromatin modulation. Together, this suggests that CLF orthologs possess a conserved region for PWO interaction, even in species lacking PWOs. It remains unclear whether this conservation is associated with the presence of other accessory proteins that may substitute for PWOs in *P. patens* or whether other conserved proteins exist across all land plants that interact with CLF through the same interaction surface as PWOs.

In addition to the PWWP domain (Supplementary Fig. 1), we identified a conserved short helical structure (C-motif) located at the C-terminal region in all PWO proteins across plant evolutionary clades (Fig. 2, Supplementary Fig. 4-5). The C-motifs that contain an array of hydrophobic amino acids may facilitate interactions with other proteins or help stabilize the structure (Gibbons et al. 2005; Jernigan et al. 2022). Structural predictions indicate that the PWO C-motif may interact with CLF in different species throughout evolution, suggesting that the C-motif may serve as a conserved component for CLF binding. Accordingly, deletion of the C-motif significantly reduces the AtPWO1-AtSWN interaction and the interaction of SmPWOa with all tested CLF orthologs (Fig. 8). Therefore, we propose that the C-motif of the PWO proteins plays an important role in mediating the evolutionarily conserved interaction between PWOs and the PRC2 catalytic subunit.

Overall, we have shown that the PWO-PRC2 interaction is evolutionarily conserved (Fig. 7A, B, E, F). However, the biological relevance of PWO interaction with PRC2 throughout evolution remains unresolved. Recent evidence demonstrated that PWO1 binds to the boundaries of H3K27me3-CDs and plays a crucial role in maintaining these boundaries. This function contributes to the 3D nuclear organization and the repressed state of these CDs (Yang et al. 2024). Although the impact of the PWO1-PRC2 interaction on this activity is still unknown, it has been proposed that PWOs could help to phase-separate PRC2 to reinforce its CD boundaries (Yang et al. 2024). Hence, it is tempting to speculate that the appearance of PWOs contributed to the relocation of PRC2 and H3K27me3, a hypothesis that we aim to explore in future research.

Despite an evolutionary separation of 429 million years (Kumar et al. 2017), the partial complementation of the Arabidopsis *pwo1;pwo2* double mutant by SmPWOs suggests a certain degree of functional conservation. Hence, SmPWOs seem to retain key molecular functions that contribute to plant development, gene regulation and nuclear organization (Fig. 9, Supplementary Fig. 16). On the other hand, the incomplete rescue of the *pwo* double mutant phenotype may be connected to the low level of conservation of the IDR-containing region of PWOs. Differences in these regions are likely to significantly influence PWO functions, as most PWO1 interactors (ARIDs, EPCR1, TRB1-2, and UBP5) primarily bind to the IDRs in Arabidopsis (Zheng et al. 2023). Therefore, it may be possible that SmPWOs cannot establish a proper interaction network in Arabidopsis. It will be intriguing to further investigate the level of conservation of the PWO-interacting protein network across evolution and to explore how PWOs contribute to establishing novel interactions or to strengthening existing ones.

## Material and Methods

### Plant materials and growth conditions

All the lines used in the current study were in Columbia (Col-0) ecotype. *pwo1-1* (Sail_342_C09; At3g03140) single mutant was crossed with *pwo2-2* single mutant (Salk 136093; At1g51745) to obtain the *pwo1-1;pwo2-2* double mutant (Hohenstatt et al. 2018). The homozygous mutants were confirmed by PCR-based genotyping using primers P1–P8 (Supplementary Table 8). Subsequently, the *Agrobacterium tumefaciens* (GV3101) harboring the pMDC85-*2X35S_pro_::SmPWOa-GFP* and pMDC85-*2X35S_pro_::SmPWOb-GFP* constructs (Curtis and Grossniklaus 2003a) were transformed into *pwo1-1;pwo2-2* double mutants using the floral dip transformation method (Karimi et al. 2007). The Col-0, *pwo1-1;pwo2-2* double mutant, *2X35S_pro_::SmPWOa-GFP; pwo1-1;pwo2-2* and *2X35S_pro_::SmPWOb-GFP*; *pwo1-1;pwo2-2* were grown in pots containing peat, vermiculite, and perlite in ratio of 5:1:1. The plants were grown under long day (LD) (16/8 light/dark) conditions at 20°C/18°C temperature regime under fluorescent lamps at 120 μmol m−2 s−1. At least 10 plants per genotype were used for flowering time analyses. Flowering time was scored by counting the number of days when the first flower opened. *In vitro* studies, such as root measurement, were carried out on ½ MS medium supplemented with 0.8% agar and 0.5% sucrose. Seeds were first stratified at 4°C for 3 days for breaking dormancy and synchronizing germination. Three-week-old *N. benthamiana* plants were used for agroinfiltration for transient gene expression.

### Homolog identification and phylogenetic analyses

The complete predicted proteome sequences of studies organisms (Supplementary Table 1) were obtained from the NCBI GeneBank (https://ncbi.nlm.nih.gov), UniProt-Proteomes database (https://www.uniprot.org), and JGI (http://genome.jgi.doe.gov). The 1000 Plants project (OneKP) database (http://www.onekp.com) was an additional source for predicted proteome sequences inferred from transcriptomic data. The *A. thaliana* PWO1 (At3g03140) reference amino acid sequence was used to search for all predicted proteomes using the Hidden Markov model (HMM)-based tool jackhammer (Johnson et al. 2010). Evolutionary genealogy of genes: non-supervised Orthologous Groups (eggNOG) mapper was used for hierarchical resolution of orthology assignments (Huerta-Cepas et al. 2019b). Only eggNOG-hits with ENOG5028MA3 and ENOG5028N12 were selected. Finally, the SMART and Pfam databases were employed to identify conserved domains present in PWOs from different organisms (Letunic and Bork 2018; El-Gebali et al. 2019), both SMART and Pfam databases were merged, redundant domains were filtered-out and the Hidden Markov model (HMM)-based tool hmmscan (https://github.com/EddyRivasLab/hmmer) was used to scan domains architecture. All identified ortholog sequences were aligned using MAFFT software (Katoh and Standley 2013) and ambiguously aligned regions were excluded for further analysis using trimAl software (Capella-Gutiérrez et al. 2009). Alignments were tested using ProtTest v3 (Darriba et al. 2011) to choose an appropriate model for nucleotide substitution. Two separated Maximum likelihood (ML) phylogenetic trees were computed using RAxML-NG (Kozlov et al. 2019) and IQ-TREE 2 (Minh et al. 2020) software. ML analyses were performed using 1000 bootstrap replicates. The supporting values from both software were noted on the ML-rooted tree. The phylogenetic tree was rooted with Lycopodiopsida sequences, which were considered as the emerging point of the PWO family.

### Cloning and Plasmid Preparation

The full-length coding sequences of SmPWOa (P9-P10), SmPWOb (P11-P12), SmCLF (P13- P14), and PpCLF (P15-P16) were amplified (Supplementary Table S8) from a total cDNA library prepared from *S. moellendorffii* and *P. patens*. They were then cloned into pJET1.2/blunt (CloneJET™ PCR Cloning Kit, Thermo Scientific #K1231, #K1232). Subsequently, all pJET1.2 clones were validated by colony PCR, plasmid digestion, and Sanger sequencing, and used as templates for further cloning. Similarly, *SmPWOa* (P17-P18), *SmPWOb* (P19-P20), *SmCLF* (P21-P22), and *PpCLF* (P23-P24) cDNAs were cloned into pDONR221 (Supplementary Table S8), followed by subcloning into destination vectors using the Gateway™ technology (Invitrogen) (pGBKT7-GW, pGADT7-GW, pMDC7:i35S-GFP, pMDC7:i35S-mCherry, pMDC85-GFP) (Bleckmann et al. 2009; Lu et al. 2009). Additionally, the *ARR1* (AT3G16857) and *ARR2* (AT4G16110) cDNAs were amplified using primers P25- P26 (*ARR1*) and P27-P28 (*ARR2*). The resulting fragments were recombined into the donor vector pDONR™/Zeo, following the manufacturer’s protocol for the Gateway cloning system (Thermo Fisher Scientific), and subsequently subcloned into the pB7WGR2 destination vector (Karimi et al. 2007) (Supplementary Table 8). In-Fusion Snap Assembly (Takara) was used for the *SmPWOa* (P29-P30) and *SmPWOb* (P31-P32) cDNAs cloning into pMDC85 vector (Curtis and Grossniklaus 2003b) (Supplementary Table 8). *A. thaliana* gene clones, including pMDC7:i35S-AtPWO1-GFP, pMDC7:i35S-AtPWO1-mCherry, pGBKT7-AtCLFΔSET, and pGBKT7-AtSWNΔSET, were generated before (Hohenstatt et al. 2018; Mikulski et al. 2019). For C-motif deletion, mutated primers were used to amplify *AtPWO1* (P33-P34) and *SmPWOa* (P35-P36) cDNAs without the C-motif, followed by cloning into pDONR221 vectors and subsequent subcloning into pGBKT7-GW, pGADT7-GW, pMDC7:i35S-GFP, and pMDC7:i35S-mCherry.

### Yeast two hybrid assays

For yeast two-hybrid (Y2H) assays, the *S. cerevisiae* strain AH109 was used to co-transform both pGADT7-Gal4-AD and pGBKT7-Gal4-BD constructs in various combinations, following the protocol outlined in the Yeast Protocols Handbook (version no. 325, PR973283 21; Clontech). Transformed yeast cells were selected on synthetically defined (SD) medium lacking Leucine (L) and Tryptophan (T), supplemented with Adenine (A) and Histidine (H); (SD/-Trp/- Leu/+Ade/+His) at 30°C for 2-3 days. Interaction assays were conducted on SD medium with both low stringency (SD/-Trp/-Leu/+Ade/-His) and high stringency selection plates (SD/-Trp/- Leu/-Ade/-His). Co-transformation involving empty Gal4-AD and Gal4-BD constructs, as well as empty bait with prey constructs served as controls.

### Subcellular localization and FRET assays

*Agrobacterium tumefaciens* (GV3101) was transformed with pMDC7 (estradiol-inducible) and pMDC85 clones tagged with GFP/mCherry employing the Freeze-Thaw method (Weigel and Glazebrook 2006). After transformation, the colonies were allowed to grow for 2 days on yeast extract broth (YEB) (Sigma-Aldrich, Y1625) medium plates. Transformed colonies were then cultured in YEB medium overnight, followed by subculture for an additional 3-4 hours. Subsequently, the cultures were harvested and resuspended in infiltration medium (containing 10 mM MgCl_2_ (Sigma-Aldrich, M8266), 10 mM MES (Sigma-Aldrich, M3671) [pH 5.6], and 200 μM acetosyringone (Sigma-Aldrich, D134406) to an optical density (OD) of 0.6 and left at room temperature for 1 hour. Bacteria were then infiltrated into 3–4-week-old *N. benthamiana* leaves using 2-mL syringes and left for 48 hours. The induction buffer (Beta-estradiol (Sigma-Aldrich, E2758) solution in 0.1% [v/v] TWEEN-20 (Sigma-Aldrich, P1379) was sprayed on the abaxial side of the leaves. At 6-8 hours post-induction, infiltrated leaves were subjected to confocal microscopy (Olympus FV3000). Image analysis was carried out using FIJI/ImageJ software (Schindelin et al. 2012) (National Institutes of Health).

For FLIM-FRET experiments, all plasmids were transiently expressed in *N. tabacum* (SR1 Petit Havana) leaf epidermal cells using the infiltration procedures described by (Voinnet et al. 2000). Confocal microscopy was performed using a laser scanning confocal imaging microscope Zeiss LSM 780 AxioObserver equipped with an external In Tune laser (488-640 nm, < 3nm width, pulsed at 40 MHz, 1.5 mW) and a C-Apochromat 63x water objective with NA 1.2. FLIM-FRET data acquisition was conducted using an HPM-100-40 Hybrid Detector from Becker and Hickl GmbH, which includes Simple-Tau 150N (Compact TCSPC system based on SPC-150N) with a DCC-100 detector controller for photon counting. Zen 2.3 light version from Zeiss was utilized for processing confocal images. The acquisition and analysis of FLIM data involved the utilization of SPCM 64 version 9.8 and SPCImage version 7.3 from Becker and Hickl GmbH, respectively. A multiexponential decay model was employed for fitting the data. For the prediction of NLSs in PWO proteins, DeepLoc-2.0 (Thumuluri et al. 2022) was utilized. This tool is designed to predict the subcellular localization of proteins based on their amino acid sequences. Typically, threshold values closer to 1 indicate higher confidence in the prediction, while values closer to 0 indicate lower confidence.

### Alphafold2 (AF2) analyses

Protein structures and interaction interface were predicted using UCSF ChimeraX (Pettersen et al. 2021; Meng et al. 2023), utilizing its built-in tool for structure prediction via the AF2 server (Evans et al. 2021; Jumper et al. 2021). Each AF2 predicted file produced results with pdb and json files, graph for sequence homology coverage, pLDDT plot and 5 rank model PEA plots. The PDB files contain the structures predicted by ColabFold, while the .json file contains the individual scores for each amino acid in the predicted structure. Moreover, PAE plots shows the uncertainty in the distance prediction between individual protein domains represented with color code ranging from blue (0–15 Å) to red (15–30 Å). For the analysis, the pLDDT, PAE and ipTM have been taken into consideration and results were visualized using ChimeraX (Pettersen et al. 2021; Meng et al. 2023).

### Gene Expression analyses

For expression analyses, total RNA was extracted from 10-day-old seedlings grown on ½ MS media at ZT16 using Thermo Scientific kit according to manufacturer’s instructions. 2μg of total RNA was used to synthesize first strand cDNA using RevertAid First strand cDNA synthesis kit by Thermo Scientific. The PCR amplification from cDNA was carried out using Takyon™ No ROX SYBR 2X MasterMix blue dT^1^TP by Eurogentec using the Primers P37- P44 (Supplementary Table 8). *TIP41* (*TAP42 INTERACTING PROTEIN OF 41 kDa*) (P45-P46) was used as an internal control (Supplementary Table 8). Error bars show standard deviation of three biological replicates.

### Nuclear Morphology Analyses

Nuclei isolation was conducted on 10-day-old seedlings of Col-0 (WT), *pwo1-1;pwo2-2* double mutants, *2X35S_pro_::SmPWOa-GFP; pwo1-1;pwo2-2* and *2X35S_pro_::SmPWOb-GFP*; *pwo1-1;pwo2-2* genotypes following the protocol described by Kalyanikrishna et al. 2020 (Kalyanikrishna et al. 2020). Fluorescence imaging of the isolated nuclei stained with DAPI was performed using a Leica SP8 confocal microscope, with a 405 nm diode laser for DAPI excitation. Z-stacks of 40-60 images were acquired with a step size of 0.25 µm, maintaining consistent imaging settings across genotypes. Nuclear area and circularity index were measured using Fiji/ImageJ (Schindelin et al. 2012), with z-stacks analyzed according to Kalyanikrishna et al. 2020 (Kalyanikrishna et al. 2020). More than 100 nuclei per genotype were analyzed, and for each genotype, three technical replicates and two biological replicates were used to conduct the experiment.

### Reference Accession Numbers

Sequence information for the genes used in this article can be found in GenBank/EMBL data libraries under accession numbers At3g03140 (*PWO1*), At4g02020 (*AtSWN*), At2g23380 (*AtCLF), XP_024515258.1* (*SmPWOa*)*, XP_024540635.1* (*SmPWOb*)*, XP_002987466.1* (*SmCLF*), AB472766*.1* (*PpCLF*), AT2G37620 (*ACT1*), AT1G62360 (*STM*), AT5G59980 (*GAF1*), AT1G32310 (*SAMBA*), AT4G34270 (*TIP41*).

## Funding

This work was supported by ERC-CZ (ERC200961901) awarded to IM, the Marie Skłodowska-Curie Individual Fellowship 2020 (MSCA-IF 2020; ID: 101026898) awarded to AK, and Research Ireland Frontiers for ^2^the Future Programme (20/FFP-P/8693) awarded to SF and SH. The work of PPS and AKu were supported by the Czech Science Foundation [21-15841S], and further, PPS was supported by the Ministry of Education, Youth and Sports of the Czech Republic (INTER-COST LUC24056). For JS, JH, PPS and AKu, the work was additionally supported from the project TowArds Next GENeration Crops, reg. no. CZ.02.01.01/00/22_008/0004581 of the ERDF Programme Johannes Amos Comenius.

## Acknowledgements

We thank Dr. Alexey Bondar, Research Scientist at the Laboratory of Microscopy and Histology (LMH), Biology Centre, for his invaluable assistance with confocal microscopy and image analysis. We also thank Biswajit Ghosh, Institute of Biology, Freie Universität Berlin, for his assistance with the nuclear morphology experiments. We thank Prof. Volker Knoop, University of Bonn, DE for donating *S. moellendorffii* plant material and Jesús López-Corrales for originally cloning the *SmPWO* cDNAs.

## Author contributions

AK, IM, and SF conceptualized the experimental approach and designed the methodology. AK performed most of the molecular experiments, including construct generation, protein localization, transgenic plant generation, confocal microscopy analyses, and data visualization, as well as manuscript writing and revision. SH conducted genotyping and phenotyping of transgenic lines as well as gene expression analyses for the transgene complementation experiments. AS and AK conducted the phylogenetic analyses. AKu and JS carried out the FLIM-FRET experiments. CJ and AK performed the nuclear morphology analysis. AK and MR performed and interpreted AF modeling. All authors contributed to the interpretation of the results. AK, IM, and SF wrote and revised the manuscript.

---

**Supplementary Figure 1.** Multiple Sequence Alignment (MSA) of the conserved PWWP domain across plant evolution. The MSA was generated using MAFFT v7 with default parameters, and ESPript v3 (Robert and Gouet 2014) was used to highlight conserved regions, secondary structure elements, and other sequence features. (A separate alignment file is provided in PDF format.)

**Supplementary Figure 2.**
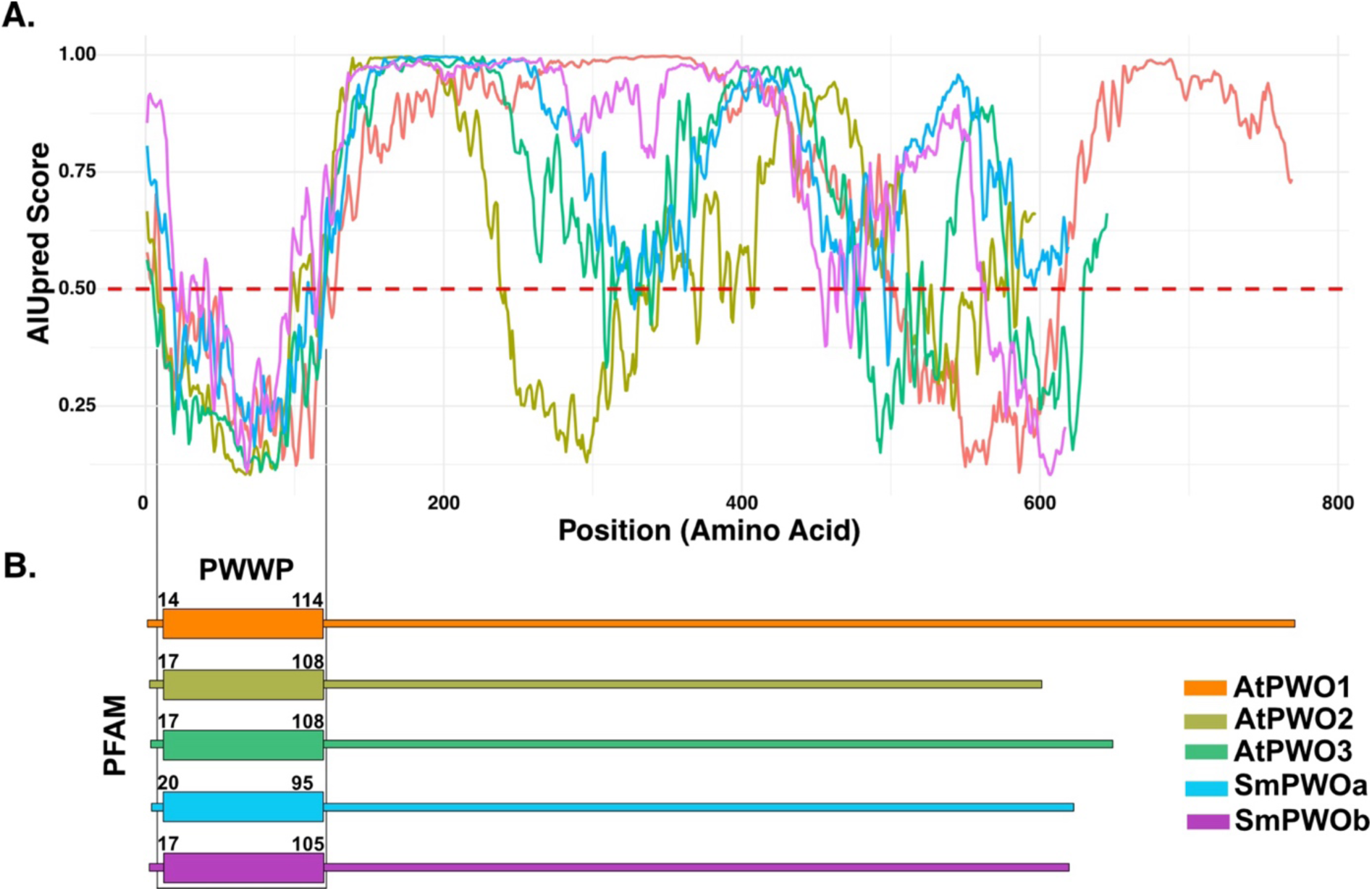
Bioinformatic analysis of Intrinsically Disordered Regions (IDRs) in PWO proteins. **A.** Prediction of intrinsically disordered regions in full-length PWO proteins (AtPWO1-3, SmPWOa-b) using Artificial Intelligence-based Unstructured Region Prediction (AIUPred) (Erdős and Dosztányi 2024). Protein regions with an AIUPred score greater than 0.5 are considered disordered. **B.** Position of the PWWP domain as predicted by PFAM. The aa positions marking the start and end of the PWWP domain are indicated.

**Supplementary Figure 3.**
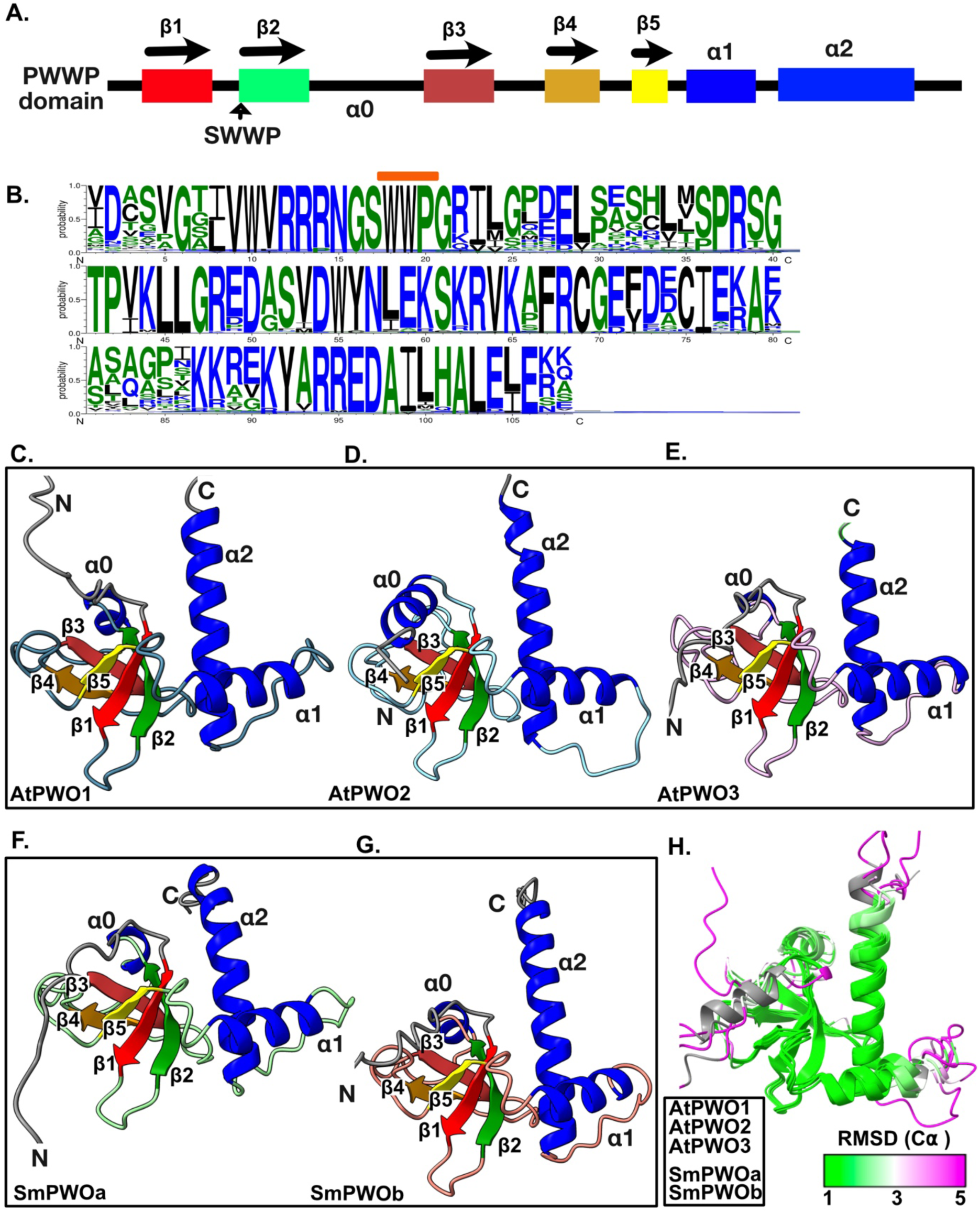
AlphaFold2 structure prediction of *A. thaliana* and *S. moellendorfii* PWO protein PWWP domains. **A.** Schematic representation of the PWWP domain, displaying the positioning of β strands (β1: red, β2: green, β3: brown, β4: orange, β5: yellow) and α-helices (α1, α2: blue), with α0 (unmarked) being more variable. The position of SWWP motif at the beginning of the β2 strand is indicated by an arrow. **B**. Consensus sequence plot showing the PWWP domain for all four clades (Clade I-IV), with the orange line indicating the conserved SWWP motif. **C-G.** Structure prediction of the PWWP domain of *A. thaliana* PWO orthologs (AtPWO1, AtPWO2, AtPWO3) (**C-E)** and *S. moellendorffii* PWO orthologs (SmPWOa, SmPWOb) **(F-G)**, showing the positioning of secondary structures (β-strands and α-helices) as illustrated in panel A. **H.** Superposition of the PWWP domains of Arabidopsis and S. *moellendorffii* PWO orthologs. The RMSD (Cα) value and color range is displayed, with green indicating values of 1 Å or less, signifying high structural similarity.

**Supplementary Figure 4:**
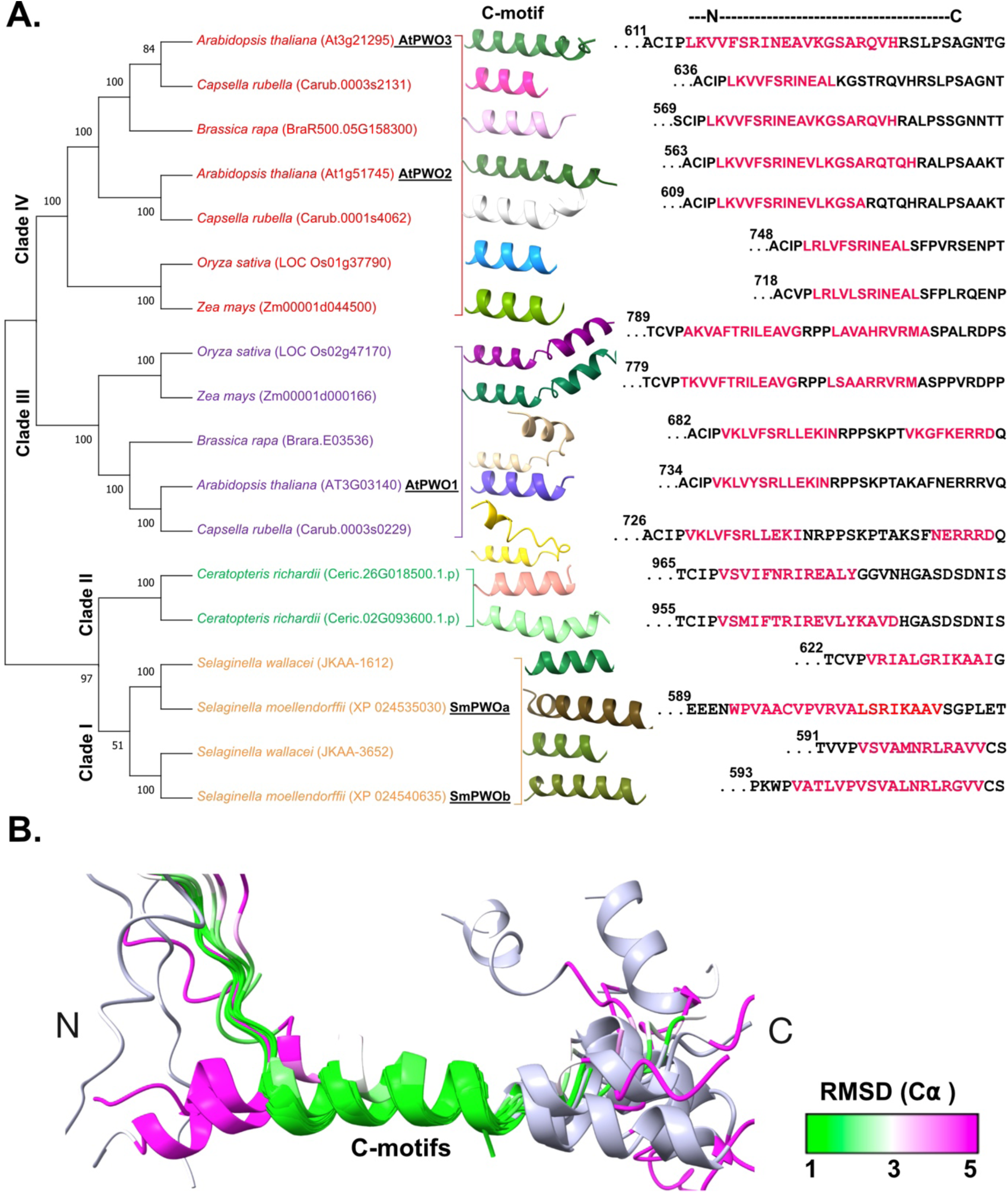
AlphaFold2-based structure prediction of the C-motif in PWO proteins across representative species with PWO clades. **A.** The maximum likelihood phylogenetic tree of PWO protein orthologs from 18 selected representative species across the four PWO clades (Clade I, Clade II, Clade III, and Clade IV) depicts the predicted C-motif structures and C-motif sequences, highlighted in pink. **B.** Superposition of all predicted C-motifs in panel 3A is shown, with the RMSD (Cα) color range values displayed. Green represents values of 1 Å or less, indicating high structural similarity.

**Supplementary Figure 5.**
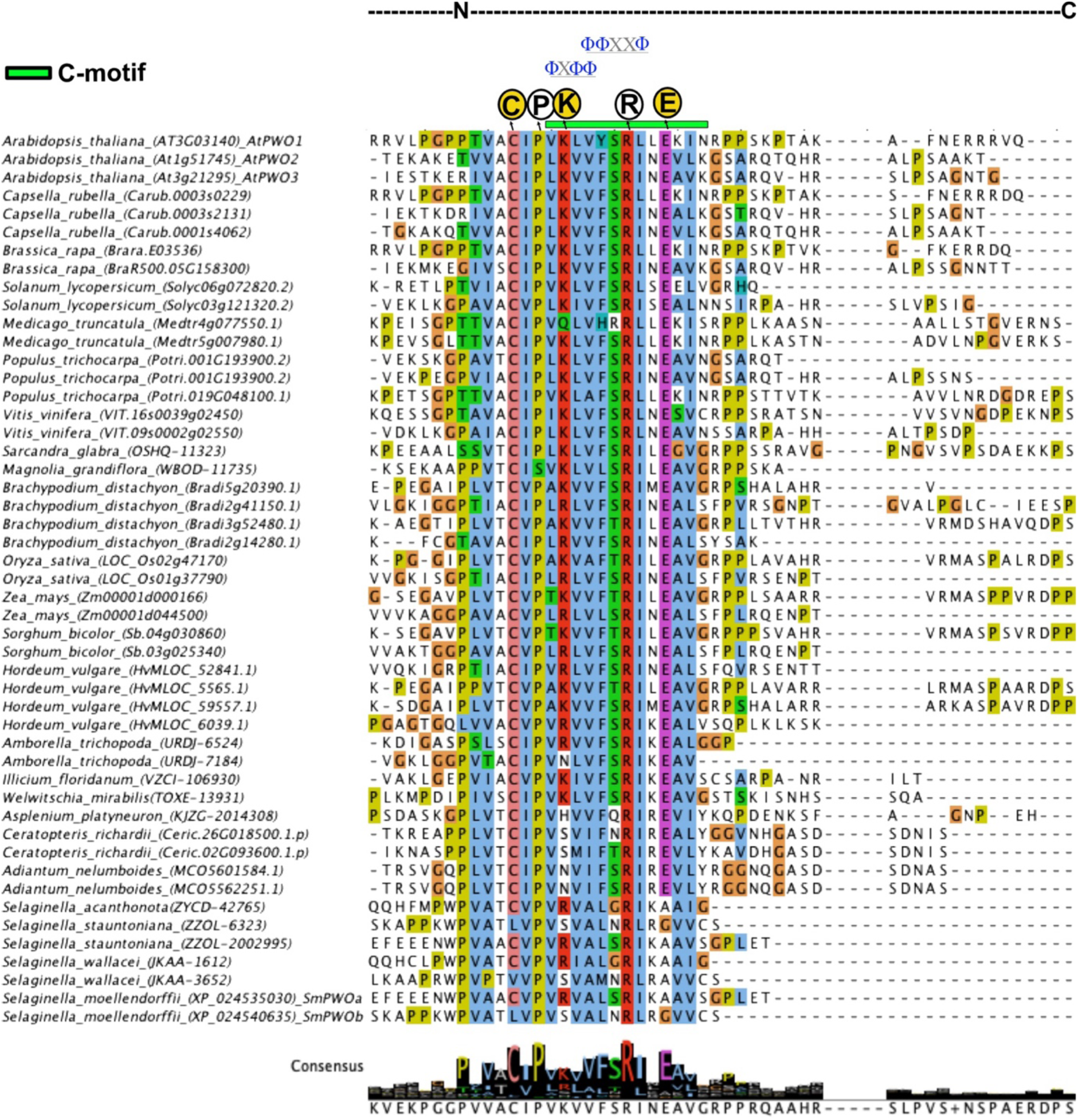
Amino acid multiple sequence alignments of PWO C-terminal region including C-motif. Sequence alignment was generated using ClustalO, displayed using JalView employing the Clustal color mode. The consensus tracks are calculated by JalView (Waterhouse et al. 2009). The sequence coverage is represented by a black bar. Φ indicates the hydrophobic amino acid (aa) residues. The conserved aa residues are circled black, while a yellow-filled circle indicates lower level of conservation in basal land plants.

**Supplementary Figure 6.**
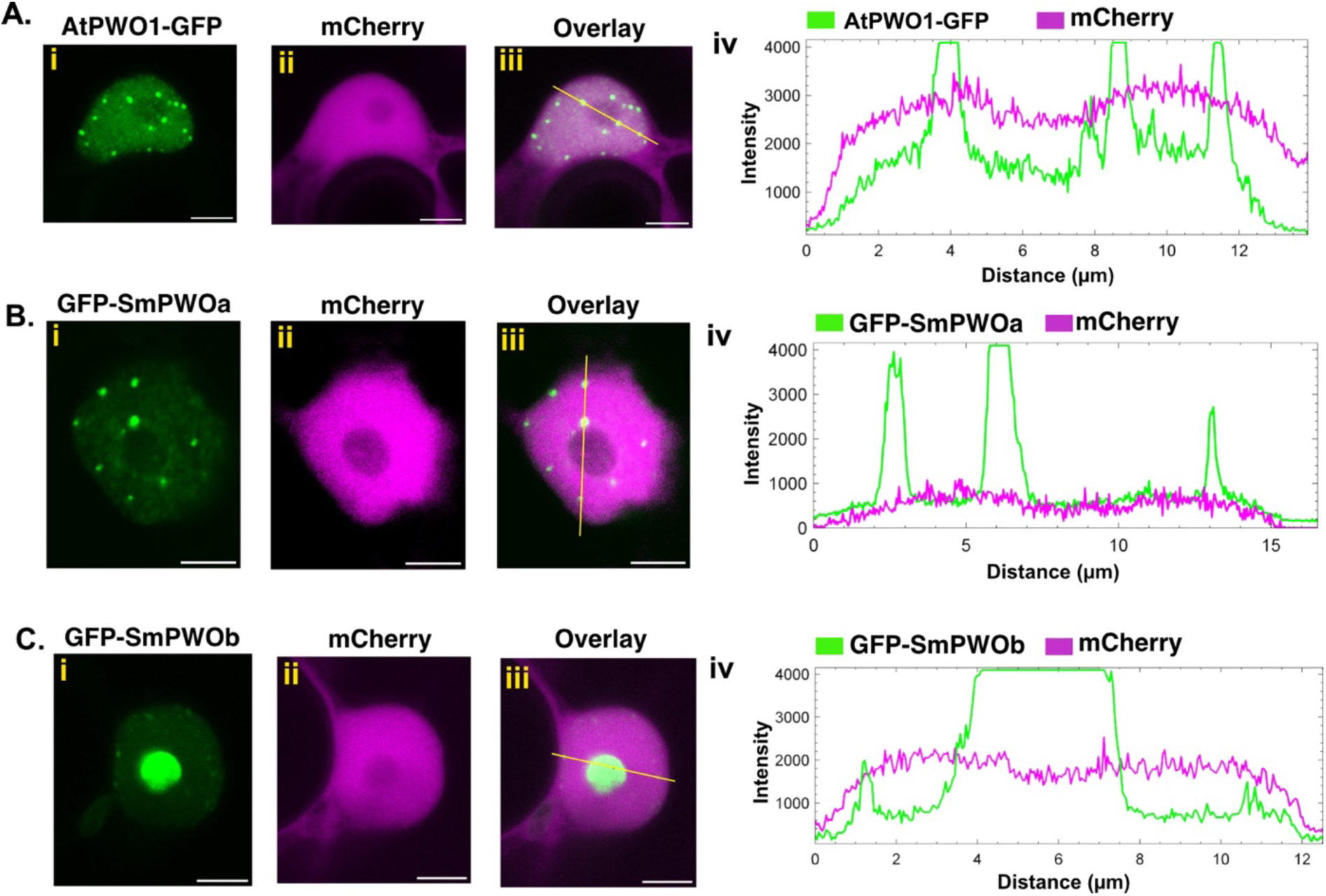
Co-infiltration of PWOs (AtPWO1, SmPWOa, SmPWOb) with empty mCherry. Representative confocal microscopy images of nuclei of *N. benthamiana* leaf cells co-infiltrated with *i35S_pro_::mCherry* and **Ai-iii.** *i35S_pro_::AtPWO1-GFP*, **Bi-iii.** *i35S_pro_::GFP-SmPWOa*, and **Ci-iii.** *i35S_pro_::GFP-SmPWOb*. **A-Civ:** Profiles of mCherry and GFP fluorescence intensities along the yellow line in **A-Ciii**. Scale bar = 5 µm.

**Supplementary Figure 7.**
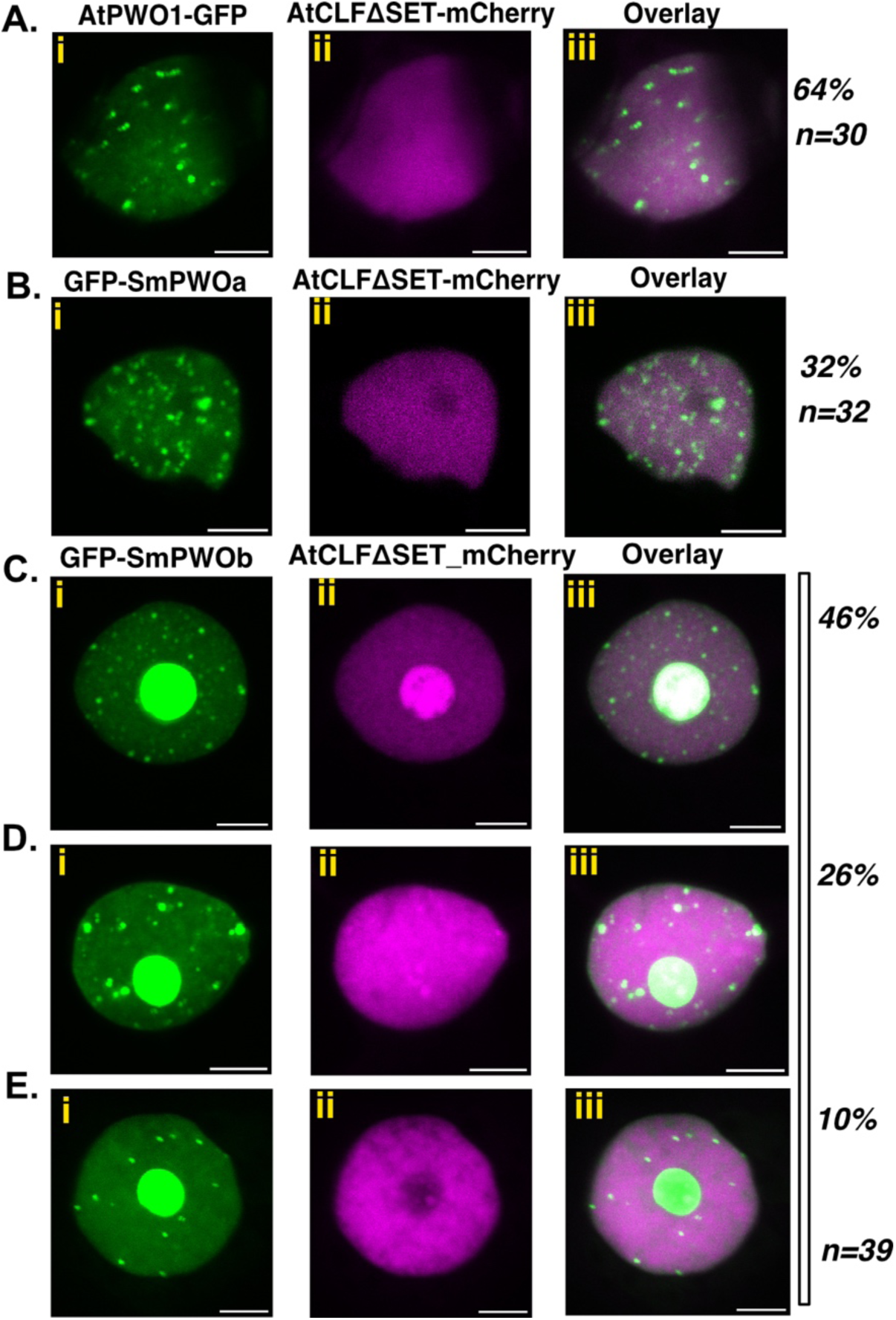
Colocalization of SmPWOs and AtWPO1 with AtCLF in *N. benthamiana*. Representative confocal microscopy images showing nuclei of *N. benthamiana* leaf cells infiltrated with *i35S_pro_::mCherry-AtCLFΔSET* and **A.** *i35S_pro_::AtPWO1-GFP*, **B.** *i35S_pro_::GFP- SmPWOa*, **C-D.** *i35S_pro_::GFP-SmPWOb*. The percentage of nuclei with the observed pattern related to the total number of analyzed nuclei (n) is indicated on the right. ‘ΔSET’ denotes the deletion of the SET domain. Scale bar = 5 µm

**Supplementary Figure 8.**
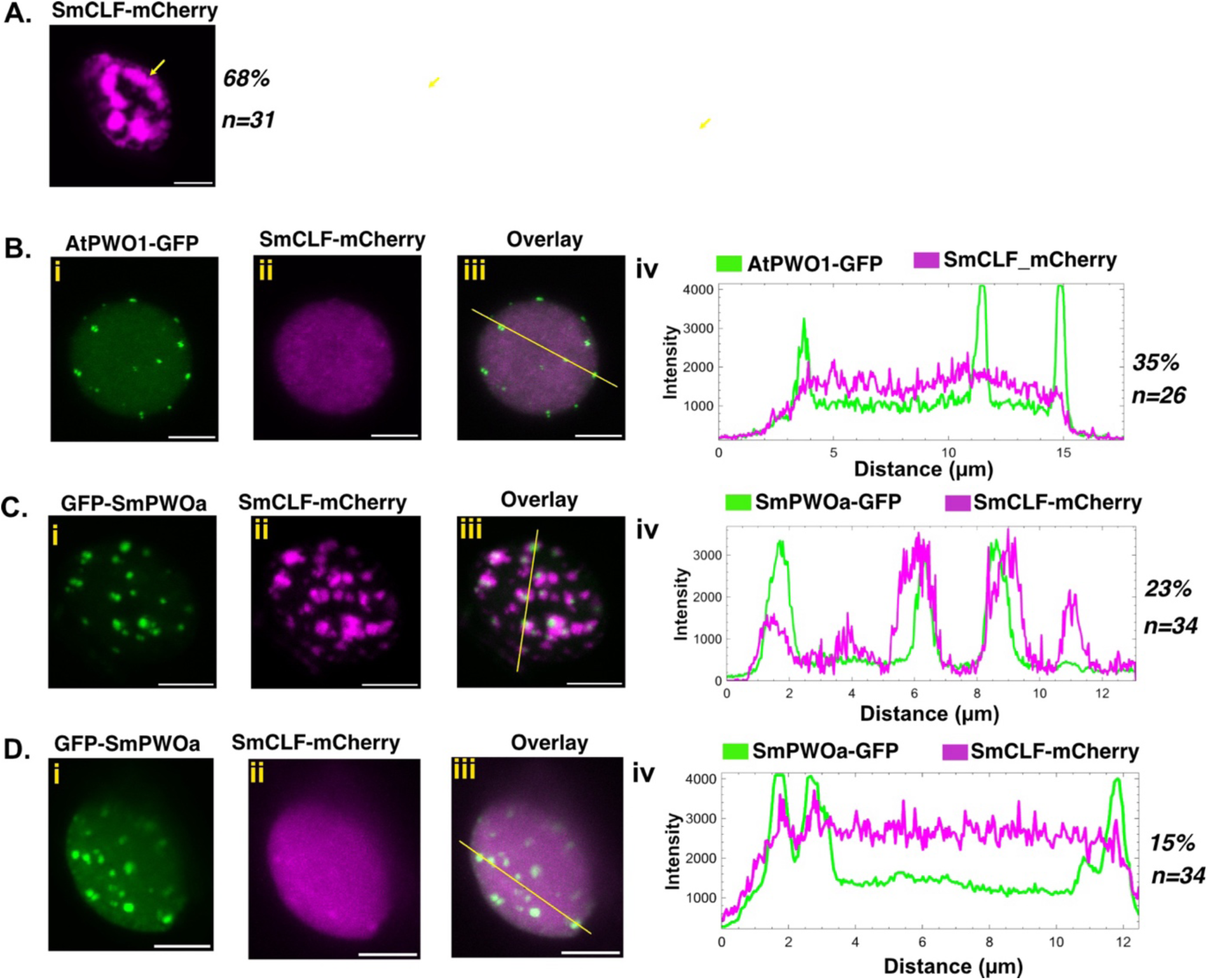
Localization and colocalization of SmCLF with AtPWO1 and SmPWOa in *N. benthamiana*. Representative confocal microscopy images showing nuclei of *N. benthamiana* leaf cells infiltrated with **A.** *i35S_pro_::SmCLF-GFP*, co-infiltrated with **B.i-iii.** *i35S_pro_::AtPWO1-GFP* and *i35Spro::SmCLF-GFP.* **C-D.i-iii.** *i35S_pro_::GFP-SmPWOa* and *i35S_pro_::SmCLF-GFP*. **B-D.iv:** Profiles of GFP and mCherry fluorescence intensities along the yellow line shown in B-D. iii. The percentage of nuclei with the observed pattern related to the total number of analyzed nuclei (n) is indicated on the right. Arrow indicates larger nuclear patches. Scale bar = 5 µm.

**Supplementary Figure 9.**
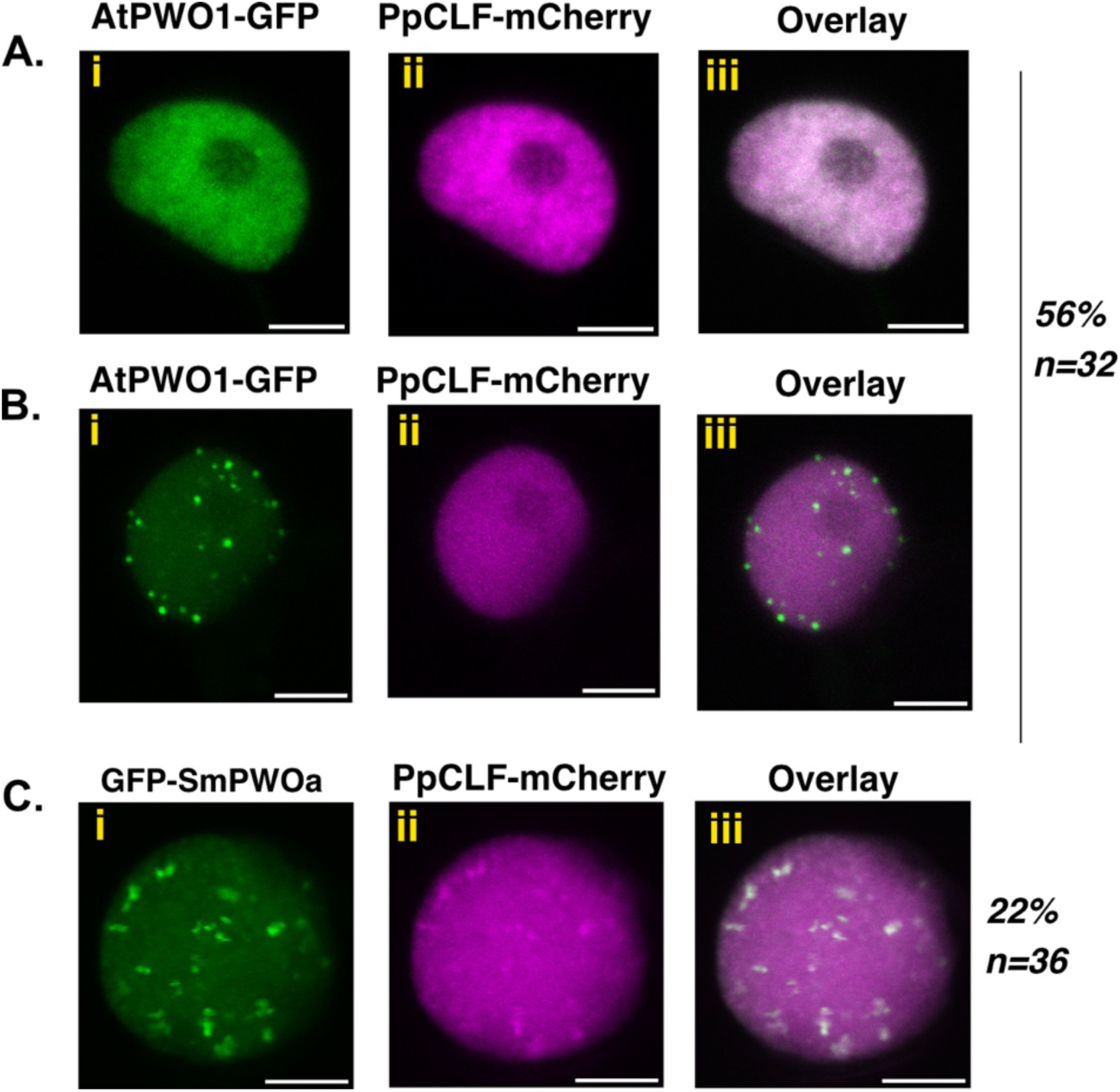
Colocalization of AtPWO1 and SmPWOa with *P. patens* (Pp)CLF in *N. benthamiana*. Representative confocal microscopy images showing nuclei of *N. benthamiana* leaf cells co-infiltrated with **A-B.i-iii.** *i35S_pro_::AtPWO1-GFP* and *i35S_pro_::PpCLF-mCherry*. **C.i-iii.** *i35S_pro_::GFP-SmPWOa* and *i35S_pro_::PpCLF-mCherry*. The percentage of nuclei with the observed pattern related to the total number of analyzed nuclei (n) is indicated on the right. Scale bar = 5 µm.

**Supplementary Figure 10.**
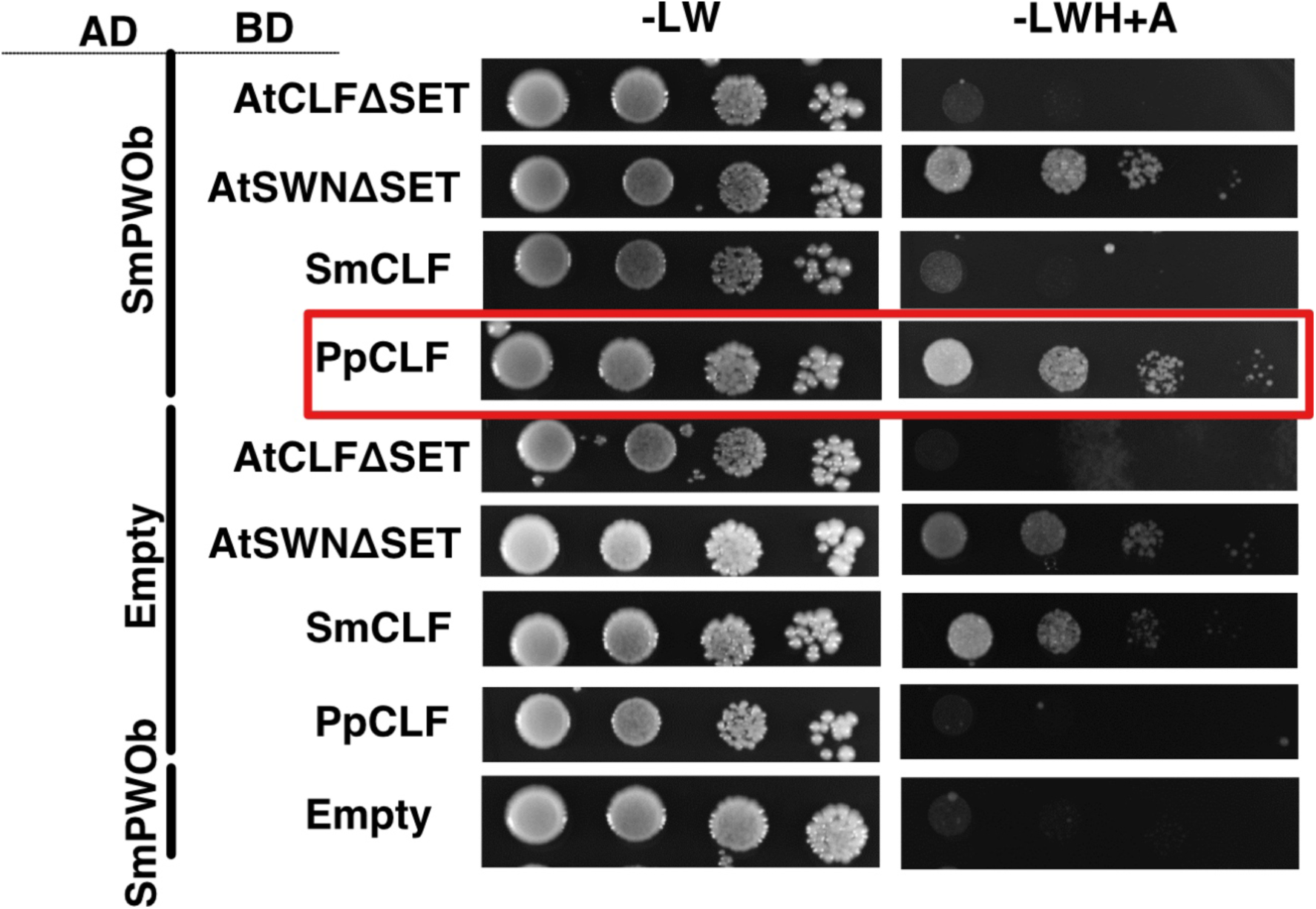
Yeast two-hybrid assays for SmPWOb and PRC2 catalytic subunit interactions. Interactions between SmPWOb and AtCLF, AtSWN, PpCLF, and SmCLF were tested on low-stringency medium lacking leucine, tryptophan, histidine, and containing adenine (−LWH + A). The red box highlights a weak interaction between SmPWOb and PpCLF.

**Supplementary Figure 11.**
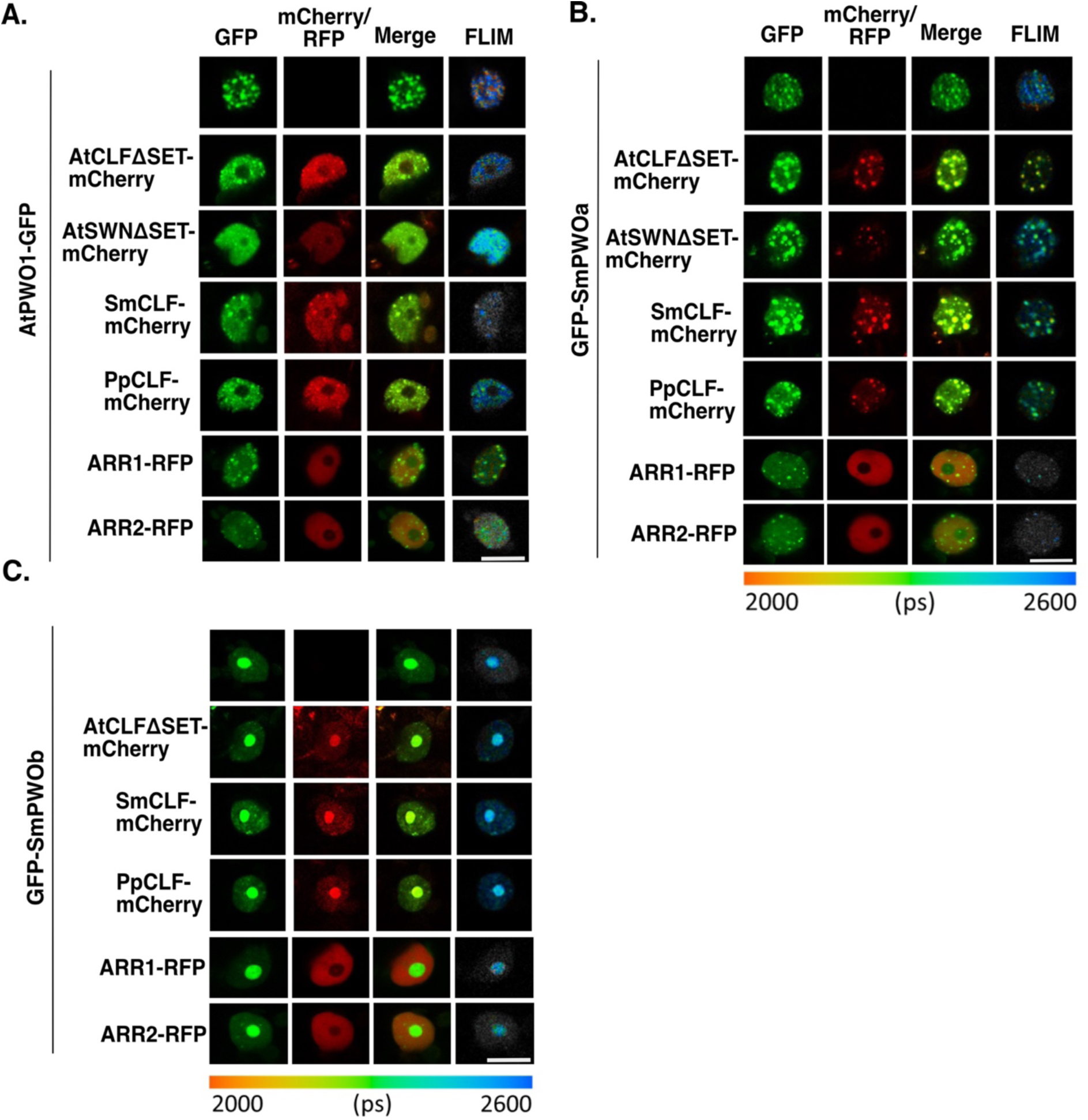
FLIM-FRET confocal microscopy images for PWOs and CLF orthologs. **A.** AtPWO1, **B.** SmPWOa, and **C.** SmPWOb interactions with the PRC2 catalytic subunits from Arabidopsis (AtCLF, AtSWN), *S. moellendorffii* (SmCLF), and *P. patens* (PpCLF). Scale bar = 10 μm. The FLIM-FRET data are displayed using a lifetime Look-Up Table (LUT). ARR1/2- RFP serves as negative controls.

**Supplementary Figure 12.**
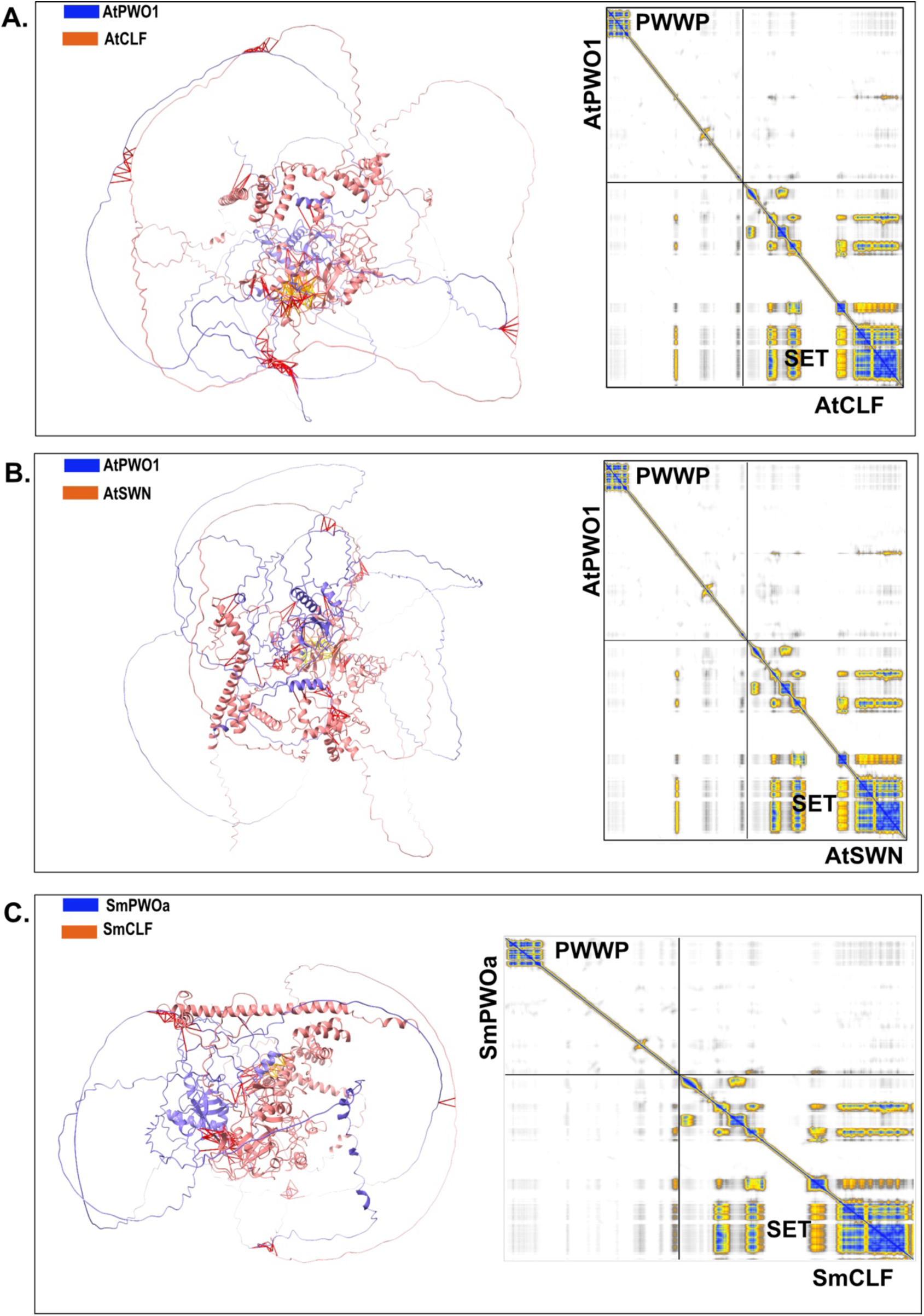
AlphaFold2-Multimer (AF2-M)-based prediction of interaction surfaces between PWOs and CLF or SWN in *A. thaliana* and *S. moellendorffii.* AF2-M prediction of the interaction surface between full-length **A.** AtPWO1 and AtCLF, **B.** AtPWO1 and AtSWN, **C.** SmPWOa and SmCLF, along with the Predicted Aligned Error (PEA) plot for rank 1, for each of the combinations.

**Supplementary Figure 13.**
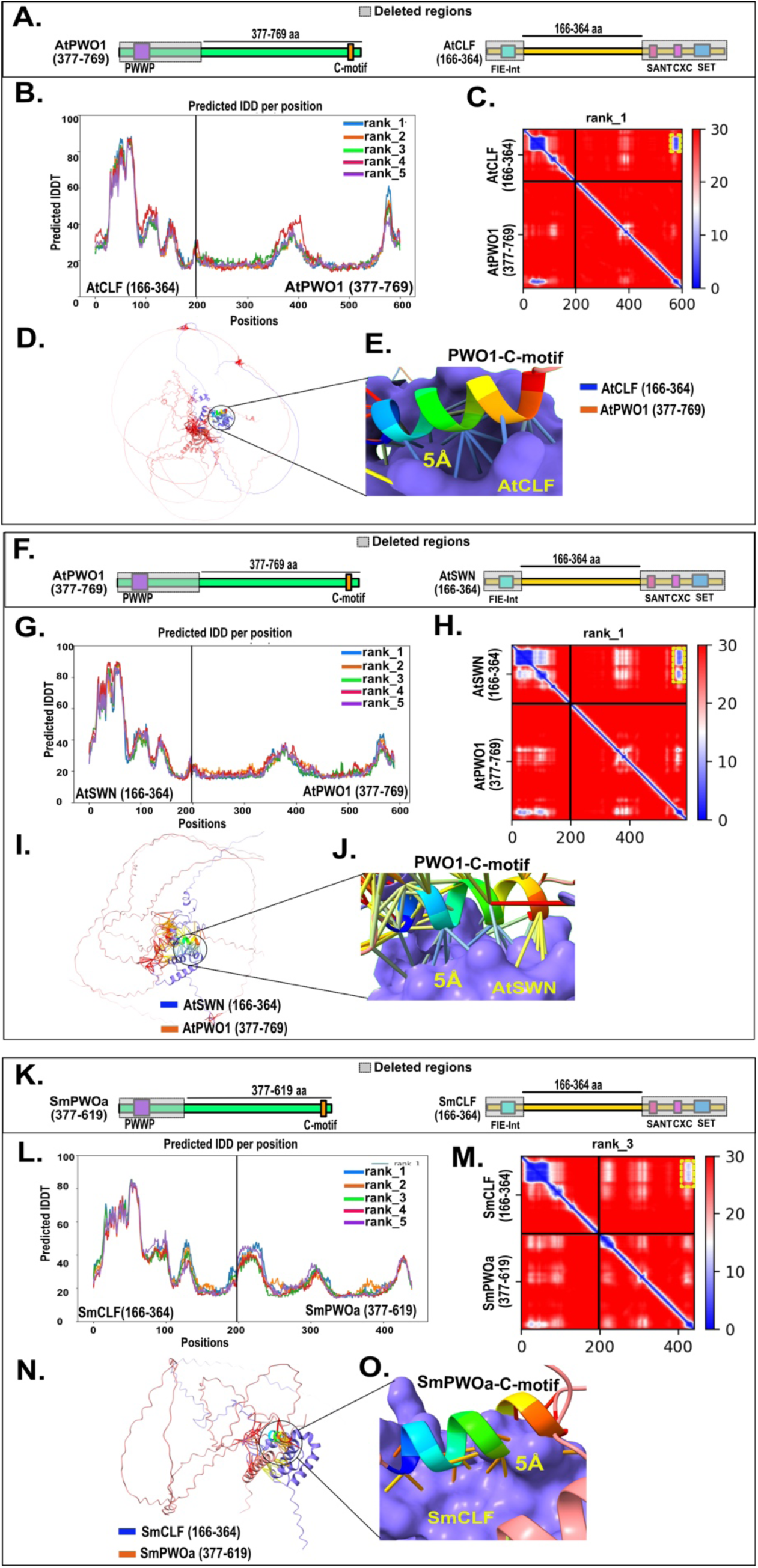
AF2-M-based prediction of the interaction surface between truncated PWOs and CLF/SWN. **A.** Schematic diagram showing the truncated fragments of AtPWO1 and AtCLF used for interaction prediction. **B.** pLDTT graph showing values for five models for AtPWO1 (377-769 aa) interaction with AtCLF (166-364 aa). **C.** PAE plot displaying the rank 1 models from panel B with noticeable local interactions in PAE maps, outlined by yellow dotted box. **D.** 3D view of the predicted AtPWO1–AtCLF interaction from panel B rank 1 with, **E.** Zoom-in view of the interaction interface, highlighting the interaction between the AtPWO1 C-motif (rainbow-colored) and AtCLF (blue surface). **F.** Schematic diagram showing the truncated fragments of AtPWO1 and AtSWN used for interaction prediction**. G.** pLDTT graph showing values for five models for AtPWO1 (377-769 aa) interaction with AtSWN (166-364 aa). **H.** PEA plot displaying the rank 1 models from panel G with local interactions in PAE maps, outlined by yellow dotted box. **I.** 3D view of the predicted AtPWO1–AtSWN complex shown from panel G rank 1 with, **J.** Zoom-in view of the interaction interface, highlighting the interaction between the AtPWO1 C-tail (rainbow-colored) and AtSWN (blue surface). **K.** Schematic diagram showing the truncated fragments of SmPWOa and SmCLF used for interaction prediction**. L.** pLDTT graph showing values for five models for SmPWOa (377-619 aa) interaction with SmCLF (166-364 aa). **M.** PEA plot displaying the rank 3 models from panel L with local interactions in PAE maps, outlined by yellow dotted box. **N.** 3D view of the predicted SmPWOa–SmCLF complex shown in panel L rank 3 with, **O.** Zoom-in view of the interaction interface, highlighting the interaction between the SmPWOa C-tail (rainbow-colored) and SmCLF (blue surface).

**Supplementary Figure 14.**
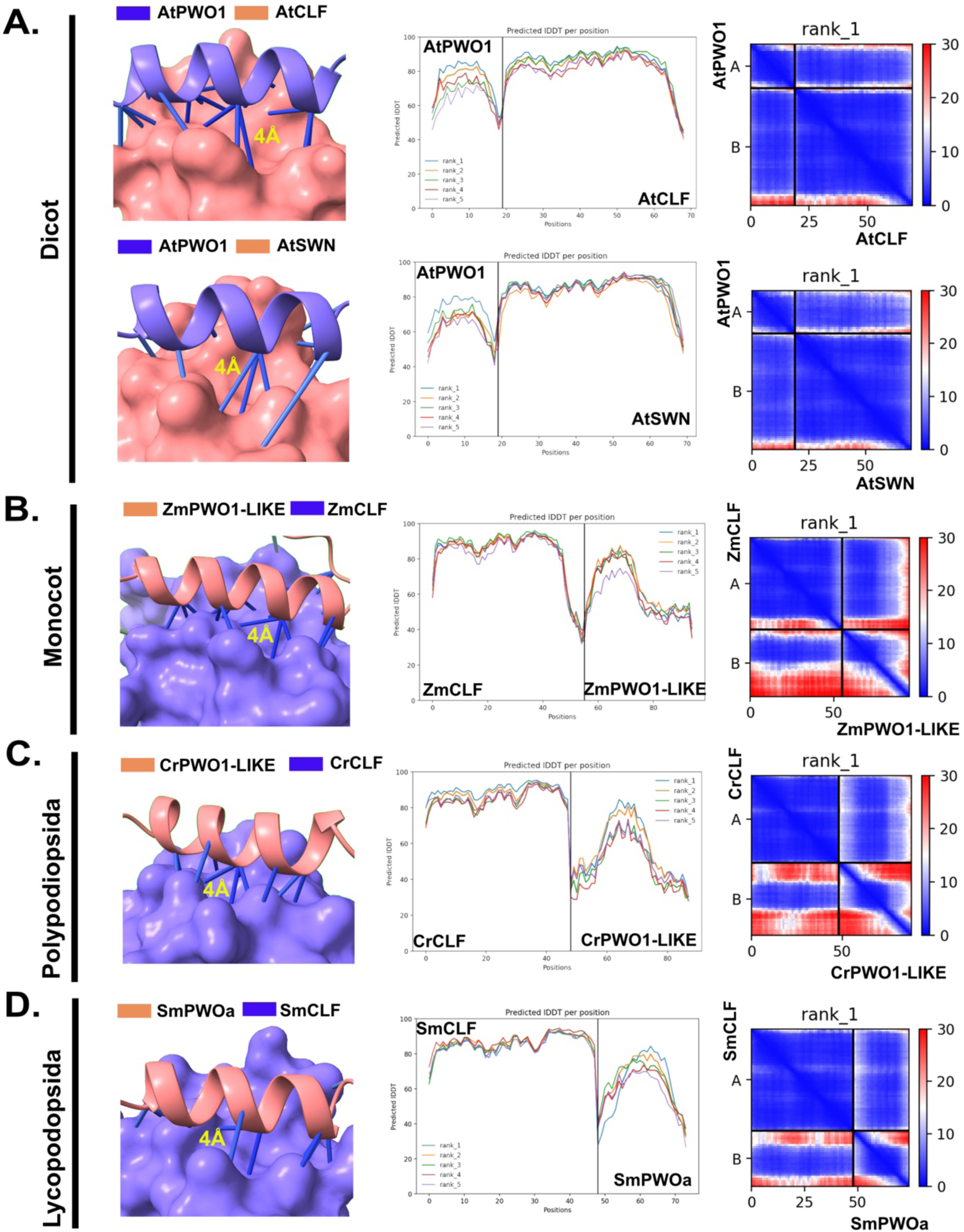
AF2-M prediction of interaction surfaces between PWOs and PRC2 catalytic subunit (CLF) across plant species. Predicted interaction surfaces between PWOs and the PRC2 catalytic subunit (CLF) in selected representative plant species. AF2-M-based predictions of specific interaction interfaces are shown for: **A.** AtPWO1 (735–754 aa) with AtCLF (198–248 aa) and AtPWO1 (735–754 aa) with AtSWN (185–236 aa). **B.** ZmPWO1 (777–817 aa) with ZmCLF (223–281 aa). **C.** CrPWO1 (756–796 aa) with CrCLF (221–269 aa). **D.** SmPWOa (593–691 aa) with SmCLF (182–230 aa). Each panel includes a pLDDT graph showing the confidence values across five models and a PAE plot displaying the top-ranked model for each interaction.

**Supplementary Figure 15.**
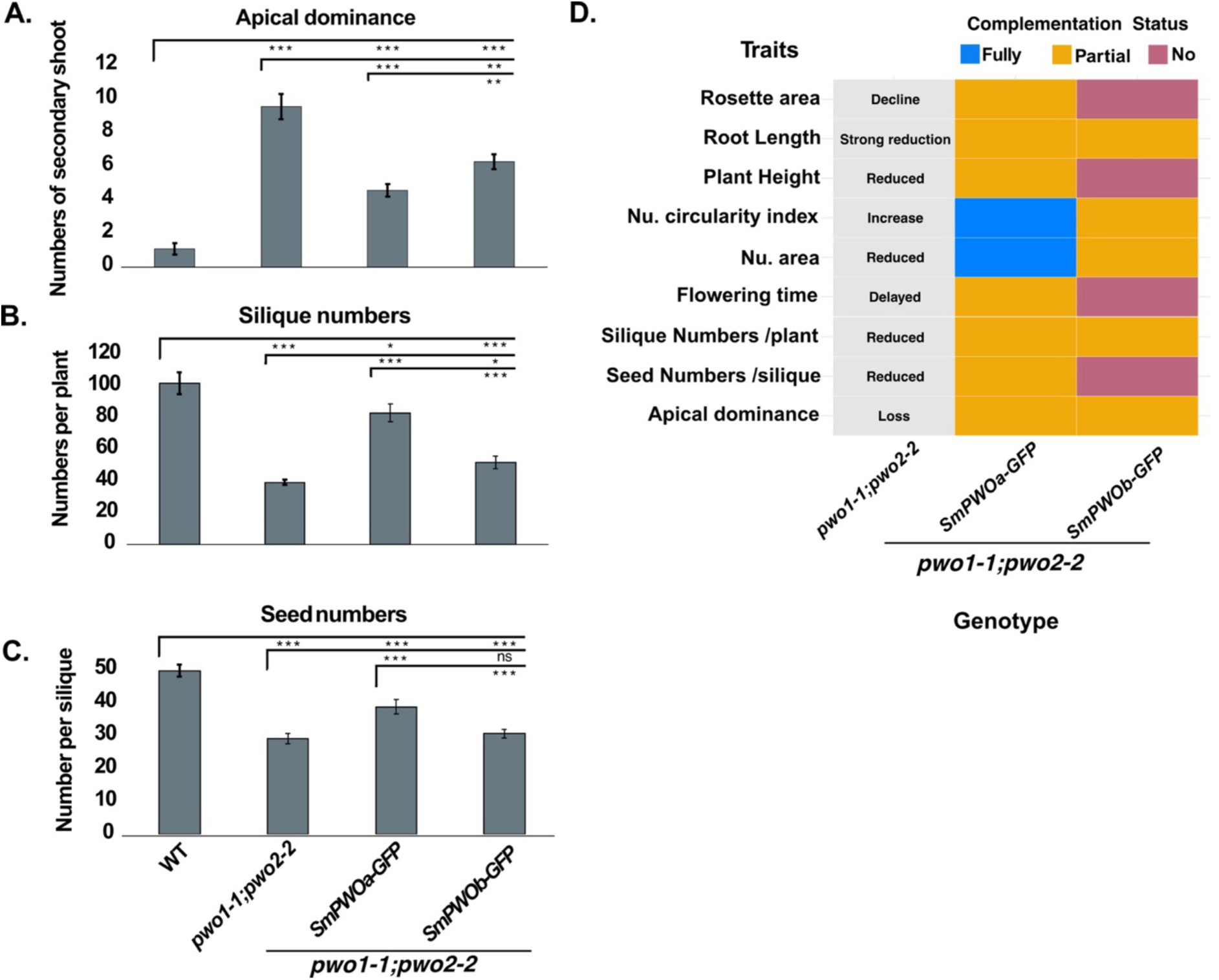
Role of PWOs in maintaining apical dominance, fertilization, and seed development in Arabidopsis. **A.** Number of secondary shoots in 55-day-old seedlings (n=10); **B.** Number of siliques per plant (n=10); **C.** Seed number per silique (n=10) for the genotypes Col-0, *pwo1-1;pwo2-2*, *2×35S_pro_::SmPWOa-GFP*/*pwo1-1; pwo2-2*, and *2×35S_pro_::SmPWOb-GFP*/*pwo1-1;pwo2-2* lines. **D.** Summary of all developmental defects observed in *pwo1-1;pwo2-2* mutants and their complementation status (partial, fully, or none) by the *SmPWOa-GFP* and *SmPWOb-GFP* transgenic lines. Error bars correspond to ±SD. Asterisks represent p-values: ***p ≤ 0.001, **p ≤ 0.01, *p ≤ 0.05. The two-tailed Student’s t-test was applied to calculate the significance level.

**Supplementary Figure 16.**
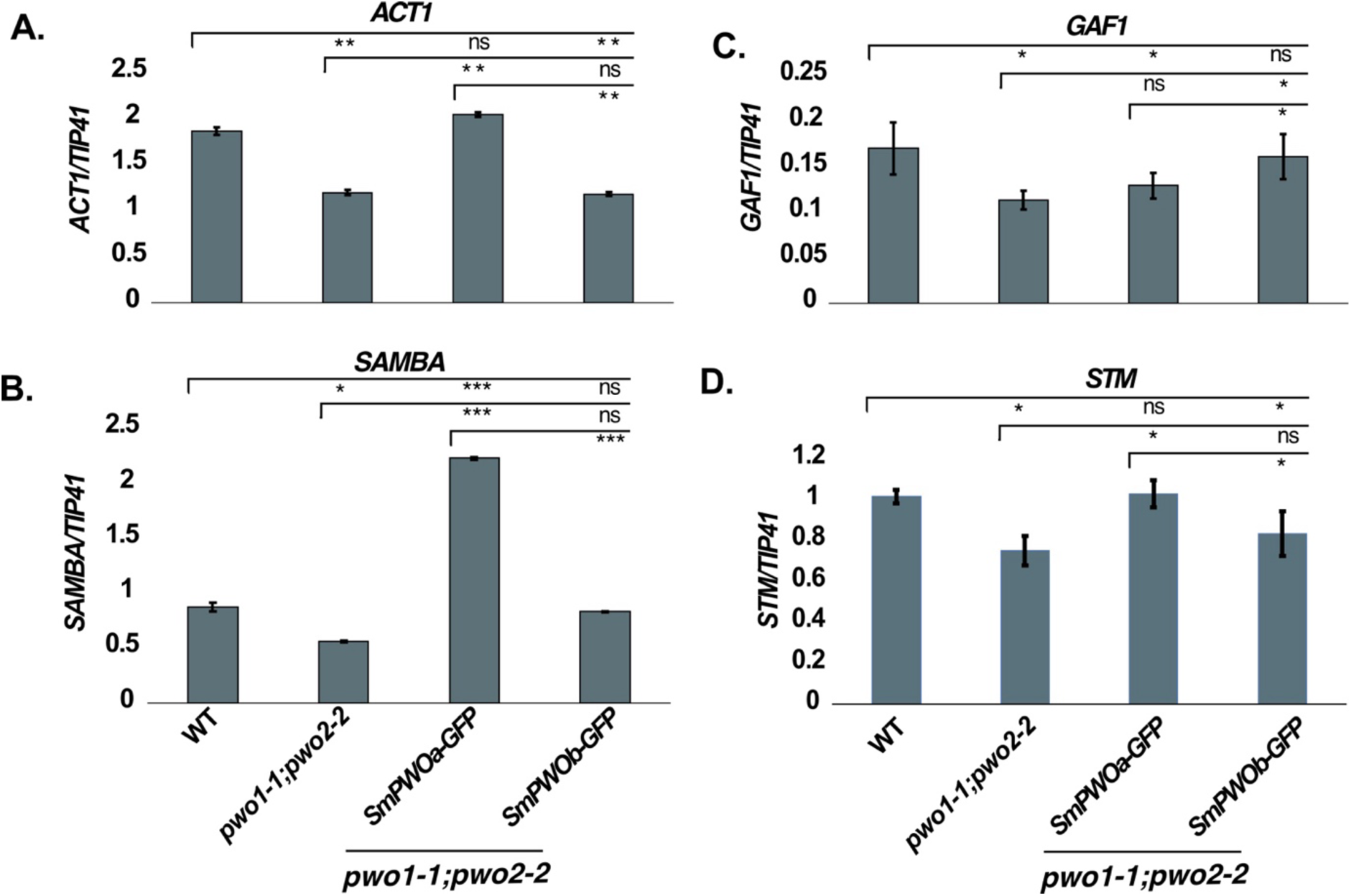
RT-qPCR analyses of *pwo1-1;pwo2-2* misregulated genes. Quantitative RT-qPCR analyses of **A.** *ACTINE 1* (*ACT1*), **B.** *SAMBA*, **C.** *GAMETOPHYTE DEFECTIVE 1* (*GAF1*) and **D.** *SHOOT MERISTEMLESS* (*STM*) expression levels in the genotypes Col-0, *pwo1-1;pwo2-2*, *2×35S_pro_::SmPWOa-GFP*/*pwo1-1;pwo2-2*, and *2×35S_pro_::SmPWOb-GFP*/*pwo1-1;pwo2-2* lines. Results were normalized using *TAP42 INTERACTING PROTEIN OF 41 KDA* (*TIP41*) as housekeeping gene. All samples were from 10-day-old seedlings. Data represent the mean values of three replicates. Two biological repeats were conducted. Error bars correspond to ±SD. Asterisks represent p-values: ***p ≤ 0.001, **p ≤ 0.01, *p ≤ 0.05. The two-tailed Student’s t-test was applied to calculate the significance level.

## Notes

### Competing Interest Statement

The authors have declared no competing interest.

