## Supplementary Figure 1 for "PWO proteins are associated with PRC2 since their emergence in vascular plants"

Supplementary Figure 1. Multiple sequence alignment of conserved PWWP domain across the plant evolution.

```
                                                     1        10        20        30        40        50        60
AT1G51745.1_Arabidopsis_thaliana_Eudicot             INASVGRLVWVRRRNGSWWPGQTLVHDQVPDNSLVGPKVGTPIKLLGRDDVSVDWYILEN
AT3G03140.1_Arabidopsis_thaliana_Eudicot             VDWTVGSIVWVRRRNGSWWPGRILGQEDLDSTHITSPRSGTPVKLLGREDASVDWYNLEK
AT3G21295.1_Arabidopsis_thaliana_Eudicot             IDASVGGLVWVRRRNGAWWPGRIMAHHEVPDGTIVSPKSGTPIKLLGRDDASVDWYNLEK
AT3G21295.2_Arabidopsis_thaliana_Eudicot             IDASVGGLVWVRRRNGAWWPGRIMAHHEVPDGTIVSPKSGTPIKLLGRDDASVDWYNLEK
AT3G21295.3_Arabidopsis_thaliana_Eudicot             IDASVGGLVWVRRRNGAWWPGRIMAHHEVPDGTIVSPKSGTPIKLLGRDDASVDWYNLEK
KAH7296303.1_Ceratopteris_richardii_ferns_Horseta    IDSSVGALVWVRRRNGSWWPGRILGPEELPGSHLLSPRTGTPVKLLGREDGSVDWYNIEK
KAH7296310.1_Ceratopteris_richardii_ferns_Horseta    IDSSVGALVWVRRRNGSWWPGRILGPEELPGSHLLSPRTGTPVKLLGREDGSVDWYNIEK
KAH7435469.1_Ceratopteris_richardii_ferns_Horseta    AESYVGTLVWVRRRNGHWWPGRVLAAHEVPSSHLLPPKSGTPVKLLGKDDASVDWYNIGK
KAH7444823.1_Ceratopteris_richardii_ferns_Horseta    IDNSVGALVWVRRRNGSWWPGRILGPEELPGSHLLSPRTGTPVKLLGREDGSVDWYNIEK
KAH7444826.1_Ceratopteris_richardii_ferns_Horseta    IDNSVGALVWVRRRNGSWWPGRILGPEELPGSHLLSPRTGTPVKLLGREDGSVDWYNIEK
XP_024515257.1_Selaginella_moellendorffii_spike_m    VDTAVGTIVWVRRRNGSWWPGRIISDSELSTANLLSPRSGTPVKLLGREDGSVDWYNIEK
XP_024515258.1_Selaginella_moellendorffii_spike_m    VDTAVGTIVWVRRRNGSWWPGRIISDSELSTANLLSPRSGTPVKLLGREDGSVDWYNIEK
XP_024535029.1_Selaginella_moellendorffii_spike_m    VDTAVGTIVWVRRRNGSWWPGRIISDSELSTANLLSPRSGTPVKLLGREDGSVDWYNIEK
XP_024535030.1_Selaginella_moellendorffii_spike_m    VDTAVGTIVWVRRRNGSWWPGRIISDSELSTANLLSPRSGTPVKLLGREDGSVDWYNIEK
XP_024540635.1_Selaginella_moellendorffii_spike_m    VKPVPGTIVWVRRRNGSWWPGRIISKKELLASNVMSPRSGTPVKLLGRDDASVDWYNIEK
XP_024541679.1_Selaginella_moellendorffii_spike_m    VKPVPGTIVWVRRRNGSWWPGRIISKKELLASNVMSPRSGTPVKLLGRDDASVDWYNIEK
XP_021890214.1_Carica_papaya_Eudicot                 IDASVGGLVWVRRRNGSWWPGRIMGLNEVSESCLVSPRSGTPVKLLGRDDASVDWYNLEK
XP_021890215.1_Carica_papaya_Eudicot                 IDASVGGLVWVRRRNGSWWPGRIMGLNEVSESCLVSPRSGTPVKLLGRDDASVDWYNLEK
XP_021890216.1_Carica_papaya_Eudicot                 IDASVGGLVWVRRRNGSWWPGRIMGLNEVSESCLVSPRSGTPVKLLGRDDASVDWYNLEK
XP_021890217.1_Carica_papaya_Eudicot                 IDASVGGLVWVRRRNGSWWPGRIMGLNEVSESCLVSPRSGTPVKLLGRDDASVDWYNLEK
XP_021900190.1_Carica_papaya_Eudicot                 VDCGVGSIVWVRRRNGSWWPGKILGPEELAASHLTSPRSGTPVKLLGREDASVDWYNLEK
XP_021900191.1_Carica_papaya_Eudicot                 VDCGVGSIVWVRRRNGSWWPGKILGPEELAASHLTSPRSGTPVKLLGREDASVDWYNLEK
XP_004297615.1_Fragaria_vesca_Eudicot                IDASVGSLVWVRRRNGSWWPGRIVGLDELSEDFVVSPRSGTPVKLLGRDDASVDWYNLEK
XP_004307887.1_Fragaria_vesca_Eudicot                GDFSVGSIVWVRRRNGSWWPGKIVGPEELSTSHLTSPRSGTPVKLLGREDASVDWYNLEK
XP_011463342.1_Fragaria_vesca_Eudicot                IDASVGSLVWVRRRNGSWWPGRIVGLDELSEDFVVSPRSGTPVKLLGRDDASVDWYNLEK
XP_031483191.1_Nymphaea_colorata_Eudicot             MDCSVGTIVWVRRRNGSWWPGRILGQDELSETHLMSPRSGTPVKLLGREDASVDWYNLEK
XP_031502312.1_Nymphaea_colorata_Eudicot             VDCGVGTIVWVRRRNGSWWPGRILGQDELSATHLMSPRSGTPVKLLGREDASVDWYNLEK
XP_043631659.1_Erigeron_canadensis_Eudicot           VDCGVGTIVWVRRRNGSWWPGKILGPQHLSSSHLMSPRSGTPVKLLGREDASVDWYNLEK
XP_043637694.1_Erigeron_canadensis_Eudicot           LDPKIGGLVWVRRRNGSWWPGRILGPEELPESCLIPIRSGTPVKLLGREDASVDWYNLER
XP_042949184.1_Carya_illinoinensis_Eudicot           IDASVGGLVWVRRRNGSWWPGRIMGLDELSEGSLVSPRSGTPVKLLGREDASVDWYNLEK
XP_042949194.1_Carya_illinoinensis_Eudicot           IDASVGGLVWVRRRNGSWWPGRIMGLDELSEGSLVSPRSGTPVKLLGREDASVDWYNLEK
XP_042949203.1_Carya_illinoinensis_Eudicot           IDASVGGLVWVRRRNGSWWPGRIMGLDELSEGSLVSPRSGTPVKLLGREDASVDWYNLEK
XP_042949210.1_Carya_illinoinensis_Eudicot           IDASVGGLVWVRRRNGSWWPGRIMGLDELSEGSLVSPRSGTPVKLLGREDASVDWYNLEK
XP_042949220.1_Carya_illinoinensis_Eudicot           IDASVGGLVWVRRRNGSWWPGRIMGLDELSEGSLVSPRSGTPVKLLGREDASVDWYNLEK
XP_042967952.1_Carya_illinoinensis_Eudicot           IDASVGGLVWVRRRNGSWWPGRIMGLDELSEGSLVSPRSGTPVKLLGREDASVDWYNLEK
XP_042979081.1_Carya_illinoinensis_Eudicot           SACIPGSIVWVRRRNGSWWPGKILGPEELASSHLTSPRSGTPVKLLGREDASVDWYNLEK
XP_042983017.1_Carya_illinoinensis_Eudicot           AGCSPGSIVWVRRRNGSWWPGKILGPEELGASHLTSPRSGTPVKLLGREDASVDWYNLEK
XP_042983018.1_Carya_illinoinensis_Eudicot           AGCSPGSIVWVRRRNGSWWPGKILGPEELGASHLTSPRSGTPVKLLGREDASVDWYNLEK
XP_042983019.1_Carya_illinoinensis_Eudicot           AGCSPGSIVWVRRRNGSWWPGKILGPEELGASHLTSPRSGTPVKLLGREDASVDWYNLEK
XP_042983020.1_Carya_illinoinensis_Eudicot           AGCSPGSIVWVRRRNGSWWPGKILGPEELGASHLTSPRSGTPVKLLGREDASVDWYNLEK
XP_042983022.1_Carya_illinoinensis_Eudicot           AGCSPGSIVWVRRRNGSWWPGKILGPEELGASHLTSPRSGTPVKLLGREDASVDWYNLEK
XP_042983023.1_Carya_illinoinensis_Eudicot           AGCSPGSIVWVRRRNGSWWPGKILGPEELGASHLTSPRSGTPVKLLGREDASVDWYNLEK
Thupl.29378772s0023.1.p_Thuja_plicata_Pinopsida     IDTVVGTIVWVRRRNGSWWPGRILGADELSETHLLSPRSGTPVKLLGREDASVDWYNIEK
Thupl.29378772s0023.7.p_Thuja_plicata_Pinopsida     IDTVVGTIVWVRRRNGSWWPGRILGADELSETHLLSPRSGTPVKLLGREDASVDWYNIEK
Thupl.29378772s0023.3.p_Thuja_plicata_Pinopsida     IDTVVGTIVWVRRRNGSWWPGRILGADELSETHLLSPRSGTPVKLLGREDASVDWYNIEK
Thupl.29378772s0023.6.p_Thuja_plicata_Pinopsida     IDTVVGTIVWVRRRNGSWWPGRILGADELSETHLLSPRSGTPVKLLGREDASVDWYNIEK
Thupl.29378772s0023.4.p_Thuja_plicata_Pinopsida     IDTVVGTIVWVRRRNGSWWPGRILGADELSETHLLSPRSGTPVKLLGREDASVDWYNIEK
Thupl.29378772s0023.5.p_Thuja_plicata_Pinopsida     IDTVVGTIVWVRRRNGSWWPGRILGADELSETHLLSPRSGTPVKLLGREDASVDWYNIEK
Thupl.29378772s0023.2.p_Thuja_plicata_Pinopsida     IDTVVGTIVWVRRRNGSWWPGRILGADELSETHLLSPRSGTPVKLLGREDASVDWYNIEK
Thupl.29378072s0023.1.p_Thuja_plicata_Pinopsida     VDDSIGSIVWVRRRNGSWWPGRILAANELSLSHLMSPRSGTPVKLLGREDASVDWYNLEK
scaffold_ABIJ_2009220_Selaginella_lepidophylla_sp    VDTAVGTIVWVRRRNGSWWPGRIISDAELSTANLLSPRSGTPVKLLGREDGSVDWYNIEK
scaffold_CQPW_2008166_Anemia_tomentosa_ferns_Hors    VDSSVGTVVWVRRRNGRWWPGRVISELEVRRSRLLTNRSGTPVKLLGKEDASVDWYNFGK
scaffold_CQPW_2008167_Anemia_tomentosa_ferns_Hors    VDSSVGTVVWVRRRNGRWWPGRVISELEVRRSRLLTNRSGTPVKLLGKEDASVDWYNFGK
scaffold_CQPW_2008862_Anemia_tomentosa_ferns_Hors    IDDAVGTVVWVRRRNGSWWPGRVISHEEVTRSCLLTKRSGTPVKLLGKDDASVDWYNFGM
scaffold_CQPW_2020670_Anemia_tomentosa_ferns_Hors    IDSSVGALVWVRRRNGSWWPGRILAPEELPGSHLLSPRTGTPVKLLGREDGSVDWYNIEK
scaffold_DFHO_2051597_Danaea_nodosa_ferns_Horseta    VDQSVGTLVWVRRRNGSWWPGRILGAEELPGSHLLSPRSGTPVKLLGREDASVDWYNIEK
scaffold_EEAQ_2011348_Sceptridium_dissectum_ferns    .EASVGTLVWVRRRNGSWWPGRILGPEELPGSHLLSPRSGTPVKLLGREDASVDWYNIEK
RWR72543.1_Cinnamomum_micranthum_Magnoliopsida       IDATVGGLVWVHRPNGSWWPGQIIGLDELPAACLVSQKSGTPVKLLGREDANIDWYNLEK
RWR81067.1_Cinnamomum_micranthum_Magnoliopsida       IDCSVGTIVWVRRRNGSWWPGRILGSDELSASHLMSPRSGTPVKLLGREDASVDWYNLEK
RWR86839.1_Cinnamomum_micranthum_Magnoliopsida       VDCSVGTIVWVRRRNGSWWPGRILGPNELSASHLMSPRSGTPVKLLGREDASVDWYNLEK
RWR87272.1_Cinnamomum_micranthum_Magnoliopsida       IDCSVGTIVWVRRRNGSWWPGRILGPGELSASHLMSPRSGTPVKLLGREDASVDWYNLEK
KAG1362729.1_Cocos_nucifera_Monocot                  ADCSVGTIVWVRRRNGSWWPGRILGPDELSASHLMSPRSGTPVKLLGREDASVDWYNLEK
KAG1362730.1_Cocos_nucifera_Monocot                  ADCSVGTIVWVRRRNGSWWPGRILGPDELSASHLMSPRSGTPVKLLGREDASVDWYNLEK
KAF3321551.1_Carex_littledalei_Monocot               VDASVGTIVWVRRRNGSWWPGRILGPDELAESHLLSPRSGTPVKLLGREDASVDWYNLEK
KAF3324589.1_Carex_littledalei_Monocot               EDRSPGTLVWVRRRNGSWWPGRILSKEEIPTKLRRPQRSGIPIKLLGREDGNIDWYNLDK
KAF8671051.1_Digitaria_exilis_Monocot                ADAEVGALVWVRRRNGSWWPGRILGMDELPENTVIPPRSGTPIKLLGRADGSIDWYNLEK
KAF8696638.1_Digitaria_exilis_Monocot                ADAEVGALVWVRRRNGSWWPGRILGMDELPENTVIPPRSGTPIKLLGRADGSIDWYNLEK
KAF8724822.1_Digitaria_exilis_Monocot                VDVFAGTLVWLRRPNGSWWPSIVISPQDVPGGAVSPPRCATPIMLLGRREGFVDWCNIER
KAF8729756.1_Digitaria_exilis_Monocot                GDTSPGTIVWVRRRNGSWWPGRILGPEELPPSQIMSPRSGTPVKLLGREDASVDWYNLEK
KAF8750219.1_Digitaria_exilis_Monocot                GDTSPGTIVWVRRRNGSWWPGRILGPEELPPSQIMSPRSGTPVKLLGREDASVDWYNLEK
KAG9454374.1_Aristolochia_fimbriata_Magnoliopsida    VDCSAGTIVWVRRRNGSWWPGRILGPEELSASHLMSPRSGTPVKLLGREDASVDWYNLEK
KAG9455570.1_Aristolochia_fimbriata_Magnoliopsida    VDVSAGCLVWVRRRNGSWWPGRVMGLDELPERCSRSPRLGTPVKLLGRKDLNVDWYNFEK
GJM89924.1_Eleusine_coracana_Monocot                 VDVSAGTLVWLRRPNGTWWPSIVISPSDVPDGCPAPPRCAIPIMLLGRRDGFVDWCNIGR
GJM94157.1_Eleusine_coracana_Monocot                 ADAEVGALVWVRRRNGSWWPGRILGMDELPENCVIPPRSGTPIKLLGRPDGSIDWYNLEK
GJM94158.1_Eleusine_coracana_Monocot                 SNAKAGALVWVLRRNGCWWPGQILGTDELPKNYVIPPRSGTPIKLLGRPDGSIDWYNIEK
GJN18634.1_Eleusine_coracana_Monocot                 ADAEVGALVWVRRRNGSWWPGRILGMDELPENCVIPPRSGTPIKLLGRPDGSM.......
URD96891.1_Musa_troglodytarum_Monocot                INVSAGGLVWVRRRNGSWWPGRTVGLHELPVKCLLPPRSGTPIKLLGREDGSMDWYNLEK
URE09140.1_Musa_troglodytarum_Monocot                VDCSVGTIIWVRRRNGSWWPGRILGPEELSVSHLMSPRSGTPVKLLGREDASVDWYNLEK
URE09250.1_Musa_troglodytarum_Monocot                VDCGVGTIVWVRRRNGSWWPGRILGPDELSASHLMSPRSGTPVKLLGREDASVDWYNLEK
URE21899.1_Musa_troglodytarum_Monocot                NNVSAGGLVWVRRPNGSWWPGRVVGRDELPAKCVLPPRSGTPIKLLGREDGSMDWYNLAK
URE21900.1_Musa_troglodytarum_Monocot                NNVSAGGLVWVRRPNGSWWPGRVVGRDELPAKCVLPPRSGTPIKLLGREDGSMDWYNLAK
URE21901.1_Musa_troglodytarum_Monocot                NNVSAGGLVWVRRPNGSWWPGRVVGRDELPAKCVLPPRSGTPIKLLGREDGSMDWYNLAK
URE21903.1_Musa_troglodytarum_Monocot                NNVSAGGLVWVRRPNGSWWPGRVVGRDELPAKCVLPPRSGTPIKLLGREDGSMDWYNLAK
XP_010911968.1_Elaeis_guineensis_Monocot             FDASAGGLVWVRRRNGSWWPGRILGRDELPAKCLLPPRSGTPIKLLGREDGSMDWYNLEK
XP_010911976.1_Elaeis_guineensis_Monocot             FDASAGGLVWVRRRNGSWWPGRILGRDELPAKCLLPPRSGTPIKLLGREDGSMDWYNLEK
XP_010918108.1_Elaeis_guineensis_Monocot             VDCSVGTIVWVRRRNGSWWPGRILGPDELSASHLMSPRSGTPVKLLGREDASVDWYNLEK
XP_010924938.1_Elaeis_guineensis_Monocot             IDVSAGGLVWIRRRNGSWWPGRILGRDELPAKCLLPPRSGTPIKLLGREDGSMDWYNLEK
XP_010924939.1_Elaeis_guineensis_Monocot             IDVSAGGLVWIRRRNGSWWPGRILGRDELPAKCLLPPRSGTPIKLLGREDGSMDWYNLEK
XP_010943463.1_Elaeis_guineensis_Monocot             VDCSVGTIVWVRRRNGSWWPGRILGPDELSASHLMSPRSGTPVKLLGREDASVDWYNLEK
XP_019707050.1_Elaeis_guineensis_Monocot             IDVSAGGLVWIRRRNGSWWPGRILGRDELPAKCLLPPRSGTPIKLLGREDGSMDWYNLEK
XP_042377257.1_Zingiber_officinale_Monocot           SNVSAGGLVWVRRPNGSWWPGRVVGLDELEAKCVVPPRSGTPIKLLGREDGSMDWYNLAK
XP_042383299.1_Zingiber_officinale_Monocot           SNVSAGGLVWVRRPNGSWWPGRVVGLDELEAKCVVPPRSGTPIKLLGREDGSMDWYNLAK
XP_042383893.1_Zingiber_officinale_Monocot           VDCRVGTIVWVRRRNGSWWPGRILGQEELSASHLMSPRSGTPVKLLGREDASVDWYNLEK
XP_042383902.1_Zingiber_officinale_Monocot           VDCRVGTIVWVRRRNGSWWPGRILGQEELSASHLMSPRSGTPVKLLGREDASVDWYNLEK
XP_042387214.1_Zingiber_officinale_Monocot           VDCCVGTIVWVRRRNGSWWPGRILGPEELSVSHLMSPRSGTPVKLLGREDASVDWYNLEK
XP_042387215.1_Zingiber_officinale_Monocot           VDCCVGTIVWVRRRNGSWWPGRILGPEELSVSHLMSPRSGTPVKLLGREDASVDWYNLEK
XP_042392597.1_Zingiber_officinale_Monocot           VDCCVGTIVWVRRRNGSWWPGRILGPEELSVSHLMSPRSGTPVKLLGREDASVDWYNLEK
XP_042392598.1_Zingiber_officinale_Monocot           VDCCVGTIVWVRRRNGSWWPGRILGPEELSVSHLMSPRSGTPVKLLGREDASVDWYNLEK
XP_042424141.1_Zingiber_officinale_Monocot           VDCGVGTIVWVRRRNGSWWPGRILGQEELSASHLMSPRSGTPVKLLGREDASVDWYNLEK
XP_042424142.1_Zingiber_officinale_Monocot           VDCGVGTIVWVRRRNGSWWPGRILGQEELSASHLMSPRSGTPVKLLGREDASVDWYNLEK
XP_042424143.1_Zingiber_officinale_Monocot           VDCGVGTIVWVRRRNGSWWPGRILGQEELSASHLMSPRSGTPVKLLGREDASVDWYNLEK
XP_042427101.1_Zingiber_officinale_Monocot           VDCSVGTIVWVRRRNGSWWPGRILGQEELSASHLMSPRSGTPVKLLGREDASVDWYNLEK
XP_042427102.1_Zingiber_officinale_Monocot           VDCSVGTIVWVRRRNGSWWPGRILGQEELSASHLMSPRSGTPVKLLGREDASVDWYNLEK
```

```
XP_042427103.1_Zingiber_officinale_Monocot              VDCSVGTIVWVRRRNGSWWPGRILGQEELSASHLMSPRSGTPVKLLGREDASVDWYNLEK
XP_042450147.1_Zingiber_officinale_Monocot              VDCRVGTIVWVRRRNGSWWPGRILGQEELSASHLMSPRSGTPVKLLGREDASVDWYNLEK
XP_042455834.1_Zingiber_officinale_Monocot              VHYSVGTIVWVRRRNGSWWPGRILGPEELSVSHLMSPRSGTPVKLLGREDASVDWYNLEK
XP_042455835.1_Zingiber_officinale_Monocot              VHYSVGTIVWVRRRNGSWWPGRILGPEELSVSHLMSPRSGTPVKLLGREDASVDWYNLEK
XP_042461069.1_Zingiber_officinale_Monocot              VHYSVGTIVWVRRRNGSWWPGRILGPEELSVSHLMSPRSGTPVKLLGREDASVDWYNLEK
XP_042461070.1_Zingiber_officinale_Monocot              VHYSVGTIVWVRRRNGSWWPGRILGPEELSVSHLMSPRSGTPVKLLGREDASVDWYNLEK
XP_047063722.1_Lolium_rigidum_Monocot                   EDCSPGTIVWVRRRNGSWWPGRILGQDELPASQVVSPRTGTPVKLLGREDASIDWYNLEK
XP_047079877.1_Lolium_rigidum_Monocot                   GDTSPGTIVWVRRRNGSWWPGRILGQDELPPSQIMSPRSGTPVKLLGREDASVDWYNLEK
XP_047092051.1_Lolium_rigidum_Monocot                   ADAEVGALVWVRRRNGSWWPGRILGQDELPENCVVPPRSGTPIKLLGRPDGSIDWYNLEK
XP_044951029.1_Hordeum_vulgare_Monocot                  GDTSPGTIVWVRRRNGSWWPGRILGQDELPPSQIMSPRSGTPVKLLGREDASVDWYNLEK
XP_044963579.1_Hordeum_vulgare_Monocot                  ADVSAGALVWVRRPNGSWWPGRVASRLELPDDCPAPPRSATPILLLGRRDGFVEWCNLER
XP_044967807.1_Hordeum_vulgare_Monocot                  ADCSPGAIVWVRRRNGSWWPGRILGQDELLASQLVTPRTGTPVKLLGREDASVDWYNLEK
XP_044976129.1_Hordeum_vulgare_Monocot                  ADAEVGALVWVRRRNGSWWPGRILGQDELPENCVVPPRSGTPIKLLGRPDGSIDWYNLEK
scaffold_GKAG_2003823_Huperzia_lucidula_clubmoss        ....VGTLVWIRRRNGSWWPGRIISTDELSSLHLTSQRSGTPVKLLGREDASVDWYNVEK
scaffold_KAWQ_2005490_Stangeria_eriopus_Cycadopsi       VDCSVGTIVWVRRRNGSWWPGKILGHNELSVSHLMSPRSGTPVKLLGREDASVDWYNMEK
scaffold_KIIX_2013117_Pilularia_globulifera_ferns       IDSSVGALVWVRRRNGSWWPGRILAPEELPGSHLLSPRTGTPVKLLGREDGSVDWYNIEK
scaffold_KIIX_2013283_Pilularia_globulifera_ferns       IDSSVGALVWVRRRNGSWWPGRILGAEELPGSHLLSPRTGTPVKLLGREDGSVDWYNIEK
scaffold_PKOX_2013427_Isoetes_tegetiformans_quill       .....GTVVWVRRRNGSWWPGRIICKTELGASHLLSPRSGTPVKLLGREDASVDWYNIEK
scaffold_PKOX_2101345_Isoetes_tegetiformans_quill       VDTTVGTIVWVRRRNGSWWPGRIISEDELSTANLLSPRSGTPVKLLGREDGSVDWYNIEK
scaffold_PNZO_2012682_Culcita_macrocarpa_ferns_Ho       IDSSVGALVWVRRRNGSWWPGRILGPEELPGSHLLSPRTGTPVKLLGREDGSVDWYNIEK
scaffold_TOXE_2013932_Welwitschia_mirabilis_Gneto       GEYGIGTIVWVRRRNGSWWPGKVVGLNELSANHLMSPRSGTPVKLLGREDASVDWYNMEK
scaffold_UWOD_2005868_Plagiogyria_japonica_ferns_       IDSSVGALVWVRRRNGSWWPGRILGPEELPGSHLLSPRTGTPVKLLGREDGSVDWYNIEK
scaffold_UWOD_2005869_Plagiogyria_japonica_ferns_       IDSSVGALVWVRRRNGSWWPGRILGPEELPGSHLLSPRTGTPVKLLGREDGSVDWYNIEK
scaffold_UWOD_2014926_Plagiogyria_japonica_ferns_       VENSVGTLVWVRRRNGSWWPGRVLAAHELPRSHFLCPKSGTPVKLLGKEDASVDWYNIGK
scaffold_VIBO_2011581_Plenasium_javanicuma_ferns_       VDSSVGTLVWVRRRNGSWWPGRILGPDELPGSHLLSPRSGTPVKLLGREDASVDWYNIEK
scaffold_VSRH_2006872_Abies_lasiocarpa_Pinopsida        GDYSIGTIVWVRRRNGSWWPGRILGMNELSVSHLMSPRSGTPVKLLGREDASVDWYNMEK
scaffold_VSRH_2012377_Abies_lasiocarpa_Pinopsida        IDTAVGTIVWVRRRNGSWWPGRILGVEELSATHLMSPRSGTPVKLLGRDDASVDWYNIEK
scaffold_XNXF_2063615_Dendrolycopodium_obscurum_c       VKPVVGTLVWIRRRNGSWWPGRIISTDELSSSHIMSQRSGTPVKLLGREDASVDWYNVEK
scaffold_YFZK_2002000_Sciadopitys_verticillata_Pi       ...STGSIVWVRRRNGSWWPGRILGLNELSRSHLMSPRH.......GREDASVDWYNLEK
scaffold_YFZK_2004458_Sciadopitys_verticillata_Pi       IDTAVGTIVWVRRRNGSWWPGKILGPDELSETHLMSPRSGTP...LGREDASVDWYNIEK
scaffold_ZYCD_2042766_Selaginella_acanthonota_spi       VDAAVGTIVWVRRRNGSWWPGRIISDAELSTANLLSPRSGTPVKLLGREDGSVDWYNIEK
V7AHI2|V7AHI2_PHAVU_Phaseolus_vulgaris_Eudicot          MDCGVGSIVWVRRRNGSWWPGQILGPDDLSASHLTSPRSGTPVKLLGREDASVDWYNLEK
V7AQI6|V7AQI6_PHAVU_Phaseolus_vulgaris_Eudicot          IDASVGGLVWVRRRNGSWWPGRILGLHELSESCLVSPRSGTPVKLLGREDASVDWYNLEK
V7CFU1|V7CFU1_PHAVU_Phaseolus_vulgaris_Eudicot          VDCDVGSIVWVRRRNGSWWPGQILGPDHLSASHLTSPRSGTPVKLLGREDASVDWYNLEK
Q0DYD8|Q0DYD8_ORYSJ_Oryza_sativa_Monocot                GDTSPGTIVWVRRRNGSWWPGRILGPDELPPSQIMSPRSGTPVKLLGREDASVDWYNIEK
Q7X7G8|Q7X7G8_ORYSJ_Oryza_sativa_Monocot                TDCSPGIIVWVRRRNGSWWPGRILGPDELPASQVMSPKTGTPVKLLGREDASVDWYNLEK
Q0JLX9|Q0JLX9_ORYSJ_Oryza_sativa_Monocot                RDAEVGALVWVRRRNGSWWPGQILSADELPENCVVPPRSGTPIKLLGRPDGSIDWYNLEK
C5XN34|C5XN34_SORBI_Sorghum_bicolor_Monocot             PDAEVGALVWVRRRNGSWWPGRILGMDELPENIVIPPRSGTPIKLLGRPDGSIDWYNLEK
A0A1B6PAF8|A0A1B6PAF8_SORBI_Sorghum_bicolor_Monoc       VDVSVGTLVWLRRPNGSWWPSIVISPQDVPEGCPPPPRCATPIMLLGRRDGYVDWCNLDR
C5Y0E5|C5Y0E5_SORBI_Sorghum_bicolor_Monocot             GDTSPGTIVWVRRRNGSWWPGRILGPEELPPSQIMSPRSGTPVKLLGREDASVDWYNLEK
A0A3Q7H2C5|A0A3Q7H2C5_SOLLC_Solanum_lycopersicum_       GDCGVGSIVWVRRRNGSWWPGKILCPDEFSASHLMSPRSGTPVKLLGREDATVDWYNLEK
A0A3Q7FMI9|A0A3Q7FMI9_SOLLC_Solanum_lycopersicum_       LECDTGSIVWVRRRNGSWWPGKILGTNELSATHLMSPRSGTPVKLLGREDASVDWYNLEK
A0A3Q7FVX7|A0A3Q7FVX7_SOLLC_Solanum_lycopersicum_       IDASVGGLVWVRRRNGSWWPGRILGSDELPQSCSVSPRLGTPVKLLGREDASVDWYNLEK
B9I743|B9I743_POPTR_Populus_trichocarpa_Eudicot         ADGGIGPIVWVRRRNGSWWPGHIMEADELAEYNLTSPRTGTPVKLLGRDDASVDWYNLEK
A0A2K2C0C5|A0A2K2C0C5_POPTR_Populus_trichocarpa_E       IDASVGALVWVRRRNGSWWPGRIVGLDEISEGSLVSPRSGTPVKLLGREDASVDWYNLEK
A0A2K2B1H6|A0A2K2B1H6_POPTR_Populus_trichocarpa_E       IDASVGALVWVRRRNGSWWPGRIVGLDEISEGSLVSPRSGTPVKLLGREDASVDWYNLEK
A0A2K1WPP7|A0A2K1WPP7_POPTR_Populus_trichocarpa_E       ADGNIGPIVWVRRRNGSWWPGQIMEADELAEYNLTSPRTGTPVKLLGRDDASVDWYNLEK
A0A804QV00|A0A804QV00_MAIZE_Zea_mays_Monocot            PDAEVGALVWVRRRNGSWWPGRILGMGELPENIVIPPRSGTPIKLLGRPDGSIDWYNLEK
A0A804QP35|A0A804QP35_MAIZE_Zea_mays_Monocot            VDVSVGTLVWLRRPNGSWWPSIVISPQDVPEGCAPPPRCATPIMLLGRRDGYVDWCNLDR
A0A804QS22|A0A804QS22_MAIZE_Zea_mays_Monocot            PDAEVGALVWVRRRNGSWWPGRILGMGELPENIVIPPRSGTPIKLLGRPDGSIDWYNLEK
A0A804QSQ4|A0A804QSQ4_MAIZE_Zea_mays_Monocot            PDAEVGALVWVRRRNGSWWPGRILGMGELPENIVIPPRSGTPIKLLGRPDGSIDWYNLEK
A0A804QQ17|A0A804QQ17_MAIZE_Zea_mays_Monocot            PDAEVGALVWVRRRNGSWWPGRILGMGELPENIVIPPRSGTPIKLLGRPDGSIDWYNLEK
A0A804NUY1|A0A804NUY1_MAIZE_Zea_mays_Monocot            GDTSPGTIVWVRRRNGSWWPGRILGPQELPPSQIMSPRSGTPVKLLGREDASVDWYNLEK
A0A804NI78|A0A804NI78_MAIZE_Zea_mays_Monocot            PDAEVGALVWVRRRNGSWWPGRILGMDELPENTVIPPRSGTPIKLLGRSDGSIDWYNLEK
A0A1D6MDX6|A0A1D6MDX6_MAIZE_Zea_mays_Monocot            VDVSVGTLVWLRRPNGSWWPSIVISPQDVPEGCAPPPRCATPIMLLGRRDGFVDWCNLAR
K7UY21|K7UY21_MAIZE_Zea_mays_Monocot                    VDVSVGTLVWLRRPNGSWWPSIVISPQDVPEGCAPPPRCATPIMLLGRRDGYVDWCNLDR
A0A1D6NMK7|A0A1D6NMK7_MAIZE_Zea_mays_Monocot            PDAEVGALVWVRRRNGSWWPGRILGMDELPENTVIPPRSGTPIKLLGRSDGSIDWYNLEK
A0A1D6HH07|A0A1D6HH07_MAIZE_Zea_mays_Monocot            GDTSPGTIVWVRRRNGSWWPGRILGPEELPPSQIMSPRSGTPVKLLGREDASVDWYNLEK
A0A1D6HH08|A0A1D6HH08_MAIZE_Zea_mays_Monocot            GDTSPGTIVWVRRRNGSWWPGRILGPEELPPSQIMSPRSGTPVKLLGREDASVDWYNLEK
A0A1D6DQP9|A0A1D6DQP9_MAIZE_Zea_mays_Monocot            GDTSPGTIVWVRRRNGSWWPGRILGPQELPPSQIMSPRSGTPVKLLGREDASVDWYNLEK
A0A2K2CIB3|A0A2K2CIB3_BRADI_Brachypodium_distachy       SDCSPGTIVWVRRRNGSWWPGRILGPDELRASQVVSPKTGTPVKLLGRDDASVDWYNLEK
I1HNL6|I1HNL6_BRADI_Brachypodium_distachyon_Monoc       ADAEVGALVWVRRRNGSWWPGRILGQDELPENCVVPPRSGTPIKLLGRPDGSIDWYNLEK
I1HFR1|I1HFR1_BRADI_Brachypodium_distachyon_Monoc       VDVSAGTLVWVRRPNGSWWPGQVLSRADVPDGCPAPPRSATPIMLLGRRDGFVDWCNLER
I1ICW8|I1ICW8_BRADI_Brachypodium_distachyon_Monoc       GDTSPGTIVWVRRRNGSWWPGRILGQDELPPSQIMSPRSGTPVKLLGREDASVDWYNLEK
I1LIJ1|I1LIJ1_SOYBN_Glycine_max_Eudicot                 MDCGVGSIVWVRRRNGSWWPGQILGHDDLSAAHLTSPRSGTPVKLLGREDASVDWYNLEK
K7LMH1|K7LMH1_SOYBN_Glycine_max_Eudicot                 VDCDVGSIVWVRRRNGSWWPGQILGPDHLSASHLTSPRSGTPVKLLGREDASVDWYNLEK
K7LSJ6|K7LSJ6_SOYBN_Glycine_max_Eudicot                 MDCGVGSIVWVRRRNGSWWPGQILGPDDLSASHLTSPRSGTPVKLLGREDASVDWYNLEK
A0A0R0LFH3|A0A0R0LFH3_SOYBN_Glycine_max_Eudicot         VDCDVGSIVWVRRRNGSWWPGQILGPDHLSASHLTSPRSGTPVKLLGREDASVDWYNLEK
A0A0R0IPK3|A0A0R0IPK3_SOYBN_Glycine_max_Eudicot         IDASVGGLVWVRRRNGSWWPGRIMGLHELSESCLVSPRSGTPVKLLGREDASVDWYNLEK
I1JAI6|I1JAI6_SOYBN_Glycine_max_Eudicot                 VDCDVGSIVWVRRRNGSWWPGQILGPDHLSASHLTSPRSGTPVKLLGREDASVDWYNLEK
I1KGF6|I1KGF6_SOYBN_Glycine_max_Eudicot                 IDASVGGLVWVRRRNGSWWPGRIMGLHELSESCLVSPRSGTPVKLLGREDASVDWYNLEK
K7LMH0|K7LMH0_SOYBN_Glycine_max_Eudicot                 VDCDVGSIVWVRRRNGSWWPGQILGPDHLSASHLTSPRSGTPVKLLGREDASVDWYNLEK
tr|F6HEG8|F6HEG8_VITVI_Vitis_vinifera_Eudicot           VDSSVGTIVWVRRRNGSWWPGKILGPDELSASHLMSPRSGTPVKLLGREDASVDWYNLEK
tr|F6HY39|F6HY39_VITVI_Vitis_vinifera_Eudicot           IDPSVGGLVWVRRRNGSWWPGRIMGLDELSESCLVSPRSGTPVKLLGREDASVDWYNLEK
M4ET84|M4ET84_BRARP_Brassica_rapa_Eudicot               AGCTVGSIVWVRRRNGSWWPGKILGQDDLDSTHITSPRSGTPVKLLGREDASVDWYNLEK
M4ENQ0|M4ENQ0_BRARP_Brassica_rapa_Eudicot               IKASVGRLVWVRLRNASWWPAQTLLLEQLPQTS.SSPQLGTPIKLLGRDDVNVEWYVLEK
M4DXX3|M4DXX3_BRARP_Brassica_rapa_Eudicot               VDWKVGLIVWVRRRNGSWWPGRILGQDDLDSNTITSPRSGTPVKLLGREDASVDWYNLEK
M4ER48|M4ER48_BRARP_Brassica_rapa_Eudicot               IDASVGGLVWVRRRNGSWWPGRIMAHHEVPDDTIVSPKSGTPIKLLGRDDAGVDWYNLEK
U5CR03_AMBTC_Amborella_trichopoda_Eudicot               VNSSVGSIVWVRRRNGSWWPGRILGLNELSATHLMSPRSGTPVKLLGREDASVDWYNLEK
U5CQG7_AMBTC_Amborella_trichopoda_Eudicot               IDATVGGLVWVRRRNGSWWPGRIMGLDELSETSLVSPRSGTPVKLLGREDASVDWYNIEK
R0I6R0|R0I6R0_9BRAS_Capsella_rubella_Eudicot            ADWTVGSIVWVRRRNGSWWPGKILGQEDLDSTHITSPRSGTPVKLLGREDASVDWYNLEK
R0GPB7|R0GPB7_9BRAS_Capsella_rubella_Eudicot            INASVGRLVWVRRRNGSWWPGQTLVLDEIPDNSLVSPKVGTPIKLLGRDDVSVDWYILEN
R0I129|R0I129_9BRAS_Capsella_rubella_Eudicot            IDASVGGLVWVRRRNGSWWPGRIMAHHEVPDSTIVSPKSGTPIKLLGREDASVDWYNLEK
tr|V4SLI4|V4SLI4_CITCL_Citrus_sinensis_Eudicot          IDPAVGGLVWVRRRNGTWWPGRILGLDEISEGCLVSPRSGTPVKLLGREDASVDWYNLEK
tr|V4U2M5|V4U2M5_CITCL_Citrus_sinensis_Eudicot          GECGVGSIVWVRRRNGSWWPGKILEPEELTTSHLTSPRTGTPVKLLGRDDASVDWYNLEK
tr|A0A058ZXU8|A0A058ZXU8_EUCGR_Eucalyptus_grandis      VGLGVGSIVWVRRRNGSWWPGKILGPDELSTAHLTSPRAGTPVKLLGREDASVDWYNLEK
tr|A0A059BTW7|A0A059BTW7_EUCGR_Eucalyptus_grandis      IDGSVGGLVWVRRRNGSWWPGRIMGHDELPESCLVSPRSGTPVKLLGRKDASIDWYNLEK
A0A1S2YUZ3|A0A1S2YUZ3_CICAR_Cicer_arietinum_Eudic       INASVGGLVWVRRRNGSWWPGRIMSLDELSEGCLVSPRSGTPVKLLGREDASVDWYNLEK
A0A1S3EG88|A0A1S3EG88_CICAR_Cicer_arietinum_Eudic       VDCAVGSIVWVRRRNGSWWPGQILCSDDLSASHLTSPRSGTPVKLLGREDASVDWYNLEK
A0A1S2YJA8|A0A1S2YJA8_CICAR_Cicer_arietinum_Eudic       MDFGVGSIVWVRRRNGSWWPGQIIDPDDIAASHLTSPRSGTPVKLLGREDASVDWYNLEK
A0A251JRZ2|A0A251JRZ2_MANES_Manihot_esculenta_Eud       IDASVGGLVWVRRRNGSWWPGRIMGLDEISEGSLVSPRSGTPVKLLGREDASVDWYNLEK
A0A2C9VI54|A0A2C9VI54_MANES_Manihot_esculenta_Eud       ADCGIGPIVWVRRRNGSWWPGKILGPEELAECNLTSPRTGTPVKLLGREDASVDWYNLEK
A0A2C9VA71|A0A2C9VA71_MANES_Manihot_esculenta_Eud       VDCGVGPIVWVRRRNGSWWPGKIMSPEELAECNLSSPRTGTPVKLLGREDASVDWYNLEK
A0A2C9VA68|A0A2C9VA68_MANES_Manihot_esculenta_Eud       VDCGVGPIVWVRRRNGSWWPGKIMSPEELAECNLSSPRTGTPVKLLGREDASVDWYNLEK
A0A1Q3CJL5_CEPFO_Cephalotus_follicularis_Eudicot        IDSSVGGLVWVRRRNGSWWPGRIMSLDEVTEGCVISPRSGTPVKLLGREDASVDWYNLEK
A0A1Q3D3W0_CEPFO_Cephalotus_follicularis_Eudicot        VECEAGSIVWVRRRNGSWWPGKILGPEELPTSHLTSPRSGTPVKLLGREDASVDWYNLEK
A0A1J6IUT4_NICAT_Nicotiana_attenuata_Eudicot            .ECGAGSIVWVRRRNGSWWPGKILGPDELSASHLISPRSGTPVKLLGREDASVDWYNLEK
A0A1J6KD13_NICAT_Nicotiana_attenuata_Eudicot            IDASVGGLVWVRRRNGSWWPGRILGPDELPQSCSVSPRSGTPVKLLGREDASVDWYNLEK
A0A1U8JDV0|A0A1U8JDV0_GOSHI_Gossypium_hirsutum_Eu       IDASVGGLVWVRRRNGSWWPGRIMGLDELSEGCLVSPRSGTPVKLLGREDASVDWYNLEK
A0A1U8MT23|A0A1U8MT23_GOSHI_Gossypium_hirsutum_Eu       IDASVGGLVWVRRRNGSWWPGRIMGLDELSEGCLVSPRSGTPVKLLGREDASVDWYNLEK
A0A1U8KH73|A0A1U8KH73_GOSHI_Gossypium_hirsutum_Eu       IDASVGGLVWVRRRNGSWWPGRIMGLDELSEGYLVSPRSGTPVKLLGREDASVDWYNLEK
A0A1U8K8A4|A0A1U8K8A4_GOSHI_Gossypium_hirsutum_Eu       GAGGGGLIVWVRRRNGSWWPGKILGPEELPASHLASPRTGTPVKLLGREDASVDWYNLEK
A0A1U8NQ29|A0A1U8NQ29_GOSHI_Gossypium_hirsutum_Eu       DGGGVGSIVWVRRRNGSWWPGKILGPEELPTSHLMSPRTGTPVKLLGREDASVDWYNLEK
```

```
A0A251VD55|A0A251VD55_HELAN_Helianthus_annuus_Eud  IDSTVGGLVWVRRRNGSWWPGRILGPDELPESGLVSPRAGTPVKLLGREDASVDWYNLEK
A0A251TN31|A0A251TN31_HELAN_Helianthus_annuus_Eud  VDYCVGTIVWVQMRNGSWWPGKILGSDERSASHLISPKSGTPVKLLGREDASVDWYNLEK
A0A251UUN3|A0A251UUN3_HELAN_Helianthus_annuus_Eud  IDSAVGGLVWIRRRNGSWWPGRILSPNELPESCLLSPRSGVPVQLLGKEDASVEWYNLEK
A0A251SME0|A0A251SME0_HELAN_Helianthus_annuus_Eud  VDSGVGTIVWVRRRNGSWWPGKILGSDELSASHLMSPRSGTPVKLLGREDASVDWYNLEK
A0A8B7CF00|A0A8B7CF00_PHODC_Phoenix_dactylifera_M  IAASAGGLVWVRRRNGSWWPGRILGWDELPAKCLLPPRSGTPIKLLGREDGSMDWYNLEK
A0A8B7BTD1|A0A8B7BTD1_PHODC_Phoenix_dactylifera_M  VDCSVGTIVWVRRRNGSWWPGRILGPDELSASHLMSPRSGTPVKLLGREDASVDWYNLEK
A0A8B7CIS5|A0A8B7CIS5_PHODC_Phoenix_dactylifera_M  VDCSVGTIVWVRRRNGSWWPGRILGPDELSASHLMSPRSGTPVKLLGREDASVDWYNLEK
A0A2P5AGS9_PARAD_Parasponia_andersonii_Eudicot     IDASVGGLVWVRRRNGSWWPGRIMGLDELSESCLVSPRSGTPVKLLGREDASVDWYNLEK
A0A2P5AY67_PARAD_Parasponia_andersonii_Eudicot     VEFSVGSIVWVRRRNGSWWPGKIVGPEELSASHLTSPRSGTPVKLLGREDASVDWYNLEK
A0A2P6SPQ0_ROSCH_Rosa_chinensis_Eudicot            GDFSVGSIVWVRRRNGSWWPGKIVGPEELSTSHLTSPRSGTPVKLLGREDASVDWYNLEK
A0A2P6RXT9_ROSCH_Rosa_chinensis_Eudicot            IDASVGSLVWVRRRNGSWWPGRIVGLDELSEDFVVSPRAGTPVKLLGRDDASVDWYNLEK
A0A103XW46|A0A103XW46_CYNCS_Cynara_cardunculus_Eu  IDPSVGGLVWVRRRNGSWWPGRILGPDELPESSLVSPRSGTPVKLLGREDASKDWYNLEK
A0A103XCS8|A0A103XCS8_CYNCS_Cynara_cardunculus_Eu  VDCGVGTIVWVRRRNGSWWPGKILGPDELSASHLMSPRSGTPVKLLGREDASVDWYNLEK
A0A2T7FAX4|A0A2T7FAX4_9POAL_Panicum_hallii_Monoco  GDTSPGTIVWVRRRNGSWWPGRILGPEELPPSQIMSPRSGTPVKLLGREDASVDWYNLEK
A0A2T7DKV0|A0A2T7DKV0_9POAL_Panicum_hallii_Monoco  ADAEVGALVWVRRRNGSWWPGRILGMDELPENIVIPPRSGTPIKLLGRPDGSIDWYNLEK
A0A2T7E9E4|A0A2T7E9E4_9POAL_Panicum_hallii_Monoco  VDVSAGTLVWLRRPNGSWWPSIVISPQDVPVGCAAPPRCATPIMLLGRRDGFVDWCNLDR
A0A2T7FAX6|A0A2T7FAX6_9POAL_Panicum_hallii_Monoco  GDTSPGTIVWVRRRNGSWWPGRILGPEELPPSQIMSPRSGTPVKLLGREDASVDWYNLEK
A0A5J9SJ79|A0A5J9SJ79_9POAL_Eragrostis_curvula_Mo  VDVSAGTLVWLRRPNGTWWPSIVISPLDVPDGCPAPPRCAVPIMLLGRRDGFVDWCNLER
A0A5J9VDT3|A0A5J9VDT3_9POAL_Eragrostis_curvula_Mo  ADAEVGALVWVRRRNGSWWPGRILGMDELPDNCVIPPRSGTPIKLLGRPDGSIDWYNLEK
A0A5J9UYK7|A0A5J9UYK7_9POAL_Eragrostis_curvula_Mo  GDTSPGTIVWVRRRNGSWWPGRILGQDELPPSQIMSPRSGTPVKLLGREDASTDWYNLEK
A0A5J9VDJ3|A0A5J9VDJ3_9POAL_Eragrostis_curvula_Mo  ......EALVWVIRRNGSWWPGRILGTDELPENGVLP.RPGTPIKLLGSSDGTIDWYNIED
A0A4D6LTR3|A0A4D6LTR3_VIGUN_Vigna_unguiculata_Eud  VDCDVGSIVWVRRRNGSWWPGQILGSDHLSASHLTSPRSGTPVKLLGREDASVDWYNLEK
A0A4D6LK14|A0A4D6LK14_VIGUN_Vigna_unguiculata_Eud  MDCGVGSIVWVRRRNGSWWPGQILGPDDLSASHLTSPRSGTPVKLLGREDASVDWYNLEK
A0A4D6MNF0|A0A4D6MNF0_VIGUN_Vigna_unguiculata_Eud  IDASVGGLVWVRRRNGSWWPGRILGLHELSESCLVSPRSGTPVKLLGREDASVDWYNLEK
A0A6P6VK84|A0A6P6VK84_COFAR_Coffea_arabica_Eudico  ADCSIGSIVWVRRRNGSWWPGKILGPDDLSASHVMSPRSGTPVKLLGREDASVDWYNLEK
A0A6P6XF68|A0A6P6XF68_COFAR_Coffea_arabica_Eudico  IDASVGGLVWVRRRNGSWWPGRILGPEELPESCLVSPKSGTPVKLLGREDASVDWYNLEK
A0A803M2E7|A0A803M2E7_CHEQI_Chenopodium_quinoa_Eu  LDSTVGALVWVQRRNGCWWPGQIMGLDELPESGSSSPRSGTPIRLLGRDDSSVEWYDLET
A0A803L223|A0A803L223_CHEQI_Chenopodium_quinoa_Eu  RDGGAGSIVWVRRRNGSWWPGKILGPDELSASHLMSPRSGTPVKLLGREDASVDWYNLEK
A0A803MS46|A0A803MS46_CHEQI_Chenopodium_quinoa_Eu  LDTTVGALVWVQRRNGCWWPGQIMGLDELPESGSGSPRSGTPIRLLGRDDSSVEWYDLET
A0A836XPT1|A0A836XPT1_9ROSI_Salix_suchowensis_Eud  ADGNKCPIVWVRRRNGSWWPGQIMEADELAVHNLTSPRTGTPVKLLGRDDASVDWYNLEK
A0A837AV04|A0A837AV04_9ROSI_Salix_suchowensis_Eud  AAGEIGPIVWVRRRNGSWWPGHIMEADELAEYNLTSPRTGTPVKLLGRDDASVDWYNLEK
A0A836YCV2|A0A836YCV2_9ROSI_Salix_suchowensis_Eud  IDASVGALVWVRRRNGSWWPGRIVGLDEVSEGSLVSPRSGTPVKLLGREDASVDWYNLEK
A0A836DE55|A0A836DE55_9ROSI_Salix_suchowensis_Eud  ADGNKCPIVWVRRRNGSWWPGQIMEADELAVHNLTSPRTGTPVKLLGRDDASVDWYNLEK
A0A836DJT9|A0A836DJT9_9ROSI_Salix_suchowensis_Eud  IDASVGALVWVRRRNGSWWPGRIVGLEEISEGSLVSPRSGTPVKLLGREDASVDWYNLEK
```

```
                                                      70        80        90       100
AT1G51745.1_Arabidopsis_thaliana_Eudicot              SKTVKAFRCGEYDTCIEKAKAS..SKKRSGKCTLREDAINNALKIENE
AT3G03140.1_Arabidopsis_thaliana_Eudicot              SKRVKPFRCGDFDECIERVESSQAIIKKREKYARREDAILHALELEKE
AT3G21295.1_Arabidopsis_thaliana_Eudicot              SKRVKAFRCGEYDACIATAKATASTGKKAVKYARREDAIAHALEIENA
AT3G21295.2_Arabidopsis_thaliana_Eudicot              SKRVKAFRCGEYDACIATAKATASTGKKAVKYARREDAIAHALEIENA
AT3G21295.3_Arabidopsis_thaliana_Eudicot              SKRVKAFRCGEYDACIATAKATASTGKKAVKYARREDAIAHALEIENA
KAH7296303.1_Ceratopteris_richardii_ferns_Horseta     SRRVKAFRCGEFDECIERAKSLASLTKKREKYARREDAILHALELEKK
KAH7296310.1_Ceratopteris_richardii_ferns_Horseta     SRRVKAFRCGEFDECIERAKSLASLTKKREKYARREDAILHALELEKK
KAH7435469.1_Ceratopteris_richardii_ferns_Horseta     SRRIKAFRCREFDECIERARSFGCRSKNREKYARRDDAILHALELEKS
KAH7444823.1_Ceratopteris_richardii_ferns_Horseta     SRRVKAFRCGDFDECIERAKSLASLTKKREKYARREDAILHALELEKR
KAH7444826.1_Ceratopteris_richardii_ferns_Horseta     SRRVKAFRCGDFDECIERAKSLASLTKKREKYARREDAILHALELEKR
XP_024515257.1_Selaginella_moellendorffii_spike_m     SKRVKPFRCGDFEECIEKAKAVAGPTKKREKYARREDAILHALELERQ
XP_024515258.1_Selaginella_moellendorffii_spike_m     SKRVKPFRCGDFEECIEKAKAVAGPTKKREKYARREDAILHALELERQ
XP_024535029.1_Selaginella_moellendorffii_spike_m     SKRVKPFRCGDFEECIEKAKAVAGPTKKREKYARREDAILHALELERQ
XP_024535030.1_Selaginella_moellendorffii_spike_m     SKRVKPFRCGDFEECIEKAKAVAGPTKKREKYARREDAILHALELERQ
XP_024540635.1_Selaginella_moellendorffii_spike_m     SKRVKPFRCGEFDDCIEKAQAMAGQSKRREKYARREDAIVDAIELEKE
XP_024541679.1_Selaginella_moellendorffii_spike_m     SKRVKPFRCGEFDDCIEKAQAMAGQSKRREKYARREDAIVDAIELEKE
XP_021890214.1_Carica_papaya_Eudicot                  SKRVKAFRCGEYDDCIEKAKASAASSKKAVKYARREDAILHALEIESA
XP_021890215.1_Carica_papaya_Eudicot                  SKRVKAFRCGEYDDCIEKAKASAASSKKAVKYARREDAILHALEIESA
XP_021890216.1_Carica_papaya_Eudicot                  SKRVKAFRCGEYDDCIEKAKASAASSKKAVKYARREDAILHALEIESA
XP_021890217.1_Carica_papaya_Eudicot                  SKRVKAFRCGEYDDCIEKAKASAASSKKAVKYARREDAILHALEIESA
XP_021900190.1_Carica_papaya_Eudicot                  SKRVKAFRCGEFDDCIERAESSQGPIKKREKYARREDAILHALELEKE
XP_021900191.1_Carica_papaya_Eudicot                  SKRVKAFRCGEFDDCIERAESSQGPIKKREKYARREDAILHALELEKE
XP_004297615.1_Fragaria_vesca_Eudicot                 SKRVKPFRCGEYDECIAKAKAAAAPNKKAAKYARREDAIIHALEIENE
XP_004307887.1_Fragaria_vesca_Eudicot                 SKRVKPFRCGDFDDCIHKAESAQGPPKKREKYARREDAILHALELERE
XP_011463342.1_Fragaria_vesca_Eudicot                 SKRVKPFRCGEYDECIAKAKAAAAPNKKAAKYARREDAIIHALEIENE
XP_031483191.1_Nymphaea_colorata_Eudicot              SKRVKAFRCGEFDDCIRKAEAALGPPRKREKYARREDAILHALELEKQ
XP_031502312.1_Nymphaea_colorata_Eudicot              SKRVKAFRCGEFDDCIRKAESSQGPPKKREKYARREDAILHALELEKL
XP_043631659.1_Erigeron_canadensis_Eudicot            SKRVKPFRCGEFDDCIERAEASQGPPKKREKYARREDAILHALHLEKQ
XP_043637694.1_Erigeron_canadensis_Eudicot            SKRVKAFRCGEFDECIEKAKIIASSCKKTVKYARREDAIIHALKLEGS
XP_042949184.1_Carya_illinoinensis_Eudicot           SKRVKAFRCGEYDECIEKAKASAACNKKAVKYARREDAILHALEIESV
XP_042949194.1_Carya_illinoinensis_Eudicot           SKRVKAFRCGEYDECIEKAKASAACNKKAVKYARREDAILHALEIESV
XP_042949203.1_Carya_illinoinensis_Eudicot           SKRVKAFRCGEYDECIEKAKASAACNKKAVKYARREDAILHALEIESV
XP_042949210.1_Carya_illinoinensis_Eudicot           SKRVKAFRCGEYDECIEKAKASAACNKKAVKYARREDAILHALEIESV
XP_042949220.1_Carya_illinoinensis_Eudicot           SKRVKAFRCGEYDECIEKAKASAACNKKAVKYARREDAILHALEIESV
XP_042967952.1_Carya_illinoinensis_Eudicot           SKRVKAFRCGEYDECIEKAKVSAASNKKAVKYARREDAIIHALEIEST
XP_042979081.1_Carya_illinoinensis_Eudicot           SKRVKAFRCGEFDDCIEKAESSQGPIKKREKYARREDAILHALELEKQ
XP_042983017.1_Carya_illinoinensis_Eudicot           SKRVKAFRCGEFDECIEKAESSQGPIKKREKYARREDAILHALELEKQ
XP_042983018.1_Carya_illinoinensis_Eudicot           SKRVKAFRCGEFDECIEKAESSQGPIKKREKYARREDAILHALELEKQ
XP_042983019.1_Carya_illinoinensis_Eudicot           SKRVKAFRCGEFDECIEKAESSQGPIKKREKYARREDAILHALELEKQ
XP_042983020.1_Carya_illinoinensis_Eudicot           SKRVKAFRCGEFDECIEKAESSQGPIKKREKYARREDAILHALELEKQ
XP_042983022.1_Carya_illinoinensis_Eudicot           SKRVKAFRCGEFDECIEKAESSQGPIKKREKYARREDAILHALELEKQ
XP_042983023.1_Carya_illinoinensis_Eudicot           SKRVKAFRCGEFDECIEKAESSQGPIKKREKYARREDAILHALELEKQ
Thupl.29378772s0023.1.p_Thuja_plicata_Pinopsida       SKRVKAFRCGEYDDCIERAKAYANPAKKVVKYARREDAILHALELEKK
Thupl.29378772s0023.7.p_Thuja_plicata_Pinopsida       SKRVKAFRCGEYDDCIERAKAYANPAKKVVKYARREDAILHALELEKK
Thupl.29378772s0023.3.p_Thuja_plicata_Pinopsida       SKRVKAFRCGEYDDCIERAKAYANPAKKVVKYARREDAILHALELEKK
Thupl.29378772s0023.6.p_Thuja_plicata_Pinopsida       SKRVKAFRCGEYDDCIERAKAYANPAKKVVKYARREDAILHALELEKK
Thupl.29378772s0023.4.p_Thuja_plicata_Pinopsida       SKRVKAFRCGEYDDCIERAKAYANPAKKVVKYARREDAILHALELEKK
Thupl.29378772s0023.5.p_Thuja_plicata_Pinopsida       SKRVKAFRCGEYDDCIERAKAYANPAKKVVKYARREDAILHALELEKK
Thupl.29378772s0023.2.p_Thuja_plicata_Pinopsida       SKRVKAFRCGEYDDCIERAKAYANPAKKVVKYARREDAILHALELEKK
Thupl.29378072s0023.1.p_Thuja_plicata_Pinopsida       SKRVKAFRCGEFNDCIMKAESCAGPTKKREKYARREDAILHALELEKQ
scaffold_ABIJ_2009220_Selaginella_lepidophylla_sp     SKRVKPFRCGDFEECIEKAKAVAGPTKKREKYARREDAILHALELERQ
scaffold_CQPW_2008166_Anemia_tomentosa_ferns_Hors     SRRIKAFRCGEFDDCIAKASAFLE..KTRNKNARREDAIVQALEYEKK
scaffold_CQPW_2008167_Anemia_tomentosa_ferns_Hors     SRRIKAFRCGEFDDCIAKASAFLE..KTRNKNARREDAIVQALEYEKK
scaffold_CQPW_2008862_Anemia_tomentosa_ferns_Hors     SRRIKAFRCGEFDGCIEKASSVEVLPKTRGKNARREDAIFQALELERK
scaffold_CQPW_2020670_Anemia_tomentosa_ferns_Hors     SRRVKAFRCGEFDECIEKARGLASLTKKREKYARREDAILHALELERK
scaffold_DFHO_2051597_Danaea_nodosa_ferns_Horseta     SKRVKPFRCGEFDDCIERAKSVGGPSKKREKYARREDAILHALEL...
scaffold_EEAQ_2011348_Sceptridium_dissectum_ferns     SKRVKAFRCGEFDDCIERAKTFAGPTKKREKYARREDAILHALELEQQ
RWR72543.1_Cinnamomum_micranthum_Magnoliopsida        SKRVKAFRCGDYDEYIEKAKASRGSNKKAGKYASRKDAILRALELESA
RWR81067.1_Cinnamomum_micranthum_Magnoliopsida        SKRVKAFRCGEFDDCIRRAESSQGPPKKREKYARREDAIIHALELENH
RWR86839.1_Cinnamomum_micranthum_Magnoliopsida        SKRVKAFRCGEFNDCIERAESSQGPIRKREKYARREDAILHALELEKE
RWR87272.1_Cinnamomum_micranthum_Magnoliopsida        SKRVKAFRCGEFDDCIERAESAQGPIKKREKYARREDAILHALELERQ
KAG1362729.1_Cocos_nucifera_Monocot                   SKRVKAFRCGEFDACIERAEAAQGPIKKREKYARREDAILHALELEKK
KAG1362730.1_Cocos_nucifera_Monocot                   SKRVKAFRCGEFDACIERAEAAQGPIKKREKYARREDAILHALELEKK
KAF3321551.1_Carex_littledalei_Monocot                SKRVKAFRCGEFDACIERAEAAQGPIKKREKYARREDAILHALELERK
KAF3324589.1_Carex_littledalei_Monocot                SRRVKPFRCGEYDECIEKAKKHGSKNTRKGKYVRREDAILHALEIERK
KAF8671051.1_Digitaria_exilis_Monocot                 SKRVKSFRCGEYDECIEKAKALARHQKRTGRYVRREDAIIHALEIERS
KAF8696638.1_Digitaria_exilis_Monocot                 SKRVKSFRCGEYDECIEKAKALARHQKRTGRYVRREDAIIHALEIERS
KAF8724822.1_Digitaria_exilis_Monocot                 CKRVKPFRCGEFEQRITNALALSATNQTAGRNARMEDAILQALDIERE
KAF8729756.1_Digitaria_exilis_Monocot                 SKRVKAFRCGEFDACIEKAEATQGLVKKREKYARREDAILHALELERK
KAF8750219.1_Digitaria_exilis_Monocot                 SKRVKAFRCGEFDACIEKAEATQGLVKKREKYARREDAILHALELERK
KAG9454374.1_Aristolochia_fimbriata_Magnoliopsida     SKRVKAFRCGEFDDCIERAESSQGPIKKREKYARREDAILHALELERQ
KAG9455570.1_Aristolochia_fimbriata_Magnoliopsida     SSYVKAFRCRDFDGCIEKAKASASCTKTSFKYGCRDDAILHALKIENV
GJM89924.1_Eleusine_coracana_Monocot                  CKRVKPFRCGEFEQRITKAQALDARLGNKRRYARIEDAVLQALEIERA
GJM94157.1_Eleusine_coracana_Monocot                  SKRVKSFRCGEYDECIEKAKALARQQKRTGRYVRREDAIMHALEIERS
GJM94158.1_Eleusine_coracana_Monocot                  SEHVKPFRCREFDMCIENAKVQRREACNKGKYVSINDAIMHALEIEMS
GJN18634.1_Eleusine_coracana_Monocot                  ................................................
URD96891.1_Musa_troglodytarum_Monocot                 STRVKAFRCGEFDECIQKAMASSVSNKFSGKYIHREDAILHALEIEKS
URE09140.1_Musa_troglodytarum_Monocot                 SKRVKAFRCGEFDACIEKAEASLGPIKKREKYARREDAILHALELERK
URE09250.1_Musa_troglodytarum_Monocot                 SKRVKAFRCGEFDACIEKAEASLGPIKKREKYARREDAILHALELERK
URE21899.1_Musa_troglodytarum_Monocot                 STRVKAFRCGEFDECIQKAMAFAISKKSSGKYIRREDAILHALEIEKA
URE21900.1_Musa_troglodytarum_Monocot                 STRVKAFRCGEFDECIQKAMAFAISKKSSGKYIRREDAILHALEIEKA
URE21901.1_Musa_troglodytarum_Monocot                 STRVKAFRCGEFDECIQKAMAFAISKKSSGKYIRREDAILHALEIEKA
URE21903.1_Musa_troglodytarum_Monocot                 STRVKAFRCGEFDECIQKAMAFAISKKSSGKYIRREDAILHALEIEKA
XP_010911968.1_Elaeis_guineensis_Monocot              SKRVKAFRCGEYDECIERAKASAVSKKKPGKYVRREDAILHALEIENA
XP_010911976.1_Elaeis_guineensis_Monocot              SKRVKAFRCGEYDECIERAKASAVSKKKPGKYVRREDAILHALEIENA
XP_010918108.1_Elaeis_guineensis_Monocot              SKRVKAFRCGDFDACIERAEAAQGPIKKREKYARREDAILHALELEKK
XP_010924938.1_Elaeis_guineensis_Monocot              SKRVKPFRCGEYDECIERARASAVSKKKPGKYVRREDAILHALELEKA
XP_010924939.1_Elaeis_guineensis_Monocot              SKRVKPFRCGEYDECIERARASAVSKKKPGKYVRREDAILHALELEKA
XP_010943463.1_Elaeis_guineensis_Monocot              SKRVKAFRCGEFDACIERAEAAQGPIKKREKYARREDAILHALELEKK
XP_019707050.1_Elaeis_guineensis_Monocot              SKRVKPFRCGEYDECIERARASAVSKKKPGKYVRREDAILHALELEKA
XP_042377257.1_Zingiber_officinale_Monocot            SARIKAFRCGEFDECIEKVMISGVSKKRSGKYIRREDAIRHALEIEKP
XP_042383299.1_Zingiber_officinale_Monocot            SARIKAFRCGEFDECIEKVMISGVSKKRSGKYIRREDAIRHALEIEKP
XP_042383893.1_Zingiber_officinale_Monocot            SKRVKEFRCGEFDACIEKAEASLGPIKKREKYARREDAILHALELERK
XP_042383902.1_Zingiber_officinale_Monocot            SKRVKEFRCGEFDACIEKAEASLGPIKKREKYARREDAILHALELERK
XP_042387214.1_Zingiber_officinale_Monocot            SKRVKAFRCGEFDACIEKAEASLGPIKKREKYARREDAILHALELERK
XP_042387215.1_Zingiber_officinale_Monocot            SKRVKAFRCGEFDACIEKAEASLGPIKKREKYARREDAILHALELERK
XP_042392597.1_Zingiber_officinale_Monocot            SKRVKAFRCGEFDACIEKAEASLGPIKKREKYARREDAILHALELERK
XP_042392598.1_Zingiber_officinale_Monocot            SKRVKAFRCGEFDACIEKAEASLGPIKKREKYARREDAILHALELERK
XP_042424141.1_Zingiber_officinale_Monocot            SKRVKSFRCGEFDACIEKAEASLGPIKKREKYARREDAILHALELERK
XP_042424142.1_Zingiber_officinale_Monocot            SKRVKSFRCGEFDACIEKAEASLGPIKKREKYARREDAILHALELERK
XP_042424143.1_Zingiber_officinale_Monocot            SKRVKSFRCGEFDACIEKAEASLGPIKKREKYARREDAILHALELERK
XP_042427101.1_Zingiber_officinale_Monocot            SKRVKSFRCGEFDACIEKAEASLGPIKKREKYARREDAILHALELERK
XP_042427102.1_Zingiber_officinale_Monocot            SKRVKSFRCGEFDACIEKAEASLGPIKKREKYARREDAILHALELERK
```

```
XP_042427103.1_Zingiber_officinale_Monocot          SKRVKSFRCGEFDACIEKAEASLGPIKKREKYARREDAILHALELERK
XP_042450147.1_Zingiber_officinale_Monocot          SKRVKEFRCGEFDACIEKAEASLGPIKKREKYARREDAILHALELERK
XP_042455834.1_Zingiber_officinale_Monocot          SKRVKAFRCGEFDGCIEKAESSLGPVKKREKYARREDAILHALELERK
XP_042455835.1_Zingiber_officinale_Monocot          SKRVKAFRCGEFDGCIEKAESSLGPVKKREKYARREDAILHALELERK
XP_042461069.1_Zingiber_officinale_Monocot          SKRVKAFRCGEFDGCIEKAESSLGPVKKREKYARREDAILHALELERK
XP_042461070.1_Zingiber_officinale_Monocot          SKRVKAFRCGEFDGCIEKAESSLGPVKKREKYARREDAILHALELERK
XP_047063722.1_Lolium_rigidum_Monocot               SKRVKEFRCGEFDACIEKAEASLGPIKKREKYARREDAILHALELERK
XP_047079877.1_Lolium_rigidum_Monocot               SKRVKAFRCGEFDACIERAEATQGLAKKREKYARREDAILHALELERK
XP_047092051.1_Lolium_rigidum_Monocot               SRRVKAFRCGEYEECIEKAKLLARQQKRTGKYVRREDAIMHALEIERS
XP_044951029.1_Hordeum_vulgare_Monocot              SKRVKAFRCGEFDACIERAEATQGVSKKREKYARREDAILHALELERK
XP_044963579.1_Hordeum_vulgare_Monocot              SKRVKPFRCGELDDLIRRTEEQTARRRAGSRNARKEDAVLQALDIERA
XP_044967807.1_Hordeum_vulgare_Monocot              SKRVKEFRCGEFDACIEKAMSCQGPSKRREKYARREDAILHALELERK
XP_044976129.1_Hordeum_vulgare_Monocot              SKRVKAFRCGEYEECIEKAKILARQQKRTGKYVRREDAIMHALEIERS
scaffold_GKAG_2003823_Huperzia_lucidula_clubmoss    SKRVKAFRCGEFDECIENAKALGGSGRRREKYARREDAILDAIELEKK
scaffold_KAWQ_2005490_Stangeria_eriopus_Cycadopsi   SKRVKAFRCGEFDDCIVRAESCAGPAKKREKYARREDAILHALELEKQ
scaffold_KIIX_2013117_Pilularia_globulifera_ferns   SRRVKPFRCGEFDECIEKAKSLASLSKKREKYARREDAILHALELEKK
scaffold_KIIX_2013283_Pilularia_globulifera_ferns   SRRVKAFRCGDFEECIERAKALASLTKKREKYARREDAILHALELERK
scaffold_PKOX_2013427_Isoetes_tegetiformans_quill   SKRVKPFRCGEFDECIEKAKST.GLNRRREKYARREDAILDAIELERQ
scaffold_PKOX_2101345_Isoetes_tegetiformans_quill   SKRVKPFRCGDFDESIEKAQALAGPAKKREKYARREDAILHALELEKQ
scaffold_PNZO_2012682_Culcita_macrocarpa_ferns_Ho   SRRVKSFRCGEFDECIERARSLASLTKKREKYARREDAILHALELEKK
scaffold_TOXE_2013932_Welwitschia_mirabilis_Gneto   SKRVKAFRCGEFDDCISKAESFAGPSKKREKYARREDAILHALELERQ
scaffold_UWOD_2005868_Plagiogyria_japonica_ferns_   SRRVKSFRCGEFDECIERARSLASLTKKREKYARREDAILHALELEKK
scaffold_UWOD_2005869_Plagiogyria_japonica_ferns_   SRRVKSFRCGEFDECIERARSLASLTKKREKYARREDAILHALELEKK
scaffold_UWOD_2014926_Plagiogyria_japonica_ferns_   SRRIKAFRCGEFDACIERARSLTSLLKKREKYARREDAILHALELEKK
scaffold_VIBO_2011581_Plenasium_javanicuma_ferns_   SRRVKAFRCGEFDDCIERAKALASLKKKREKYARREDAILHALELEQK
scaffold_VSRH_2006872_Abies_lasiocarpa_Pinopsida    SKRVKAFRCGEFDDCILKAESCAGPTKKREKYARREDAILHALELEKL
scaffold_VSRH_2012377_Abies_lasiocarpa_Pinopsida    SKRVKAFRCGEYDDCIEKAKAYANPAKKVVKYARREDAILHALELEKK
scaffold_XNXF_2063615_Dendrolycopodium_obscurum_c   SKRVKAFRCGEFNECIETAKALGGSGKRREKY..............
scaffold_YFZK_2002000_Sciadopitys_verticillata_Pi   SKRVKAFRCGEFDDCISKAESCAGPTKKREKYARREDAILHALELEKQ
scaffold_YFZK_2004458_Sciadopitys_verticillata_Pi   SKRVKAFRCGEYDDCIERAKASANPAKKVVKY..............
scaffold_ZYCD_2042766_Selaginella_acanthonota_spi   SKRVKPFRCGDFEECIEKAKAVAGPTKKREKYARREDAILHALELEKQ
V7AHI2|V7AHI2_PHAVU_Phaseolus_vulgaris_Eudicot      SKRVKAFRCGEFDDCIEKAESAQGPLKKREKYARREDAILHALELEKQ
V7AQI6|V7AQI6_PHAVU_Phaseolus_vulgaris_Eudicot      SKRVKAFRCGEYDECIEKAKASAASNKKAVKYARREDAILHALELESA
V7CFU1|V7CFU1_PHAVU_Phaseolus_vulgaris_Eudicot      SKRVKTFRCGEFDGCIERAESAQGPLKKREKYARREDAILHALELERQ
Q0DYD8|Q0DYD8_ORYSJ_Oryza_sativa_Monocot            SKRVKAFRCGEFDACIEKAEATEGSVKKREKYARREDAILHALELEKK
Q7X7G8|Q7X7G8_ORYSJ_Oryza_sativa_Monocot            SKRVKAFRCGEFDACIEKALTSRGPVKRREKYARREDAILHALELERK
Q0JLX9|Q0JLX9_ORYSJ_Oryza_sativa_Monocot            SKRVKPFRCGEYEECIEKAKAQARQHKRAGKYVRREDAIMHALELERA
C5XN34|C5XN34_SORBI_Sorghum_bicolor_Monocot         SKRVKSFRCGEYDECIEKAKVLARQQKRTGRYVRREDAIMHALEIERS
A0A1B6PAF8|A0A1B6PAF8_SORBI_Sorghum_bicolor_Monoc   CKRVKPFRCGEFEQRITNALTLAAGNRSTGRYARMEDAILQALDIEKE
C5Y0E5|C5Y0E5_SORBI_Sorghum_bicolor_Monocot         SKRVKAFRCGEFDACIEKAEATQGVVKKREKYARREDAILHALELERK
A0A3Q7H2C5|A0A3Q7H2C5_SOLLC_Solanum_lycopersicum_   SKRVKAFRCGEFDDFIQRAEASLGPPKKREKYARREDAILHALELERQ
A0A3Q7FMI9|A0A3Q7FMI9_SOLLC_Solanum_lycopersicum_   SKRVKAFRCGEFDDCIKRAEASQGPPKKREKYARREDAILHALELERL
A0A3Q7FVX7|A0A3Q7FVX7_SOLLC_Solanum_lycopersicum_   SKRVKAFRCGEYDECIEKAKASAASSKKAVKYARREDAIIHALEIESA
B9I743|B9I743_POPTR_Populus_trichocarpa_Eudicot     SKRVKAFRCAEFSDCIKRAESALGPIKKREKYARREDAILHALELEKR
A0A2K2C0C5|A0A2K2C0C5_POPTR_Populus_trichocarpa_E   SKRVKAFRCGEYDECIEKAKASAAGNKRVVKYARREDAILHALEIENA
A0A2K2B1H6|A0A2K2B1H6_POPTR_Populus_trichocarpa_E   SKRVKAFRCGEYDECIEKAKTSAAGNKRAVKYARREDAILHALEIENA
A0A2K1WPP7|A0A2K1WPP7_POPTR_Populus_trichocarpa_E   SKRVKAFRCAEFSDCIERAESALGPIKKREKYARREDAILHALELEKQ
A0A804QV00|A0A804QV00_MAIZE_Zea_mays_Monocot        SKRVKSFRCGEYDECIEKAKALVRRQKKTGRYVRREDAIMHALEIERS
A0A804QP35|A0A804QP35_MAIZE_Zea_mays_Monocot        CKRVKPFRCGELEQRIANALTLAASDRSTGRYARMEDAILQALDIEKE
A0A804QS22|A0A804QS22_MAIZE_Zea_mays_Monocot        SKRVKSFRCGEYDECIEKAKALVRRQKKTGRYVRREDAIMHALEIERS
A0A804QSQ4|A0A804QSQ4_MAIZE_Zea_mays_Monocot        SKRVKSFRCGEYDECIEKAKALVRRQKKTGRYVRREDAIMHALEIERS
A0A804QQ17|A0A804QQ17_MAIZE_Zea_mays_Monocot        SKRVKSFRCGEYDECIEKAKALVRRQKKTGRYVRREDAIMHALEIERS
A0A804NUY1|A0A804NUY1_MAIZE_Zea_mays_Monocot        SKRVKAFRCGEFDACIEKAEATQGLVKKREKYARREDAILHALELERK
A0A804NI78|A0A804NI78_MAIZE_Zea_mays_Monocot        SKRVKSFRCGEYDECIEKAKALARQQKRTGRYVRREDAILHALEIERS
A0A1D6MDX6|A0A1D6MDX6_MAIZE_Zea_mays_Monocot        CKRVKPFRCGEFEQRITNALTLAAGNRSSGRYARMEGAILQALDIEKD
K7UY21|K7UY21_MAIZE_Zea_mays_Monocot                CKRVKPFRCGELEQRIANALTLAASDRSTGRYARMEDAILQALDIEKE
A0A1D6NMK7|A0A1D6NMK7_MAIZE_Zea_mays_Monocot        SKRVKSFRCGEYDECIEKAKALVRRQQKRTGRYVRREDAILHALEIERS
A0A1D6HH07|A0A1D6HH07_MAIZE_Zea_mays_Monocot        SKRVKAFRCGEFDACIEKAEATQGLVKKREKYARREDAVLHALELERK
A0A1D6HH08|A0A1D6HH08_MAIZE_Zea_mays_Monocot        SKRVKAFRCGEFDACIEKAEATQGLVKKREKYARREDAVLHALELERK
A0A1D6DQP9|A0A1D6DQP9_MAIZE_Zea_mays_Monocot        SKRVKAFRCGEFDACIEKAEATQGLVKKREKYARREDAILHALELERK
A0A2K2CIB3|A0A2K2CIB3_BRADI_Brachypodium_distachy   SKRVKEFRCGDFDACIEKAMSCQGPVKKREKYARREDAILHALKLESK
I1HNL6|I1HNL6_BRADI_Brachypodium_distachyon_Monoc   SKRVKAFRCGEYEECIEKAKVLAHQHKRAGKYVRREDAILHALEIERS
I1HFR1|I1HFR1_BRADI_Brachypodium_distachyon_Monoc   SKRVKPFRCGDLDDCIRKAQQQQARRRGSARYARKEDAVLQALQIERA
I1ICW8|I1ICW8_BRADI_Brachypodium_distachyon_Monoc   SKRVKAFRCGEFDACIERAEATQGLAKKREKYARREDAILHALELERK
I1LIJ1|I1LIJ1_SOYBN_Glycine_max_Eudicot             SKRVKAFRCGEFDDCIEKAEAAQGPLKKREKYARREDAILHALELEKQ
K7LMH1|K7LMH1_SOYBN_Glycine_max_Eudicot             SKRVKAFRCGEFDGCIERAESAQGPLKKREKYARREDAILHALELEKQ
K7LSJ6|K7LSJ6_SOYBN_Glycine_max_Eudicot             SKRVKAFRCGEFDDCIEKAETAQGPLKKREKYARREDAILHALELEKQ
A0A0R0LFH3|A0A0R0LFH3_SOYBN_Glycine_max_Eudicot     SKRVKAFRCGEFDGCIERAESAQGPLKKREKYARREDAILHALDLERQ
A0A0R0IPK3|A0A0R0IPK3_SOYBN_Glycine_max_Eudicot     SKRVKAFRCGEYNECIEKAKASAASSKKAVKYARREDAILHALELESA
I1JAI6|I1JAI6_SOYBN_Glycine_max_Eudicot             SKRVKAFRCGEFDGCIERAESAQGPLKKREKYARREDAILHALDLERQ
I1KGF6|I1KGF6_SOYBN_Glycine_max_Eudicot             SKRVKAFRCGEYDECIEKAKASAASSKKAVKYARREDAILHALELESA
K7LMH0|K7LMH0_SOYBN_Glycine_max_Eudicot             SKRVKAFRCGEFDGCIERAESAQGPLKKREKYARREDAILHALELEKQ
tr|F6HEG8|F6HEG8_VITVI_Vitis_vinifera_Eudicot       SKRVKAFRCGEFDDCIERAESSQGPLKKREKYARREDAILHALELEKQ
tr|F6HY39|F6HY39_VITVI_Vitis_vinifera_Eudicot       SKRVKAFRCGEYDECIEKAKASAASNKKAVKYARREDAILHALEIENA
M4ET84|M4ET84_BRARP_Brassica_rapa_Eudicot           SKRVKPFRCGDFDDCIEKVENSQGTIRKREKYARREDAILHALELEKE
M4ENQ0|M4ENQ0_BRARP_Brassica_rapa_Eudicot           SKSVKAFRCGDYDSHIDKAMAVAANKKTAKLTRRENAINIALQIENA
M4DXX3|M4DXX3_BRARP_Brassica_rapa_Eudicot           SKRVKPFRCGDFDDCIEKVETSQGTVRKREKYARREDAILHALELEKE
M4ER48|M4ER48_BRARP_Brassica_rapa_Eudicot           SKRVKAFRCGEYDACIETAKATATASKKPVKYARREDAIVHALEIESA
U5CR03_AMBTC_Amborella_trichopoda_Eudicot           SKRVKAFRCGEFDDCIGRAESSQGPAKKREKYARREDAILHALELEKQ
U5CQG7_AMBTC_Amborella_trichopoda_Eudicot           SKRVKAFRCGEYDECIEKAKAYAAPAKKAVKYARREDAILHALELESA
R0I6R0|R0I6R0_9BRAS_Capsella_rubella_Eudicot        SKRVKPFRCGDFDECIERVESSQATIRKREKYARREDAILHALELEKE
R0GPB7|R0GPB7_9BRAS_Capsella_rubella_Eudicot        SKTVKAFRCGEYDSCIEKAKAAAASNKKAAKCTRREDAIKNALEIENE
R0I129|R0I129_9BRAS_Capsella_rubella_Eudicot        SKRVKAFRCGEYDACIETAKATVSTSKKAVKYARREDAIAHALEIESA
tr|V4SLI4|V4SLI4_CITCL_Citrus_sinensis_Eudicot      SKRVKAFRCGEYDECIAKAKASAASGKKAVKYARREDAILHALELESA
tr|V4U2M5|V4U2M5_CITCL_Citrus_sinensis_Eudicot      SKRVKAFRCGEFDDCIERAEASLGPTRKREKYARREDAILHALELEKE
tr|A0A058ZXU8|A0A058ZXU8_EUCGR_Eucalyptus_grandis   SKRVKAFRCGEFDECIERAESSQGPMKKREKYARREDAILHALELEKE
tr|A0A059BTW7|A0A059BTW7_EUCGR_Eucalyptus_grandis   SKRVKAFRCGEYDDCIEKAKAAAALSKKVVKYARREDAILHALELESS
A0A1S2YUZ3|A0A1S2YUZ3_CICAR_Cicer_arietinum_Eudic   SKRVKAFRCGEYDECIEKAKASAALGKKAVKYARREDAILHALELESA
A0A1S3EG88|A0A1S3EG88_CICAR_Cicer_arietinum_Eudic   SKRVKPFRCGEFDGCIERAESAQGPLKKREKYARREDAILHALELERQ
A0A1S2YJA8|A0A1S2YJA8_CICAR_Cicer_arietinum_Eudic   SKRVKAFRCGEFDECIERVESSHEPLKKREKYARREDAVLHALELERQ
A0A251JRZ2|A0A251JRZ2_MANES_Manihot_esculenta_Eud   SKRVKAFRCGEYDECIEKAKANAAGNKKTVKYARREDAILHALEIENA
A0A2C9VI54|A0A2C9VI54_MANES_Manihot_esculenta_Eud   SKRVKAFRCGEFDDCIERAESAQGPIKKREKYARREDAILHALDLEKQ
A0A2C9VA71|A0A2C9VA71_MANES_Manihot_esculenta_Eud   SKRVKAFRCGDFDDCIERAESAQGRIKKREKYARREDAILHALELEKQ
A0A2C9VA68|A0A2C9VA68_MANES_Manihot_esculenta_Eud   SKRVKAFRCGDFDDCIERAESAQGRIKKREKYARREDAILHALELEKQ
A0A1Q3CJL5_CEPFO_Cephalotus_follicularis_Eudicot    SKRVKAFRCGEYEECIAKAKVAAAAGKKVVKYARREDAILHALELESA
A0A1Q3D3W0_CEPFO_Cephalotus_follicularis_Eudicot    SKRVKAFRCGEFDDCIERAESAQGPIKKREKYARREDAIHHALELEKE
A0A1J6IUT4_NICAT_Nicotiana_attenuata_Eudicot        SKRVKEFRCGEFDELIQRAETSLGPPKKREKYARREDAILHALELEKQ
A0A1J6KD13_NICAT_Nicotiana_attenuata_Eudicot        SKRVKAFRCGEYDECIEKAKAAAASSRKAVKYARREDAIIHALEIESA
A0A1U8JDV0|A0A1U8JDV0_GOSHI_Gossypium_hirsutum_Eu   SKRVKAFRCGEYNECIEKAKASAASSKKAVKYARREDAILHALEIESS
A0A1U8MT23|A0A1U8MT23_GOSHI_Gossypium_hirsutum_Eu   SKRVKAFRCGEYNECIEKAKASAASSKKAVKYARREDAILHALEIESS
A0A1U8KH73|A0A1U8KH73_GOSHI_Gossypium_hirsutum_Eu   SKRVKAFRCGEYNECIEKAKASAASSKKAVKYARREDAILHALEIESA
A0A1U8K8A4|A0A1U8K8A4_GOSHI_Gossypium_hirsutum_Eu   SKRVKAFRCGEFDDCIERAESSLGPPKKREKYARREDAILHALELEKE
A0A1U8NQ29|A0A1U8NQ29_GOSHI_Gossypium_hirsutum_Eu   SKRVKAFRCGEFDDCIERAESSQGPPKKREKYARREDAILHALELEKE
```

```
A0A251VD55|A0A251VD55_HELAN_Helianthus_annuus_Eud  SKRVKAFRCGEYDECIEKAKFTANSCKKAAKYARREDAILQALEIENS
A0A251TN31|A0A251TN31_HELAN_Helianthus_annuus_Eud  SKRVKPFRCGEDDG..................................
A0A251UUN3|A0A251UUN3_HELAN_Helianthus_annuus_Eud  SKRVKPFRCGEYDECIKKAKATATSCKKAAKYAHREDAILQALELENS
A0A251SME0|A0A251SME0_HELAN_Helianthus_annuus_Eud  SKRVKPFRCGEFDDCIERAEASQGPPKKREKYARREDAILHALELEKQ
A0A8B7CF00|A0A8B7CF00_PHODC_Phoenix_dactylifera_M  SKRVKAFRCREYDECIERAKASAVSKKKPGKYVRREDAILHALEIEKA
A0A8B7BTD1|A0A8B7BTD1_PHODC_Phoenix_dactylifera_M  SKRVKAFRCGEFDACIERAEAAQGPIKKREKYARREDAILHALELEKK
A0A8B7CIS5|A0A8B7CIS5_PHODC_Phoenix_dactylifera_M  SKRVKAFRCGEFGACIERAEAAQGPIKKREKYARREDAILHALELEKK
A0A2P5AGS9_PARAD_Parasponia_andersonii_Eudicot     SKRVKAFRCGEYDDCIEKAKASAASSKKAVKYARREDAIIHALEIENA
A0A2P5AY67_PARAD_Parasponia_andersonii_Eudicot     SKRVKAFRCGEFDDCIERAESSQGPIKKREKYARREDAILHALELEKQ
A0A2P6SPQ0_ROSCH_Rosa_chinensis_Eudicot            SKRVKPFRCGDFDDCIHKAESAQGPPKKREKYARREDAILHALELERQ
A0A2P6RXT9_ROSCH_Rosa_chinensis_Eudicot            SKRVKAFRCGEYDECIEKAKAAAAPNKKAVKYARREDAIIHALEIENE
A0A103XW46|A0A103XW46_CYNCS_Cynara_cardunculus_Eu  SKRVKAFRCGEYDECIEKAKVSAASCKKAVKYARREDAILHALELESS
A0A103XCS8|A0A103XCS8_CYNCS_Cynara_cardunculus_Eu  SKRVKPFRCGEFDDCIERAEASQGPPKKREKYARREDAILHALELEKQ
A0A2T7FAX4|A0A2T7FAX4_9POAL_Panicum_hallii_Monoco  SKRVKAFRCGEFDACIEKAEATQGLVKKREKYARREDAILHALELERK
A0A2T7DKV0|A0A2T7DKV0_9POAL_Panicum_hallii_Monoco  SKRVKSFRCGEYDECIEKAKALARQQKRTGRYVRREDAIIHALEIERS
A0A2T7E9E4|A0A2T7E9E4_9POAL_Panicum_hallii_Monoco  CKRVKPFRCGEFEERITNALAA..GNKTSGRYARMEDAILQALDIERE
A0A2T7FAX6|A0A2T7FAX6_9POAL_Panicum_hallii_Monoco  SKRVKAFRCGEFDACIEKAEATQGLVKKREKYARREDAILHALELERK
A0A5J9SJ79|A0A5J9SJ79_9POAL_Eragrostis_curvula_Mo  CKRVKPFRCGEFDQRITHAQAIAARVHYKGKYARMEDAVLQALEIERA
A0A5J9VDT3|A0A5J9VDT3_9POAL_Eragrostis_curvula_Mo  SKRVKSFRCGEYDECIEKAKVLARQQKRTGRYVRREDAIMHALEIERS
A0A5J9UYK7|A0A5J9UYK7_9POAL_Eragrostis_curvula_Mo  SKRVKAFRCGEFDACIEKAEATQGLVKKREKYARREDAILHALELERK
A0A5J9VDJ3|A0A5J9VDJ3_9POAL_Eragrostis_curvula_Mo  SKCVKPFRCGEFAECIENAKVRGRIAYKEGKYACRDDAIMHALAIEMS
A0A4D6LTR3|A0A4D6LTR3_VIGUN_Vigna_unguiculata_Eud  SKRVKTFRCGEFDGCIERAESAQGPLKKREKYARREDAILHALELERQ
A0A4D6LK14|A0A4D6LK14_VIGUN_Vigna_unguiculata_Eud  SKRVKAFRCGEFDDCIEKAESAQGPLKKREKYARREDAILHALELEKQ
A0A4D6MNF0|A0A4D6MNF0_VIGUN_Vigna_unguiculata_Eud  SKRVKAFRCGEYDECIEKAKASAASNKKAVKYARREDAILHALELESA
A0A6P6VK84|A0A6P6VK84_COFAR_Coffea_arabica_Eudico  SKRVKAFRCGEFDDCIERAEASQGPPKKREKYARREDAILHALELERQ
A0A6P6XF68|A0A6P6XF68_COFAR_Coffea_arabica_Eudico  SKRVKAFRCGEYDDCIEKAKAAA.SSKKVVKYARREDAILHALELENA
A0A803M2E7|A0A803M2E7_CHEQI_Chenopodium_quinoa_Eu  AKGVKAFRCGEYDTCIEKAKVSAPFLKRSLKYPRREIAIFYALEIEKS
A0A803L223|A0A803L223_CHEQI_Chenopodium_quinoa_Eu  SKRVKAFRCGEFNDCIERAEASVGPAKKREKYARREDAILHALELEKQ
A0A803MS46|A0A803MS46_CHEQI_Chenopodium_quinoa_Eu  AKGVKAFRCGEYDTCIEKAKVSAPFLKRSLKYPRREIAIFYALEIEKS
A0A836XPT1|A0A836XPT1_9ROSI_Salix_suchowensis_Eud  SKRVKAFRCAEFSDCIERAESALGPIKKREKYARREDAILHALELEKQ
A0A837AV04|A0A837AV04_9ROSI_Salix_suchowensis_Eud  SKRVKAFRCAEFSDCIKRAESALGPIKKREKYARREDAILHALELEKQ
A0A836YCV2|A0A836YCV2_9ROSI_Salix_suchowensis_Eud  SKRVKAFRCGEYDECIEKAKTSAAGNKRAVKYARREDAILHALEIENA
A0A836DE55|A0A836DE55_9ROSI_Salix_suchowensis_Eud  SKRVKAFRCAEFSDCIERAESALGPIKKREKYARTLD...........
A0A836DJT9|A0A836DJT9_9ROSI_Salix_suchowensis_Eud  SKRVKAFRCGEYDECIEKAKASAAGNKRVVKYARREDAILHALEIENA
```
